# Gigabase-scale deletion scanning of the human genome

**DOI:** 10.64898/2026.05.29.728882

**Authors:** Jonas Koeppel, Aidan Keith, Samantha Sgrizzi, Peixi Chen, Riza M. Daza, Faaiz Quaisar, Eleftheria Anastasia, Zihao Song, Jay Shendure, Sudarshan Pinglay

## Abstract

What fraction of the human genome is essential for cellular viability? To date, essentiality in human cells has been mapped almost exclusively at the level of individual open reading frames (ORFs). Whether noncoding regions and broader architectural features of the genome are required for human cells to remain viable, and where the boundaries of any such regions lie, remains largely unexplored. Here we introduce Shred-seq, which couples Type I-C CRISPR-Cas3-mediated deletions with phage polymerase-based genotyping to enable large-scale deletion scans of the human genome. Shred-seq leverages thousands of genomically integrated, mapped target sites as launchpads for Cas3 to initiate unidirectional deletions ranging in size from hundreds of base pairs (bp) to hundreds of kb. Breakpoints are directly captured at high resolution by *in vitro* or *in situ* transcription from flanking phage polymerase promoters, enabling bulk or single-cell phenotyping, respectively. In this proof-of-concept, we generate and genotype 36,257 independent deletions originating from 9,604 Cas3 launchpads, individually spanning 100 bp to 500 kb, collectively covering 461 Mb (14% of the human genome), and totaling to 2.55 Gb of deleted sequence (∼10-fold coverage of these regions). Surviving deletions are depleted not only for essential protein-coding genes, but also for active, conserved and mutation-constrained non-coding sequences, directly quantifying purifying selection across both coding and noncoding intervals. Deletion length distributions further enable annotation-agnostic fine mapping of essential region boundaries and establish an empirical lower bound on the fraction of the human genome required for cellular viability. Finally, we demonstrate compatibility with single-cell RNA-seq (scRNA-seq), enabling direct linkage of specific deletions to transcriptional phenotypes. Together, Shred-seq provides a scalable platform for the systematic dissection of genome architecture and noncoding function, as well as for generating training data for predictive and generative models of genome structure-function, analogous to the roles that conventional Perturb-seq screens play for ORFs. By enabling the empirical delimitation of the genomic content required for human cellular viability, Shred-seq may also open a path to the construction of a minimal human genome.

## INTRODUCTION

Which subsequences of the human genome are essential? At the organismal level, essentiality is defined by viability and reproductive success across generations. At the cellular level, essentiality is defined by a cell’s ability to survive and proliferate in a given environment. These constraints presumably overlap but are not equivalent, and despite substantial progress in human genetics and functional genomics, we still do not know what fraction of the human genome is dispensable versus required under either definition.

At the organismal level, human genetics provides an enormous natural experiment in loss-of-function variants, including structural variants (SVs) that remove kilobase- to megabase-scale segments (Sudmant et al. 2015; Ebert et al. 2021). However, most large deletions observed in healthy individuals are heterozygous, with allele frequencies potentially shaped by pleiotropic effects in heterozygotes rather than by selection against complete loss (Judd et al. 2025; Milind et al. 2024). More generally, the variants that are observed are the survivors of multiple filters—developmental viability, fertility, and population history—making it difficult to infer what fraction of the genome could be deleted in principle, as opposed to what is merely tolerated. Comparative genomics reinforces this uncertainty: although evolutionary conservation suggests that only 5-10% of the genome is under evolutionary constraint (Galeota-Sprung et al. 2020; Ponting and Hardison 2011; Leypold and Speicher 2021; Cooper et al. 2005), the persistently high ratio of unconstrained to constrained sequence across vertebrates implies that important aspects of genome function—particularly at larger length scales—may not be captured by base-pair-level conservation alone. Striking counterexamples, such as the extreme genome compaction observed in teleosts such as *Fugu* (Brenner et al. 1993; Aparicio et al. 2002), further underscore how incomplete our understanding remains of which features of genome architecture are truly required for organismal viability.

At the cellular level, essentiality has been mapped more systematically, but almost exclusively at the level of ORFs. Genome-wide CRISPR knockout screens have identified thousands of protein-coding genes required for proliferation and revealed strong context dependence across genetic and environmental backgrounds (Arafeh et al. 2025; Hart et al. 2015). In contrast, far less is known about essentiality outside of ORFs, nor about essentiality at the scale of genomic intervals rather than annotated elements. This limitation is not simply a matter of incomplete annotation: prevailing perturbation modalities are optimized to focally disrupt coding sequences or short regulatory elements, and are therefore poorly suited to test whether intact enhancers, multi-kilobase regulatory landscapes, or higher-order genomic features are required for cellular viability. As a result, even in well-studied cellular systems, we still lack answers to basic questions: how much of the genome is dispensable for cell survival in a defined context, where the true boundaries of essential regions lie, and whether essential noncoding elements remain undiscovered.

By interrogating contiguous genomic intervals with redundancy, deletion scans have the potential to resolve essential regions with greater precision than point mutations or targeted knockouts—whether for identifying noncoding sequences required for cellular viability or for pinpointing causal elements within disease-associated SVs (Gasperini et al. 2017; Su et al. 2000). Contemporary approaches—nested Cre-loxP engineering or paired-guide CRISPR-Cas9 deletions—have been applied at individual loci but do not readily scale genome-wide. They require pre-programmed pairs of target sites, clonal isolation and verification, or rely on indirect inference of deletion boundaries from guide identity (Gasperini et al. 2017; Su et al. 2000; Diao et al. 2017; Zhu et al. 2016). Even in larger-scale screens, functional readouts are typically limited to proliferation, leaving richer or more nuanced phenotypic consequences unexplored.

For genome-wide deletion scanning, an ideal method would generate barcoded deletions with stochastic breakpoints across a wide range of sizes—enabling scalable quantitation of both deletion boundaries and phenotypic consequences without requiring pre-specified target pairs. To this end, we were drawn to the Class 1 CRISPR-Cas3 system as a potential solution. Cas3 is the hallmark nuclease-helicase of Class 1 Type I CRISPR systems, which constitute the majority of CRISPR loci found in nature (Hille et al. 2018; Koonin et al. 2017). In Type I systems, a multi-protein Cascade complex recognizes a DNA target via a crRNA and recruits Cas3. Cas3 translocates along the DNA using its helicase activity, while its nuclease activity leaves DNA breaks in its wake, which result in large and unidirectional deletions ranging from hundreds of bp to hundreds of kb (Cameron et al. 2019; Morisaka et al. 2019; Tan et al. 2022). Initial demonstrations in human cells using Type I-E systems showed that Cas3 could efficiently induce heterogeneous 30-100 kb deletions and knock out genes of interest (Cameron et al. 2019; Morisaka et al. 2019). More recently, Tan et al. adapted the compact Type I-C system from *Neisseria lactamica* for mammalian genome editing, which proved highly efficient at creating large deletions (Tan et al. 2022).

However, despite Cas3’s ability to generate deletions of variable length, its utility for functional genomics remains limited by the inability to genotype engineered variants at scale. Existing approaches rely on PCR amplification and sequencing of anticipated deletion junctions—an approach that fails for deletions of unpredictable size because it is impractical to position primers to capture all possible junctions (Morisaka et al. 2019; Tan et al. 2022; Kosicki et al. 2018). The alternative, clonal isolation followed by whole-genome sequencing, is practical for a handful of engineered lines but cannot scale to the hundreds of thousands of variants required for genome-wide interrogation (Koeppel et al. 2025; Tsai et al. 2023). Moreover, neither strategy readily links SV genotypes to rich functional readouts beyond bulk viability.

We reasoned that genotyping based on *in vitro* or *in situ* transcription (IVT/IST) from dormant phage promoters (*e.g.*, T7, SP6, T3) could overcome this limitation (X. Li et al. 2024; Askary et al. 2020; Pinglay et al. 2025). For example, we recently leveraged phage polymerase-mediated IVT/IST to map the locations of thousands of barcoded integrations of Cas9 or PEmax target sites (X. Li et al. 2024) and to genotype SVs at scale in both bulk and single-cell contexts (Pinglay et al. 2025). Adapting this strategy to Cas3, if deletions were induced downstream of a genomically integrated and mapped phage polymerase promoter, transcripts originating from that promoter would capture the deletion breakpoint, without requiring prior knowledge of its location (**Figure 1A**).

**Figure 1.**
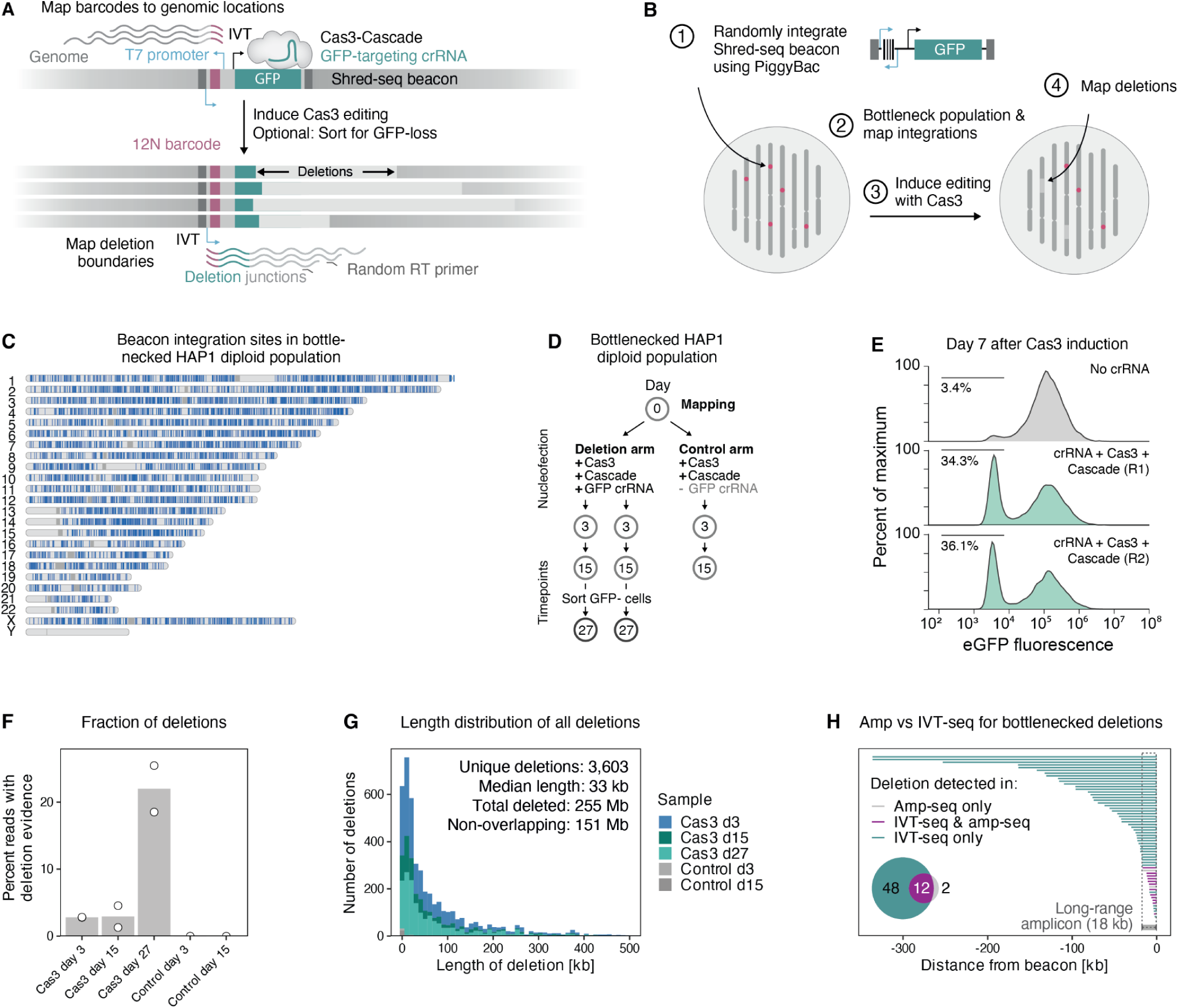
Shred-seq enables the multiplexed generation and characterization of thousands of nested deletions from launchpoints throughout the human genomes. (**A**) Schematic of integrated Shred-seq beacon (v1), which contains a T7 promoter, a random 12N DNA barcode, and an expressed GFP transgene. Cas3 editing is induced with a GFP-targeting crRNA. *In vitro* (IVT) or *in situ* (IST) transcription with T7 polymerase, followed by reverse transcription and sequencing, allows for the mapping of cassette insertion locations and Cas3-induced deletion boundaries. (**B**) Shred-seq workflow. PiggyBac is used to randomly integrate degenerately barcoded beacons, the cell pool is bottlenecked, and the insertion sites associated with each barcode are mapped. Editing is induced with Cas3, and deletions are called. (**C**) Genome-wide distribution of uniquely mapped barcodes in bottlenecked diploid HAP1 cells (rows: chromosomes; ticks: mapped integrations, centromeres darkgray). (**D**) Experimental outline for Shred-seq pilot experiment. (**E**) Distribution of eGFP fluorescence, 7 days after Cas3 induction (x-axis) for no-crRNA negative control and two biological replicates (panels). The percentage of cells with GFP loss is indicated. (F) Barplot of percent of reads containing deletion evidence across time points and samples. Dots represent biological replicates, and bars correspond to mean. (**G**) Histogram of the number of deletions (y-axis) across various lengths (x-axis) of all called deletions across samples (colors). Bars reflect the sum of two biological replicates. (**H**) Deletion coordinates (lines, x-axis) from a clonal beacon integration site colored by detection method. The 18 kb long-range PCR amplicon is indicated by the gray box and dotted lines. The inset shows a size-proportional Venn diagram of deletions detected by IVT-seq, long-range amplicon sequencing, or both.

This insight forms the basis of Shred-seq, which combines Type I-C CRISPR-Cas3 deletion generation with phage polymerase-based SV genotyping to enable highly multiplexed deletion scans in human cells. Specifically, we apply Shred-seq to generate and genotype tens of thousands of deletions, ranging from hundreds of bp to hundreds of kb, originating from thousands of genomically mapped launchpads, and collectively covering ∼461 Mb of the human genome. We characterize how the local epigenome shapes Cas3 deletion initiation and termination and characterize the repair outcomes of Cas3-induced lesions. By tracking deletion frequencies over time in haploid cells, we quantify signatures of purifying selection and show that deletion length distributions can fine-map the boundaries of essential regions. Finally, we demonstrate that Shred-seq is compatible with scRNA-seq, enabling direct linkage of deletion genotypes to transcriptional phenotypes. Together, the methods and results presented here establish a scalable framework for systematic deletion scanning of the entire human genome.

## RESULTS

### Design of Shred-seq

Shred-seq is based on the integration of uniquely barcoded ‘beacons’ into the genome, which serve as directional launchpoints for Cas3-mediated nested deletions. Each Shred-seq beacon is designed to contain: (i) a unique DNA barcode, which allows each beacon to be individually tracked across the experiment; (ii) a genome-orthogonal synthetic target site, from which Cas3 initiates deletions; and (iii) a pair of convergent phage RNA polymerase promoters (T7 or SP6) flanking the barcode, which together enable base-pair resolution mapping of genomic integration sites as well as the high-throughput genotyping of resulting deletions at bulk or single-cell resolution (Pinglay et al. 2025; X. Li et al. 2024). Optionally, beacons can also contain elements that allow for the enrichment of cells that harbor Cas3-induced genomic deletions (**Figure 1A**).

In our initial design, the beacon contained a minimal eGFP expression cassette that serves as both the target site to initiate deletions from using published crRNAs (Tan et al. 2022) and as a reporter for successful deletion events based on loss of fluorescence (**Figure 1A**). Following random or targeted integration of the Shred-seq beacons into the genome, cell populations are sorted and bottlenecked based on eGFP fluorescence (**Figure 1B**). The outward-facing T7 promoter is used to map integration sites via IVT, thereby associating each barcode with its genomic location. Upon introduction of Cas3, Cascade and a crRNA targeting the genome-orthogonal synthetic target site, large unidirectional deletions are initiated from the integrated target sites. The inward-facing T7 promoter then generates IVT transcripts that span both the barcode and the newly created genomic junction, thereby directly capturing the deletion endpoint without requiring whole-genome sequencing.

After deletions are generated, cells can be subjected to *in vitro* selections, with T7 IVT providing direct identification and quantification of deletion abundances over time. This approach also provides natural signal amplification through T7 transcription and allows sequencing resources to be focused exclusively on the informative breakpoint-containing molecules. Moreover, as previously described, T7 transcripts can be captured after IST in fixed cells, allowing Shred-seq genotypes to be linked directly to single-cell transcriptomes via scRNA-seq (Pinglay et al. 2025; X. Li et al. 2024) and enabling measurement of deletion impact on genome-wide gene expression.

Because thousands of barcoded beacons, and “allelic series” of deletions arising from each beacon, can be tracked in a single experiment, Shred-seq enables highly scalable, multiplexed, genome-wide deletion scans that meet the criteria we outlined in the introduction: variable-length deletions across the genome, direct genotyping and compatibility with functional readouts.

### Proof-of-concept of Shred-seq in diploid HAP1 cells

As an initial proof-of-concept, we cloned a Shred-seq beacon library into a PiggyBac transposon vector and used PiggyBac transposase to integrate the library into diploid HAP1 cells. Diploid cells allow us to optimize Shred-seq without the confounding of selection pressures that might be expected to operate on deletions in haploid cells. We used fluorescence-activated cell sorting (FACS) to sort for GFP-positive cells that represent successful beacon integration, which yielded a bottlenecked cell population harboring 10,744 distinct barcodes, as assessed by amplicon sequencing (**Figure S1A**). To map integrations, we sequenced IVT transcripts from the outward facing T7 promoter, and associated 6,549 (63%) of barcode sequences with ≥1 uniquely mappable genomic locations. The mapped integration sites cover all chromosomes except for Y, which is not present in HAP1 cells (**Figure 1C**) (Essletzbichler et al. 2014). The vast majority (95%) of mapped barcodes were associated with a single genomic location. Barcodes mapping to multiple locations likely reflect transposon re-hopping events during S-phase (**Figure S1B**) (Chen et al. 1992).

Having mapped beacon integration sites with high confidence, we next induced deletions using a previously published crRNA targeting the eGFP coding sequence within the integrated beacons (Tan et al. 2022). Successful Cas3-mediated editing results in loss of GFP expression, enabling optional enrichment of deletion-containing cells by FACS (**Figure 1D**). Unlike Cas9, Cas3 editing requires coordinated delivery of five protein components (Cas3, together with Cascade subunits Cas5, Cas7, Cas8, Cas11) and the crRNA. We initially tested a single polycistronic construct encoding all five proteins on a single plasmid (Tan et al. 2022), but observed little GFP loss, indicating inefficient editing (**Figure S1C**). In contrast, delivery of the same components as individual plasmids at an optimized stoichiometry (Tan et al. 2022) yielded robust editing, with 34-36% GFP-negative cells compared to 3.4% in control cells lacking the targeting crRNA (**Figure 1E**).

To characterize Cas3-mediated deletions, we collected cell populations at days 3 and 15 (timeline relative to Cas3 transfection). To assess whether GFP loss reliably enriches deletion-containing cells, we also isolated GFP-negative cells by FACS at day 18 and collected samples at day 27 (**Figure 1D**). Deletion genotypes were recovered by IVT from the inward-facing T7 promoter, which generates transcripts containing both the beacon barcode and the newly formed genomic junction (**Figure 1A**). Following RT-PCR, these breakpoint-containing amplicons were sequenced.

Because Cas3 induces deletions of highly variable size, we implemented a flexible deletion-calling pipeline that relied on two complementary sequencing strategies (**Figure S2**). First, we sequenced full-length RT-PCR products using Oxford Nanopore (ONT) long-read sequencing, enabling direct identification of deletion junctions from split alignments between the beacon and genome (**Figure S2A**). Second, we implemented a short-read strategy in which Illumina sequencing captures the beacon barcode and the adjacent genomic sequence from each IVT product (**Figure S2B**). For short-read data, we applied stringent filters, retaining only reads that: (i) contained a barcode previously mapped to a unique integration site; (ii) aligned outside the beacon sequence; (iii) mapped to the same chromosome as the corresponding beacon; (iv) were located between 1000 bp and 500 kb from the mapped beacon position; and (v) were oriented consistently with the integration orientation of the beacon (**Figure S2C**).

In Cas3-treated samples collected on day 3 (or day 15), 2.1% (or 4.6%) of reads with mappable barcodes bore evidence of a deletion, representing a 472-fold (or 2,080-fold) enrichment relative to negative-control samples lacking the GFP-targeting crRNA (**Figure 1F**; **Figure S3**). Enrichment of GFP-negative cells further increased the fraction of deletion-supporting reads to 17% (or 25%) (**Figure 1F**). Deletion lengths exhibited a right-skewed, long-tailed distribution (**Figure 1G**; median 33.1 kb, mean 70.8 kb, standard deviation 90.8 kb; max 495 kb), consistent with previous reports (Tan et al. 2022). Notably, 24% of deletions were >100 kb. From this single experiment, we identified 3,603 unique deletions that summed to 255 Mb and covered 151 Mb of the reference human genome.

We next sought to validate our T7-IVT-based breakpoint mapping using an orthogonal, locus-restricted approach. We derived clonal lines from the bottlenecked PiggyBac beacon population and used amplicon sequencing to identify each clone’s beacon integrations (mean 1.8; range 1-4), all of which corresponded to a previously mapped genomic location (**Figure S4A**). We selected a clone with a single integration and confirmed heterozygous integration of the full-length beacon by PCR genotyping (Clone 1; **Figure S4B**). Following Cas3 editing and bottlenecking, we mapped deletion junctions either by long-range PCR followed by long-read sequencing of an 18-kb amplicon, or alternatively by T7-IVT-based genotyping (**Figure 1H**; **Figure S4C**). Both approaches detected deletions originating from the same integration site. However, the PCR-based strategy was constrained to the 14 deletions whose junctions fell within the primer-defined interval. In contrast, the IVT-based approach captured 12 of the 14 deletions detected by PCR amplicon sequencing, but also 48 larger deletions that extended beyond the PCR amplicon (**Figure 1H**). This underscores the advantages of the IVT-based approach, which enables: (i) deletion mapping from arbitrary start sites without requiring locus-specific primers; and (ii) recovery of deletions spanning a wide and unbounded size range.

Taken together, these results establish that Shred-seq enables the generation and high-resolution genotyping of thousands of variable-length deletions, originating from randomly integrated launchpads and ranging from ∼100 bp to >300 kb, in a single pooled experiment, without any reliance on clonal isolation or whole-genome sequencing.

### Optimized Shred-seq enables gigabase-scale deletion scanning

Having established the proof-of-concept of Shred-seq, we next sought to optimize the beacon sequence and architecture to boost T7 IVT levels and, correspondingly, the efficiency of deletion genotyping. For this, we first padded the T7 promoter with sequences previously reported to enhance its activity (v2 beacon, **Figure 2A**) (Conrad et al. 2020). We also hypothesized that the original design with convergent T7 promoters might hinder IVT through polymerase collisions (Callen et al. 2004; Ross et al. 2025), and replaced one T7 promoter with an orthogonal SP6 promoter (v3 beacon, **Figure 2A**). To evaluate performance, we generated cell lines with a mixture of all three beacon designs, linked each barcode to its beacon version via amplicon sequencing, and quantified T7-derived transcripts originating from each construct (**Figure S5A-C**). T7 promoter padding (v2) increased yield by 1.7-fold relative to the v1 beacon, while the combination of T7 promoter padding and non-convergent T7/SP6 promoters (v3) exhibited a dramatic 18.3-fold improvement over the v1 beacon (**Figure 2B**).

**Figure 2.**
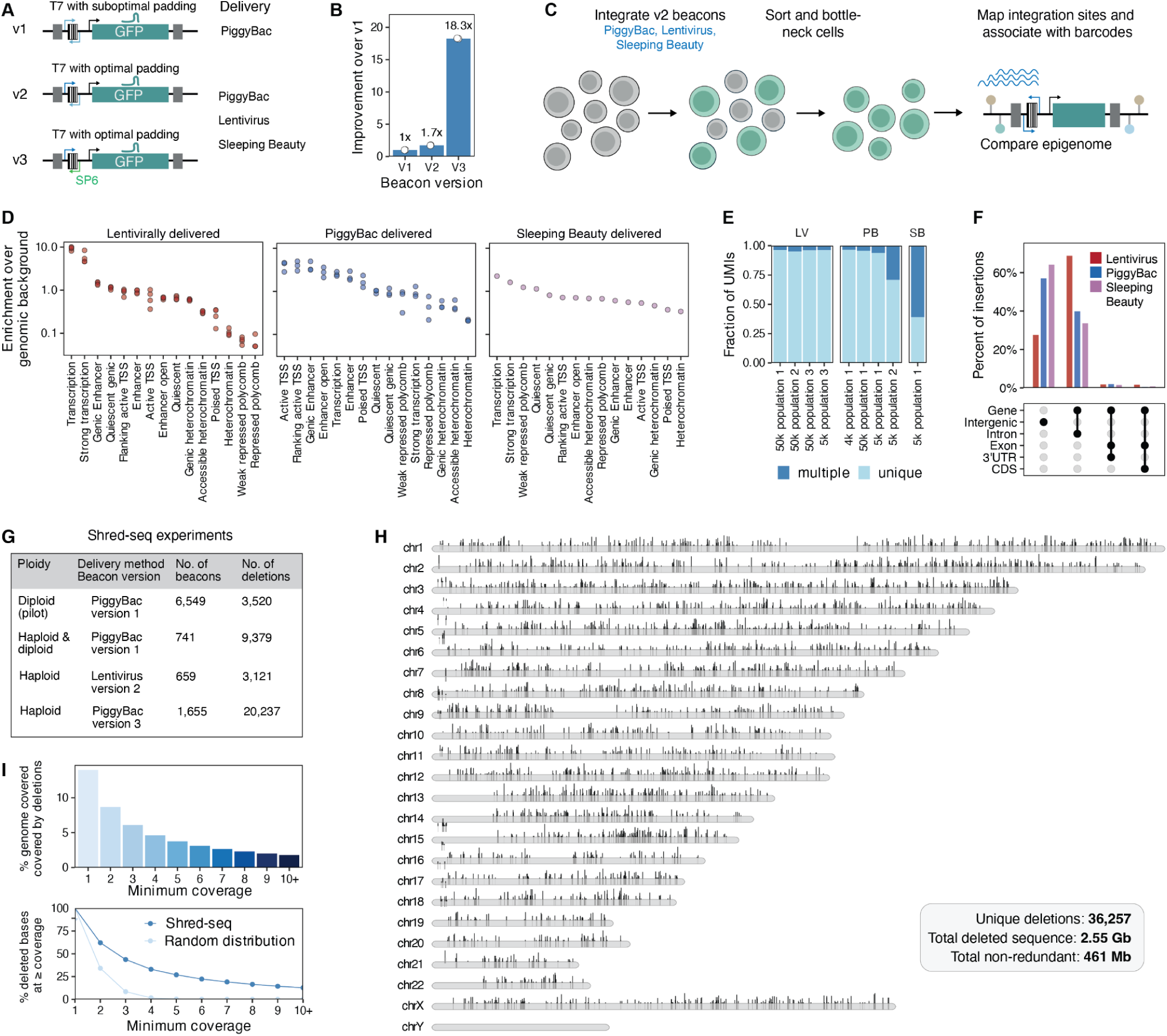
Optimized Shred-seq enables gigabase-scale deletion scanning. (**A**) Schematics of v1, v2 and v3 beacons. The v2 beacon optimizes the sequence context surrounding the T7 promoter, and the v3 beacon further replaces the upstream-oriented T7 promoter with the SP6 promoter to avoid polymerase collision. Additional v2 beacons were designed for delivery via lentiviral vectors or Sleeping Beauty transposase. (**B**) Barplot of T7 read recovery per barcode for various beacon designs, normalized to performance of the v1 beacon. Bars are the mean of two biological replicates, individually shown as dots. (**C**) v2 beacons were integrated with lentivirus, PiggyBac, or Sleeping Beauty. Cells were bottlenecked, and insertion sites were mapped. (**D**) Enrichment of integration sites over genomic background (y-axis) across different chromatin states (x-axis) for different delivery modalities (panels and colors). Dots represent independently bottlenecked populations. Columns are arranged from highest to lowest mean enrichment within each panel. (**E**) Barplot of the fraction of UMIs (y-axis) mapping uniquely or to multiple loci (colors) for each delivery method and independent bottlenecking (x-axis). (**F**) Percent of insertions (y-axis) across gene features for each delivery method (x-axis). Bars reflect the mean of 1 (Sleeping Beauty), 3 (PiggyBac), or 4 (lentivirus) independently bottlenecked populations. (**G**) Large-scale Shred-seq experiments included in the combined analysis. No. of beacons refers to the number of uniquely mapped beacons, and no. of deletions to the number of unique deletions detected. (**H**) Genome-wide distribution of deletions across all experiments and cell lines. Shaded areas on the ideogram indicate the largest deletion intervals for each beacon, and the height of the vertical bars on top of the ideogram corresponds to the log-scaled number of deletions at each beacon. (**I**) Top: The percent of the human reference genome covered (y-axis) at various coverage thresholds (x-axis) by deletions observed in the union of multiple large-scale Shred-seq experiments, including all timepoints and replicates. Bottom: The percent of deleted bases (y-axis) at various coverage thresholds (x-axis) for observed Shred-seq deletions and randomly distributed deletions matched for number and length (mean of 10 simulations).

PiggyBac has a known preference for open chromatin (Li et al. 2013). To explore whether alternate integration modalities might enhance genome coverage, we generated v2 beacon constructs for lentivirus and Sleeping Beauty transposase. We integrated the three v2 beacon subtypes into diploid HAP1 cells, sorted GFP-positive cells, and mapped integration sites in bottlenecked populations (**Figure 2C**). We then compared their genomic site preferences by quantifying the relative enrichment of chromatin states surrounding each integration coordinate (**Figure 2D**). In brief, all three systems favored open and active chromatin, but in distinctive ways. Lentiviral integration was 10-fold enriched in actively transcribed regions vs. genomic background, whereas PiggyBac was most enriched in sequences near transcription start sites and enhancers. Sleeping Beauty exhibited the most balanced profile overall but also a much stronger tendency for multiple integrations of the same barcode, making it less suitable for Shred-seq (**Figure 2E**). Heterochromatin and polycomb-repressed DNA were disfavored across all three delivery modalities.

We also examined the overlap of integration sites with transcribed genes (TPM > 1) in HAP1 cells. While lentiviral beacons were most commonly found in the introns of expressed genes (66%) PiggyBac and Sleeping Beauty-delivered beacons were predominantly intergenic (58-64%) (**Figure 2F**). Overall, these results indicate that the choice of delivery modality for Shred-seq ought to be determined by the experimental question. In particular, PiggyBac may be optimal for targeting regulatory sequences, whereas lentivirus may be better suited for the interrogation of actively expressed coding and non-coding genes.

With optimized beacons and diversified delivery modalities in hand, we scaled Shred-seq via three additional large-scale screens. For chronological reasons (*i.e.* the timing of when the aforedescribed optimizations were explored), these were conducted with PiggyBac v1, lentiviral v2 or PiggyBac v3 beacons in haploid and/or diploid HAP1 cells (**Figure 2G**; **Supplementary Table 1**). All analyses described below were performed using aggregated data from these four screens unless otherwise specified.

To investigate the selection pressures shaping SV survival, we collected cells from both early (pre-selection; day 3-4) and late (post-selection; 2-4 weeks of culturing without bottlenecking) timepoints. Altogether, we recovered 36,257 deletions spanning 2.55 Gb of cumulative deleted sequence, including 461 Mb of unique (non-overlapping) sequence (13.9% of the human reference genome; **Figure 2H**). The detection of regions under selection would be ideally supported by many independent deletion events, and Shred-seq was designed to facilitate such depth. Consistent with this, we identify 3.8% of the human reference genome with ≥5 independent deletions and 1.8% with ≥10 independent deletions (**Figure 2I**). In contrast, if we randomly distribute 36,257 deletion events with an identical size distribution and footprint (2.55 Gb), only 0.02% of the genome is expected to be covered by more than 5 deletions, 200-fold less than what we observed with Shred-seq, and ∼0% with ≥10 independent deletions (**Figure 2I**).

### Biology of Cas3-induced deletions in mammalian cells

To investigate how Cas3 interacts with native chromatin contexts and repair machinery, we initially focused on 19,529 deletions observed in pre-selection samples, and sought to assess: (i) the length distribution of induced deletions, (ii) precisely where deletions were initiated, (iii) how epigenomic context shapes both the initiation and termination of deletions; and (iv) how Cas3-induced deletions are repaired (**Figure 3A**).

**Figure 3.**
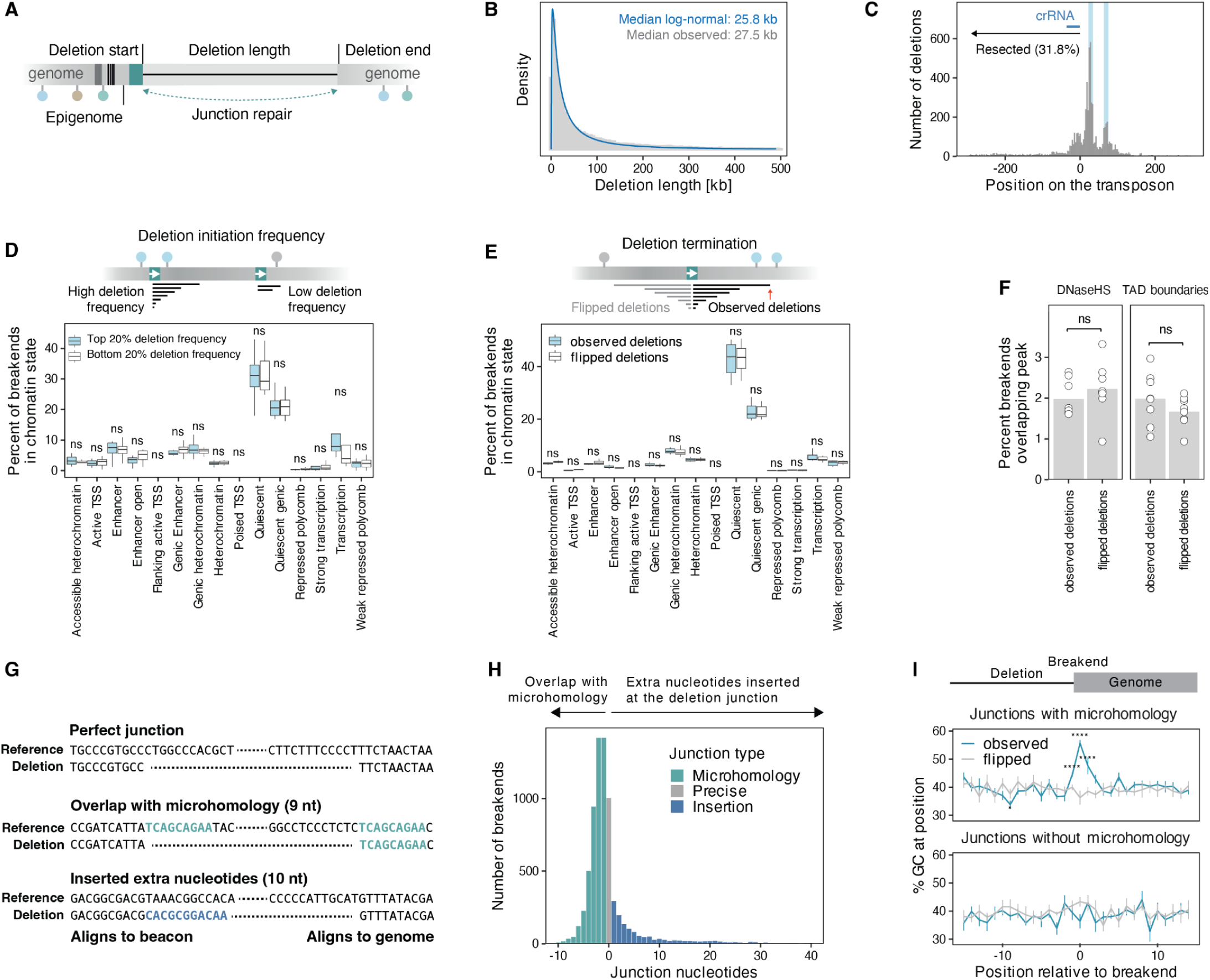
Biology of Cas3-induced deletions in mammalian cells. (**A**) Schematic highlighting features of Cas3-induced deletions, including: deletion start and end coordinates, deletion length, local epigenomic state at the breakpoints, and junction repair outcome. (**B**) Length distribution (x-axis) of all observed deletions (gray histogram) with a log-normal fit (blue line). (**C**) Histogram of the number of deletion start sites (y-axis) by position relative to beacon’s crRNA binding site (x-axis). Start sites that would destroy crRNA binding site are marked by ‘resected’ arrow. Blue shading highlights the hotspots discussed in text. (**D**) Impact of chromatin neighborhood on deletion initiation. Top: Schematic of beacon sites initiating many (left) or few (right) deletions. Bottom: Percent of beacon sites (y-axis) across different chromatin states (x-axis) for beacons initiating many (top 20%) or few (bottom 20%) deletions. P-values were computed using a student’s t-test and corrected for multiple hypothesis testing (Benjamini-Hochberg). FDR thresholds: ns, >0.05; * ≤0.05; ** ≤0.01; *** ≤0.001; **** ≤0.0001. (**E**) Impact of chromatin neighborhood on deletion termination. Top: Schematic of observed deletions (black) and a “flipped” null control (gray). Bottom: Percent of breakends (y-axis) across different chromatin states (x-axis) for observed deletions or flipped controls (colors). P-values and thresholds as in panel **D**. (**F**) Percent of deletion breakpoints that overlap with DNase I hypersensitive or TAD boundary peaks (y-axis) of HAP1 cells for observed deletions or flipped controls (x-axis). Dots are individual experiments. Bars are average across all experiments. P-values and thresholds as in panel **D**. (**G**) Representative junction sequences illustrating three repair classes. For each example, the top line shows the two reference sequences surrounding the initiation site within the beacon (left) and the termination site within the genome (right), while the bottom line shows the post-repair read, split in correspondence with its alignment to beacon and genome references. Microhomologous bases are highlighted in teal and inserted bases in blue. (**H**) Histogram showing the count of deletions (y-axis) across different junction nucleotide lengths (x-axis), with negative values indicating overlap. Deletions with >=1 overlapping bases were called as microhomologies, deletions with 0 overlapping bases were called as precise, and deletions with >=1 junction nucleotides were called as insertions. (**I**) Sequence composition around deletion breakends, stratified by repair outcome. Top: Schematic of a deletion breakend illustrating the coordinate system used in the plot below. Negative positions represent the genomic bases upstream of the breakend that are removed by the deletion, and positive positions represent retained genomic bases downstream of the breakend, with position 0 denoting the first retained genomic nucleotide. Bottom: Average GC content (y-axis) across a 30 bp window around the deletion breakend (x-axis) for observed deletions and flipped controls (lines and colors), separated by repair outcome (panels). Whiskers show the standard error of the mean. P-values were computed with a two-sided test for equality of proportions and corrected for multiple testing across positions (Benjamini-Hochberg).

We fit the length-distribution of Cas3-induced deletions using exponential, log-normal, or Weibull models (**Figure 3B**; **Figure S6**). The log-normal distribution had the best fit (lowest Akaike information criterion; **Figure S6**). Log-normal distributions can arise from processes driven by the product of multiple independent positive random steps. This distribution makes intuitive sense if the final lesions detected after Cas3 induction are considered the product of binding, unwinding, DNA strand cutting, and DNA repair. The fitted distribution had a median deletion length of 25.8 kb, close to the observed median of 27.5 kb.

Cas3 initiates most deletions within 200 bp of the crRNA target (Tan et al. 2022; Morisaka et al. 2019), but precise initiation sites have not been previously mapped at scale. We used split reads to map 7,467 deletion start sites (**Methods**). The majority of deletions initiated downstream of the protospacer binding site, with the strongest enrichment 22-34 nucleotides downstream of the last base-pairing nucleotide of the protospacer (25.1% of deletions, **Figure 3C**). A second peak was apparent further downstream (positions 63-75, 7.7% of deletions, **Figure 3C**), potentially corresponding to the binding of one additional Cas7 monomer, the periodicity of Cas3 helicase-nuclease activity, or biases introduced during the repair of Cas3-generated lesions. Of note, the initiation of deletion downstream of the protospacer leaves the binding site intact, which could result in Cas3 rebinding and initiating another round of deletions. Consistent with this, 2.3% of nanopore sequencing reads bore evidence of multiple deletions (**Figure S7A,B**). In contrast, if a deletion initiates within or upstream of the protospacer, or if a proximally downstream Cas3-induced lesion compromises the crRNA binding site, the protospacer can no longer bind, locking the current genotype into place, as is observed for 31.8% of deletions (**Figure 3C**). Deletion initiation patterns were consistent across different beacon versions, delivery modalities, and sequencing strategies (**Figure S7C**).

To assess whether local chromatin influences Cas3 deletion initiation, we quantified the deletion frequency of each beacon as the number of recovered deletions normalized by the beacon’s relative abundance in the bottlenecked population. We then contrasted the local chromatin environment (1 kb window centered on the integration site) for the top 20% vs. bottom 20% of beacons by deletion frequency. Surprisingly, no chromatin state showed a significant difference between high- and low-frequency beacons (**Figure 3D**). The absence of a detectable chromatin effect is in sharp contrast with Class 2 CRISPR systems such as Cas9 and Cas12, whose editing efficiencies are generally higher in open chromatin and actively transcribed regions (X. Li et al. 2024; Mathis et al. 2025; Verkuijl and Rots 2019; Strohkendl et al. 2021). However, because we selected for GFP-expressing beacons, we may have biased our cells towards those with reduced sensitivity of Cas3 activity to the local chromatin environment.

We also examined how the genomic context shapes deletion termination. For this, we generated a matched background by constructing a control set of “flipped” deletions that preserved each observed start site but extended in the opposite direction (**Figure 3E**). We then compared the local chromatin environment (1 kb windows) of the observed vs. flipped breakends, and again found no clear differences (**Figure 3E**). As Type I E Cas3 systems can be roadblocked by dead Cas9 (J. Li et al. 2024), we wondered if endogenously DNA-bound proteins could block Type I C Cas3. We intersected deletion endpoints with HAP1 DNase hypersensitivity sites and TAD boundaries (ENCODE Project Consortium 2012; Sanborn et al. 2015), but neither feature was enriched relative to flipped controls (**Figure 3F**). Overall, these results lead us to the conclusions that: (i) Cas3 is relatively insensitive to the local chromatin context in mammalian cells; (ii) endogenously DNA-bound proteins do not measurably impede Cas3 progression (Ishida et al. 2026).

### Repair outcomes of Cas3 lesions

To understand how Cas3-induced lesions are resolved in mammalian cells, we examined alignments of breakpoint-spanning reads to the beacon and genome sequences. We categorized junctions into three types: (i) perfect junctions, where beacon sequence transitions seamlessly into genome sequence; (ii) microhomology overlaps, where the terminal beacon nucleotides are identical to the first aligning genome nucleotides; and (iii) insertions, where extra nucleotides appear between beacon- and genome-aligning portions of the read (**Figure 3G**).

The majority of Cas3-induced breakpoint junctions (66.5%) show 1-10 bp overlaps consistent with microhomology-mediated end joining (MMEJ) (Sfeir and Symington 2015). The remaining junctions show inserted nucleotides (19.2%) or perfect transitions (14.1%), both characteristic of non-homologous end joining (NHEJ) (Chang et al. 2017). Overlaps exceeding 20 nucleotides, which would be indicative of homology-directed repair (HDR) (Liao et al. 2024), were rare (0.04%), although the likelihood of such extended overlaps between any given beacon sequence and downstream genomic regions is low. These proportions were consistent across beacon versions, delivery modalities, and sequencing strategies (**Figure S8A**), suggesting MMEJ as the predominant repair mechanism for Cas3-induced lesions in mammalian genomes, with the caveat that we may be underpowered for scenarios that would promote HDR-mediated repair.

To further characterize repair of Cas3-induced lesions, we examined GC content across a 30 bp window around deletion end sites. While *in silico* flipped control deletions (**Figure 3E**) exhibited a uniform GC content across the window (mean 40.2%, near genome-wide expectation of 40.9%) (Piovesan et al. 2019), observed deletions exhibited a sharp peak of up to 46.5% GC in the first three post-junction nucleotides (**Figure S8B**). These are the same nucleotides that would form a microhomology in MMEJ-mediated repair. Indeed, when we stratified the deletions by junction type (**Figure 3I**), we observed a pronounced peak in GC content for junctions with microhomology (>1 bp of homology, 56.3%) that vanished for junctions without microhomology (37.0%). Intriguingly, this is similar to patterns observed for unintended on-target long deletions from editing with Cas9 nucleases or paired Cas9D10A nickases (70% of junctions with microhomology; elevated GC content in region of microhomology) (Owens et al. 2019). These analyses further support the conclusion that MMEJ is a major contributor to the repair of Cas3-induced lesions.

### Selection pressures shape the characteristics of surviving deletions

To evaluate their impact on cellular viability, we compared deletions observed shortly after their generation (pre-selection) vs. after several weeks of cell culture (post-selection). Across four experiments in haploid and diploid HAP1 cells (**Figure 4A**; **Figure 2G**), there were 34,515 deletions spanning 2.42 Gb of cumulative deleted sequence (460 Mb of unique sequence spanning 13.9% of hg38) associated with experiments for which we had sampling from both pre-selection and post-selection timepoints.

**Figure 4.**
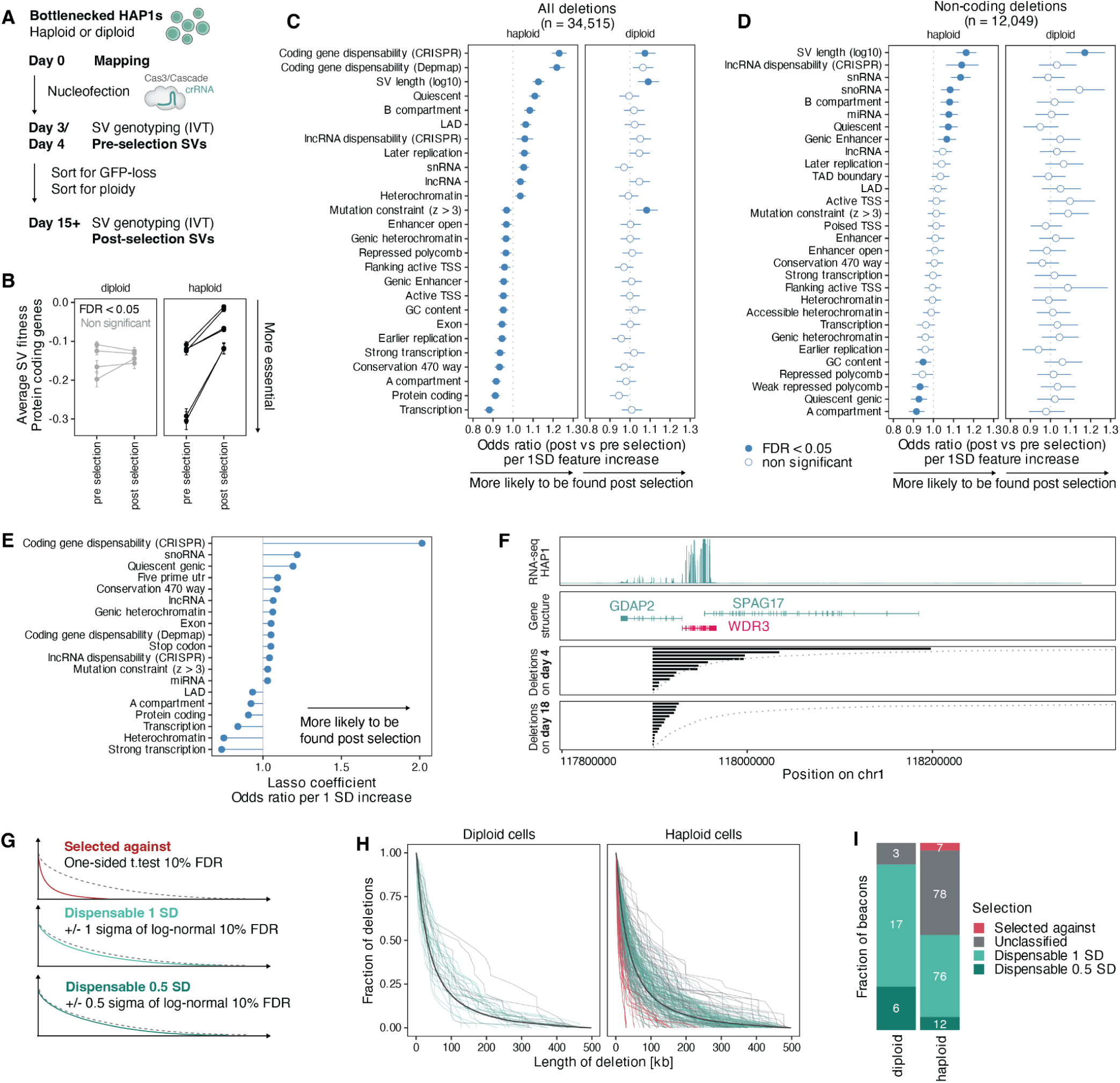
Selection pressures shape the characteristics of surviving deletions. (**A**) Schematic of Shred-seq experiments used for analysis of selection pressures. (**B**) Average deletion fitness (y-axis) across Shred-seq experiments and biological replicates (points and lines) pre- and post-selection (x-axis), separated by ploidy (panels), and shaded by FDR, following adjustment for multiple hypothesis testing. Whiskers represent standard error of mean. (**C**) Odds ratios per standard deviation increase (x-axis) in various features (y-axis) in regions overlapping pre-selection vs. post-selection deletions in haploid (left panel) or diploid (right panel) HAP1 cells. Whiskers indicate 95% confidence intervals. Unless indicated otherwise, the fraction of each variant covered by the respective feature is used as the statistic. Only features that are significant in haploid HAP1 cells are shown. (**D**) As in panel **C** but for the subset of 12,049 non-coding deletions. All features applicable to non-coding regions are shown. (**E)** Non-zero coefficients at the selected λ were extracted and displayed as odds ratios per one standard deviation increase in the corresponding feature. (**F**) Deletions and genomic features (panels) at representative region bearing an integrated beacon (x-axis). From top to bottom: (i) RNA-seq coverage in HAP1 cells. (ii) Exon structure and names of genes in region. (iii) Deletions observed on day 4, with each line corresponding to length of one deletion. (iv) Deletions observed on day 18. (**G**) Schematic of deletion profiles (colored lines) and their consistency with the log-normal distribution (dotted line). (**H**) Cumulative fraction of deletions (y-axis) by length (x-axis) at all post-selection beacon sites with >10 unique deletions (lines) in diploid and haploid cells (panels) according to consistency with log-normal distribution (colors). Solid line: expected log-normal distribution. (**I**) Fraction of beacons (y-axis) across ploidies (x-axis) according to their consistency with log-normal distribution (colors). Number of beacons representing that fraction are indicated within each section.

We first examined 15,907 (46.1%) deletions that overlapped protein-coding exons. We compiled gene fitness scores from CRISPR-Cas9 knockout screens in HAP1 cells, as well as from >1000 cancer cell lines from the Cancer Dependency Map (**Methods**) (Aregger et al. 2020; Arafeh et al. 2025). We defined the fitness of each deletion as the minimal fitness score among the genes that it overlapped. In haploid HAP1 cells, the average fitness score of post-selection deletions was significantly higher than that of pre-selection deletions (mean increase: 0.05-0.19, FDR: 2.4×10^-5^-9.0×10^-3^; **Figure 4B**), consistent with the depletion of surviving deletions for essential protein-coding genes. This shift was enhanced when we restricted the analysis to 6,600 (19.3%) deletions originating from beacons within 500 kb of a “core essential” gene (mean Chronos score < 0.5 across Cancer Dependency Map cell lines) (**Figure S9A**). In contrast, pre-selection vs. post-selection deletions exhibited no significant shift in mean fitness score in diploid HAP1 cells (**Figure 4B**), nor in haploid HAP1 cells if analysis is restricted to a residual 30 Mb diploid region (Essletzbichler et al. 2014), consistent with buffering by a second allele (**Figure S9B**). Separately, we leveraged long non-coding RNA (lncRNA) CRISPR-Cas13 knockout screens performed in HAP1 cells to calculate fitness scores for 1,927 (5.6%) deletions overlapping lncRNAs with detectable expression (TPM > 1; deletion fitness score defined as minimal fitness score among overlapping lncRNAs) (Liang et al. 2025). Once again, we observed a shift towards higher average fitness score in haploid HAP1 cells only, albeit of smaller magnitude than the shift observed for protein-coding genes (**Figure S9C**).

To systematically map which genomic features underlie selection, we compared the characteristics of 17,817 pre-selection vs. 16,698 post-selection deletions across 38 features spanning gene essentiality, chromatin states, functional elements, conservation, mutational constraint, replication timing, and genome organization (**Figure 4C**; **Figure S10A; Supplementary Table 2**) (ENCODE Project Consortium 2012; Christmas et al. 2023; Dekker et al. 2017; Pollard et al. 2010; Reiff et al. 2022; Harrow et al. 2012; Chen et al. 2024; Aregger et al. 2020; Arafeh et al. 2025; Liang et al. 2025). Surviving deletions in haploid HAP1 cells exhibited depletion for features associated with active chromatin (*e.g.* A compartments, transcription, early-replicating regions, functional elements), mutation-constrained DNA, and highly conserved DNA from a 470-way placental mammal alignment (Christmas et al. 2023). Conversely, surviving deletions were enriched for features associated with inactive chromatin (*e.g.* B compartments, quiescent or heterochromatin, lamina-associated domains, late replicating regions). These differences were largely absent when the same analysis was applied to pre-vs. post-selection deletions in diploid HAP1 cells, again consistent with buffering by a second allele (**Figure 4C**; **Figure S10A**). Interestingly, the mean length of surviving deletions was 1.1-fold greater in both haploid and diploid HAP1 cells, which may be due to the persistence of the transfected Cas3 plasmid beyond day 4 (**Figure 4C**).

Because many of these genomic features are correlated (**Figure S10B**), it is possible that the observed depletions and enrichments are exclusively driven by selection acting on essential protein-coding genes. To evaluate this, we repeated the feature analysis for 12,049 deletions that did not overlap any protein-coding gene. However, even in this non-coding subset, post-selection deletions were depleted for features associated with active chromatin (A compartment, earlier replicating, higher GC-content, polycomb repressed) and enriched for features associated with inactive chromatin (B compartment, later replicating, quiescent) (**Figure 4D**). A similar picture emerged when we included all deletions but used protein-coding gene essentiality as a covariate in calculating the odds ratios (**Figure S10C**). Together, these results indicate that selection is not driven solely by the disruption of essential protein-coding genes, and that non-coding deletions also experience measurable purifying selection.

To shed further light on how these correlated annotations jointly shape post-selection survival, we trained a multivariate elastic-net logistic regression model that included all features (**Methods**) (Friedman et al. 2010; Zou and Hastie 2005). Fitted coefficients recapitulated major axes seen in univariate analyses. Deletions overlapping dispensable coding genes and quiescent chromatin were more likely to survive, and those overlapping transcriptional annotations, A compartments, or protein-coding regions less likely (**Figure 4E**). These multivariate results reinforce our conclusion that the observed patterns of selection reflect the combined contribution of multiple, partially redundant genomic features, rather than any single feature.

### Deletion length distributions enable annotation-agnostic finemapping of essential regions

On average, Cas3-induced deletions follow a log-normal distribution (**Figure 3B**), but the presence of essential sequence elements downstream of the beacon is expected to constrain the deletion profile post-selection. For example, this expectation is met at a beacon site integrated 32 kb upstream of the core essential gene *WDR3* on chromosome 1 (**Figure 4F**). Deletions originating from this beacon in haploid HAP1 cells followed a log-normal profile on day 4, with 7/18 (39%) deletions overlapping *WDR3*. In contrast, 0/16 (0%) of post-selection deletions overlapped *WDR3,* and the distribution was significantly shifted from log-normal (p = 0.00013; **Figure 4F**). More such examples are shown in **Figure S11A-C**.

We wondered whether deviations from the baseline expectation that unselected deletions should follow a log-normal distribution could be used to more systematically identify specific regions under selection (**Figure 4G**). To do this, we first identified all beacon sites with >10 deletions at both pre- and post-selection time points (**Figure S12A**). Because our goal was to detect selection-driven distortions of deletion length profiles, rather than site-specific biases in Cas3 deletion generation, we excluded beacons whose pre-selection deletion lengths already deviated from the log-normal expectation (10% of sites; **Figure S12A-B**). Collectively, these filters left us with 26 beacons in the diploid data, and 173 beacons in the haploid data, with sufficient coverage for assessing deviation from the log-normal length distribution as a signature of essentiality.

In diploid cells, none of the analyzed post-selection beacons (0/26; 0%) significantly departed from the log-normal expectation (FDR < 10%), and most sites (23/26; 88%) remained within one standard deviation (**Figure S12B**; **Figure 4I**). By contrast, in haploid cells, 7/173 beacons (4%) were significant (FDR < 10%), and a reduced proportion (88/173; 51%) remained within one standard deviation (**Figure 4H,I**). For 6 of the 7 significant haploid beacons, the constraint can be attributed to an essential protein-coding gene downstream of the beacon (examples in **Figure S11**). The remaining haploid-constrained beacon fell within a region dense in highly transcribed genes, although none of the downstream genes were themselves classified as nominally essential (chr3:49419658; **Supplementary Table 3**; **Supplementary Note 1**).

The coincidence of deletion length distribution deviations and known essential genes suggests that we can leverage these results to place lower and upper bounds on the proportion of the genome that is “dispensable” in a manner that is agnostic to preexisting annotations. This contrasts with CRISPR essentiality screens, which are overwhelmingly limited to frameshift disruptions of protein-coding genes, focal disruptions of candidate non-coding regulatory elements, or tiling disruptions of selected genomic regions (Gasperini et al. 2017; Aregger et al. 2020; Herger et al. 2025). Specifically, our results imply a lower bound of ∼50% on the proportion of genomic regions that are dispensable for the proliferation of haploid cells in culture and an upper bound of ∼96%. Of course, this assumes the 173 beacons are representative, and resolution is limited by the density of beacons and the number of independent deletions recovered per beacon. However, obtaining more accurate estimates and, eventually, a genome-wide, high-resolution map of haploid essentiality, may simply be a matter of scaling the application of Shred-seq to more beacons, cells and mapped deletions.

### Single-cell profiling links Shred-seq deletion genotypes to transcriptional phenotypes

Our experiments so far have relied on viability in cell culture as the functional readout. However, this readout only captures a narrow subset of potential phenotypic consequences. For example, induced deletions may produce transcriptional effects *in cis* or broader cellular changes *in trans* that do not appreciably impact cell viability. As a central feature of Shred-seq is the use of phage promoters to transcribe deletion junctions into RNA, we reasoned that the same architecture could be leveraged to capture deletion genotypes within single-cell assays (*e.g.* scRNA-seq), enabling broader phenotypic scope.

Adapting Shred-seq to scRNA-seq (‘sc-Shred-seq’) required addressing several key technical challenges. First, most scRNA-seq workflows rely on poly(A)-tail capture during reverse transcription, but transcripts generated by T7 IST are not polyadenylated. Second, because deletion junctions occur at unpredictable positions, they cannot be captured using fixed sequence-specific primers of the kind widely used to recover guide RNAs (Replogle et al. 2020; Mimitou et al. 2019) or transcribed junctions (Pinglay et al. 2025).

We reasoned that both of these challenges could be solved by a single intervention: *in situ* polyadenylation with poly-A polymerase, which has previously been used to recover non-polyadenylated endogenous RNAs in single-cell workflows (Dinçaslan et al. 2024; McKellar et al. 2023; Isakova et al. 2026). Appending a poly(A) tail to T7-derived transcripts would allow them to be captured by oligo-dT priming alongside endogenous mRNAs, regardless of where within the transcript the deletion junction falls, thereby yielding matched deletion genotypes and whole-transcriptome profiles from the same cells (**Figure 5A**).

**Figure 5.**
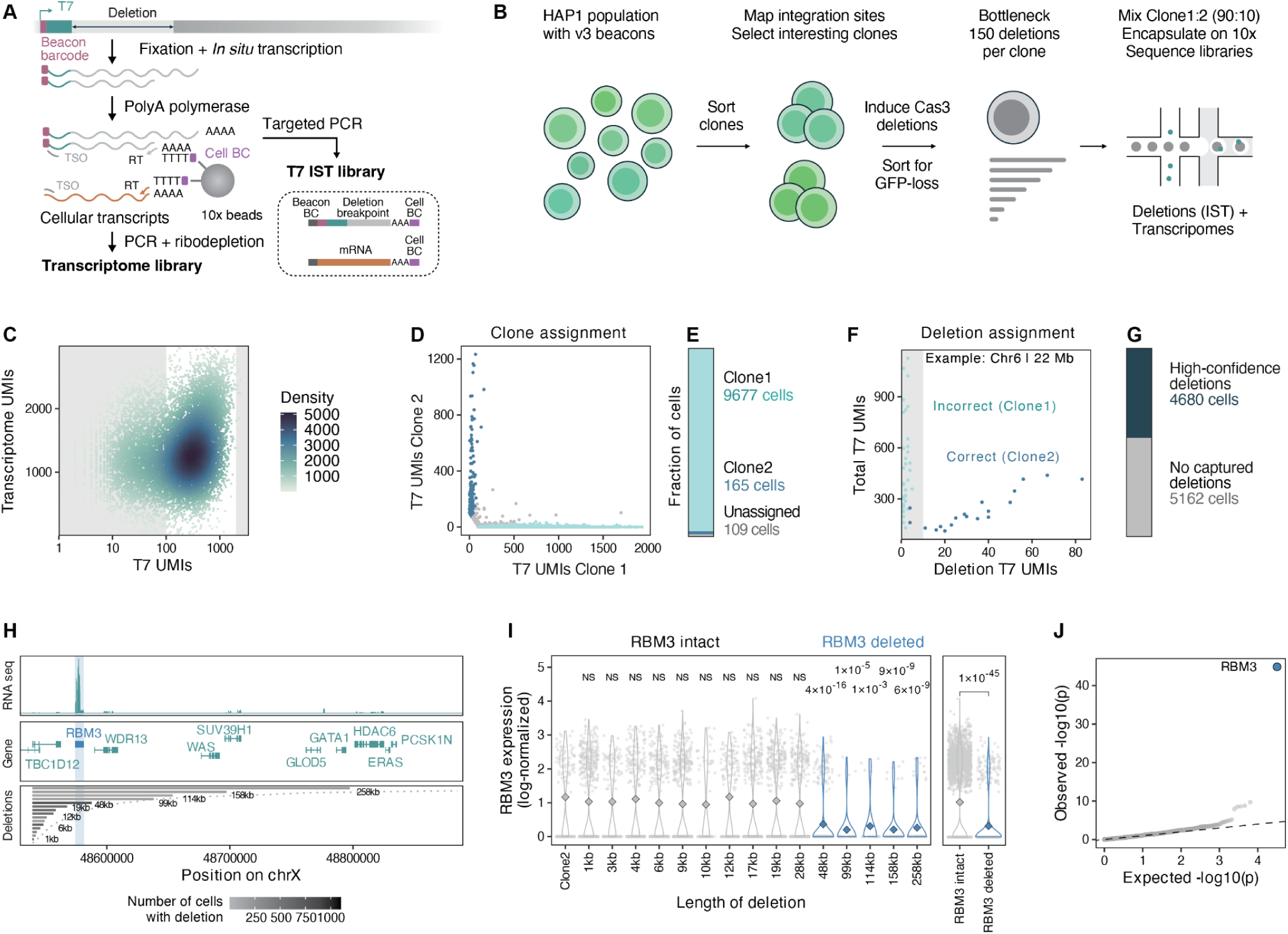
Measuring deletion-associated transcriptional changes in single-cells. (**A**) Schematic of scShred-seq workflow. After fixation and IST from beacon-associated T7 promoters, deletion-derived transcripts are polyadenylated and captured together with endogenous cellular mRNAs on beads (10x Genomics). Separate libraries are generated for IST products and transcriptome. (**B**) Proof-of-concept experiment. A HAP1 population carrying v3 Shred-seq beacons was single-cell sorted to isolate clones, integration sites were mapped, and selected clones were expanded. Cas3 deletions were induced, and 150 cells with reduced GFP expression were sorted, expanded and profiled with the scShred-seq workflow shown in panel **A**. (**C**) Distribution of T7 IST UMIs (x-axis) and transcriptome UMIs (y-axis) per cell barcode (dots). Dots colored based on number of neighbors (density). Cells in shaded area were excluded from further analysis. (**D**) Barnyard analysis of UMIs assigned for clone 1 (x-axis) and clone 2 (y-axis) based on beacon barcodes. Points represent cells, colored by clone assignment. (**E**) Barplot showing proportion of cells assigned to clones or unassigned (colors). (**F**) Deletion-specific T7 UMIs (x-axis) and total T7 UMIs (y-axis) for a representative clone 2 deletion on chr6. Cells (dots) colored based on matching of deletion assignment and clone identity. The shaded region, which corresponds to cells falling below our requirement for at least 10 deletion-supporting UMIs, includes all incorrectly assigned cells for this particular deletion. (**G**) Barplot showing the proportion of cells passing clone assignment for which a deletion genotype could be confidently assigned. (**H**) Genomic features and deletions (panels) at a region on chromosome X bearing an integrated beacon (x-axis; shading indicates location of beacon). From top to bottom: (i) RNA-seq coverage in HAP1 cells. (ii) Exon structure and names of genes in region. (iii) Deletions observed in the scShred-seq experiment with ≥ 20 cells, with each line corresponding to deletion length and shaded according to the number of confidently genotyped cells. (**I**) *RBM3* expression (y-axis) for cells based on deletion status (x-axis). Deletions overlapping *RBM3* are highlighted in blue. Markers represent individual cells, violins show the distribution, and the diamond represents the mean expression. P-values (two-tailed Wilcoxon test; Benjamini-Hochberg adjustment for multiple hypothesis testing), comparing clone 1 cells with indicated deletions to clone 2 cells. The right panel compares cells with deletions at this locus that spare vs. overlap *RBM3*. (**J**) Quantile-quantile plot depicting expected (x-axis) vs observed p-values (y-axis) from genome-wide differential expression testing (two-tailed Wilcoxon test) between cells with deletions at this locus that spare vs. overlap *RBM3.* Points represent individual genes and the dashed line shows the expected distribution under null hypothesis.

A third anticipated technical challenge was that *in situ* transcription (IST) at chromatinized genomic DNA in fixed cells is unlikely to be as efficient as IVT on purified genomic DNA. In the bulk Shred-seq experiments described above, most deletions initiated ∼500 bp downstream of the T7 promoter (**Figure 3C**). As such, any short T7-derived transcripts would fail to contain both the beacon barcode and deletion junction. To address this, we identified a crRNA target site 107 bp downstream of the T7 promoter (“crRNA-107”) for which the major deletion-initiation window was expected to fall 129-141 bp downstream of the promoter, thereby increasing the likelihood that IST-derived T7 transcripts would cover the deletion breakpoint (**Figure S13A**). This guide retained robust activity, as measured by GFP depletion (**Figure S13B**).

In order to be well-powered to detect transcriptional effects in this proof-of-concept, we sought to limit the number of beacons and Shred-seq-induced deletions. We began by isolating a GFP-positive clone with 3 mapped beacon integrations (“clone 1”), together with a second GFP-positive clone with 5 mapped beacon integrations (“clone 2”) (**Figure S13C**). After inducing Cas3 deletions with crRNA-107, we sorted and expanded ∼150 cells with reduced GFP expression (deletion-bearing) from each clone. Because one of its beacons was located near a cluster of highly expressed, non-essential genes (chrX: 48,539,463), we focused on scRNA-seq of post-sorting clone 1 cells, mixing in a smaller proportion (10%) of post-sorting clone 2 as an internal control (**Figure 5B; Figure S13C**). We then generated matched transcriptome and T7 IST libraries from 20,000 input cells (**Figure 5A**; 10x Genomics). After QC filtering (**Methods**), we retained scRNA-seq profiles for 13,944 cells (median 1237 transcriptome UMIs per cell; median 1014 genes detected per cell; **Figure S14A-C**). We focused our subsequent analyses on the 9,951 (71%) of these that also had high recovery of T7 IST reads (≥ 100; median 362; **Figure 5C, Figure S14D**).

Because we had already mapped the barcodes present in each clone, simply tallying counts was sufficient to infer the clone-of-origin of each profiled cell, effectively an inline barnyard control (**Figure 5D**; **Figure S15A**). Upon filtering potential doublets, we were able to assign 9,842 scRNA-seq profiles (98.9%) to a single clone-of-origin (**Figure 5E**). Dimensionality reduction of the transcriptome data by Uniform Manifold Approximation and Projection (UMAP) showed a largely homogenous population, as expected for a single cell line, with no overt segregation by clone-of-origin (**Figure S15B**).

We called deletions as previously (**Figure S2**), and then assigned called deletion junctions to single-cells. The clone 2 spike-in provided a means of estimating the reliability of these assignments, which might be confounded by ambient T7 IST transcripts or barcode-swapping artifacts. Valid clone 2 deletion junction calls should be associated with cells assigned to clone 2 based on their beacon barcode, whereas spurious clone 2 deletion junction calls should predominantly be associated with cells assigned to clone 1, as these comprise the vast majority of the experiment (**Figure 5E**). We indeed observe this for cells with few UMIs supporting a clone 2 deletion junction, while cells with greater support were correctly assigned to clone 2 (**Figure S16**; representative clone 2 deletion shown in **Figure 5F**). Based on this observation, we required at least 10 deletion-supporting UMIs per cell, which largely eliminated clone-deletion mismatches. Applying this threshold across all deletions yielded 4,680 cells with high-confidence deletion annotations, corresponding to 33.6% of all recovered transcriptomes (**Figure 5G**).

We identified 26 unique deletions from clone 1, and none from clone 2, for which we had profiled ≥ 20 cells (mean 201 cells per deletion). Removing the minimum cell requirement increased these counts only to 31 (clone 1) and 10 (clone 2). The three clone 1 beacons were associated with 5, 6, and 15 deletions (**Figure S17A-B**). Each clone 1 cell can carry up to three independent deletions (*i.e.* one from each beacon), and we identified 582 cells (12.7%) with evidence for compound deletions, spanning 151 of 325 possible unique pairs. However, just four deletion pairs were present at markedly higher abundances and accounted for the majority of compound genotypes (335/582; mean Jaccard index of 0.259 for these top four vs. mean 0.005 for all other pairs). Overall, these data are consistent with the presence of at least ∼23 singleton and ∼4 compound deletions at the time of bottlenecking (**Figure S17C-D**).

Together, these results show that scShred-seq can confidently recover both single and compound deletion genotypes from individual cells. We also asked whether single-cell deletion calls were associated with the expected loss of GFP expression from the originating beacon. Indeed, for clone 1 cells with genotyped deletions, we we observed a 21% (single deletion) or 65% (compound deletion) reduction in log-normalized GFP expression, relative to all other clone 1 cells (Wilcoxon test, p = 2.6×10^-7^ for single deletions, p = 2.8×10^-8^ for compound deletions; **Figure S17E**). These clear reductions in GFP expression confirm that scShred-seq calls correspond to bona fide loss-of-function events.

To explore whether scShred-seq can detect deletion-dependent changes in endogenous gene expression, we focused on an intergenic clone 1 beacon positioned within a gene-dense region on chromosome X that initiated 15 unique deletions (28-483 cells; mean 239) of varying length (1-258 kb, median 17 kb) (**Figure 5H**). This allelic series included genotypes that fully deleted *RBM3*, a highly expressed RNA-binding protein involved in hypothermic stress responses, as well as others that fully spared it. All five *RBM3*-overlapping deletions, but none of ten *RBM3-*sparing deletions, significantly reduced *RBM3* expression when compared with clone 2 cells, which lack a beacon at this site (**Figure 5I**; Wilcoxon test, adjusted p = 1.3×10^-5^ to 3.7×10^-16^). Differential expression analysis comparing cells carrying *RBM3*-overlapping deletions with cells carrying *RBM3-*sparing deletions further identified *RBM3* as the strongest hit genome-wide (p = 1.4×10^-45^; **Figure 5J**; **Figure S18A**).

The same allelic series also captured a weaker but directionally consistent effect at *WDR13*, a modestly expressed gene in the same region. All four *WDR13*-overlapping deletions showed reduced *WDR13* expression relative to both clone 2 cells and deletions that spared *WDR13* (**Figure S18B-C**). Although individual deletion-level tests did not reach statistical significance, likely because *WDR13* was detected in fewer cells, pooled analysis across *WDR13*-overlapping deletions revealed significant downregulation compared with non-overlapping deletions (p = 0.01). Together, these results provide proof of principle that scShred-seq can jointly recover allelic deletion series and their transcriptional consequences at single-cell resolution, enabling its application to the systematic dissection of non-coding sequence function beyond fitness-based readouts.

## DISCUSSION

Genetic screens — whether through random or programmed mutagenesis, in cell lines or model organisms, with organismal, fitness, or molecular readouts — lie at the very heart of how our field has approached the systematic dissection of biology in organisms ranging from bacteria to humans. For the human genome, the limitations of available methods (gene traps, RNAi, CRISPR) are such that genetic screens have overwhelmingly been confined to the genome-wide disruption of open reading frames, leaving the vast non-coding majority of the genome unexplored in a systematic manner.

To address this gap, we developed Shred-seq, which couples CRISPR-Cas3-mediated deletions and phage polymerase-based genotyping to enable gigabase-scale genome-wide deletion scans. In the course of reducing the method to practice, we characterized the determinants of Cas3 editing in human cells, and scaled to 36,257 unique deletions collectively scanning 14% of the human genome with ∼10-fold coverage of the assayed intervals. This depth enabled quantification of selection pressures in an annotation-agnostic manner, revealing purifying selection not only on protein-coding genes but also, independently, on conserved, mutation-constrained, and biochemically active non-coding sequences. At the same time, leveraging deviations from the expected log-normal deletion length distribution to fine-map essential regions, we estimate empirical bounds of 50% (lower) and 96% (upper) on the proportion of the haploid genome that is dispensable for cellular viability. Finally, we integrate Shred-seq and scRNA-seq, opening the door to phenotypic readouts beyond bulk viability.

Shred-seq, in its present form, has several limitations that merit emphasis. First, our reliance on flow sorting constrains scalability. Alternatives, such as magnetic sorting or counter-selectable markers to enrich for deletion-bearing cells, and drug- or filtration-based methods to maintain haploidy, may prove more scalable (Qu et al. 2018; Freimann and Wutz 2017; Olbrich et al. 2019; Li et al. 2021; Haney et al. 2018). Second, certain applications may require different deletion length distributions than Cas3 naturally produces; protein engineering of Cas3/Cascade or alternative nucleases could enable more controlled deletion sizes. Third, although scShred-seq enables transcriptional phenotyping of individual deletions, the scale and statistical power of such experiments remain constrained by the cost and throughput of scRNA-seq. As with all single-cell perturbation screens, detecting subtle effects, rare phenotypes, or combinatorial interactions will require greater statistical power. This may be achievable through a combination of more beacons per cell and shifting to lower-cost, higher-throughput, higher-sensitivity single-cell molecular profiling platforms (Gasperini et al. 2019; Cusanovich et al. 2018; Bradu et al. 2026). Finally, while we are encouraged that both Cas3 editing and T7-based genotyping function across diverse cellular contexts including pluripotent stem cells (Tan et al. 2022; Pinglay et al. 2025; Dolan et al. 2019; Lu et al. 2024), the experiments presented here are limited to a single, near-haploid, cancer-derived cell line. Broader deployment across cell types, developmental stages, and environments will be needed to build a comprehensive picture of genome essentiality.

These limitations notwithstanding, we envision several applications for Shred-seq in the future. First, scaling to genome-wide saturating coverage would refine the empirical bounds presented here and provide a direct experimental framework for revisiting a long-standing question in genome biology -- what fraction of the human genome is required for cellular viability and function? Unlike approaches that infer function indirectly from conservation, biochemical activity, or existing annotations, Shred-seq screens empirically test the consequences of deleting contiguous genomic intervals across the genome, and at saturating coverage could uncover novel essential sequence elements that have escaped existing annotations as well as refine the boundaries of known essential regions. Performing Shred-seq screens across multiple cell types, or in stem cells subsequently differentiated into diverse lineages, could further define how the essential fraction of the genome changes across cell types and developmental stages.

Second, beyond genome-wide scans, where signals for subtle locus-specific effects may be underpowered, a targeted implementation of Shred-seq may be helpful for mapping essential enhancers and/or probing the relevance of regulatory element spacing. Targeted Shred-seq may also be useful for the fine-mapping of causal sub-sequences within broader SV regions implicated in human disease, a longstanding challenge in human genetics (Gasperini et al. 2017; Su et al. 2000). Recurrent SVs, typically mediated by segmental duplications, often span multiple genes and regulatory elements, *e.g.* autism-associated 16p11.2 deletions (Chung et al. 2021) and cancer-associated 9p21.3 deletions (Novara et al. 2009; Zhao et al. 2025). The targeted delivery of multiple Shred-seq beacons, followed by the generation and phenotyping of densely nested deletions, could facilitate the fine-mapping of causal sequence(s) within such loci.

Third, the emerging vision of genomic and cellular AI models, inclusive of both generative genomes and virtual cells, depends on training data that capture the consequences of perturbing biology (Bunne et al. 2024; Roohani et al. 2025). To date, such data are dominated by Perturb-seq screens, which are overwhelmingly focused on perturbing protein-coding genes (Replogle et al. 2022; Roohani et al. 2024). The non-coding genome, regulatory architecture, and larger-scale features of genome organization remain grossly unrepresented, leaving these models reliant on evolutionarily derived sequences shaped by selection and demographic history, with limited capacity to generalize beyond the distribution of naturally observed sequences. Shred-seq begins to address this gap through the systematic generation of large-scale perturbation-phenotype pairs — deletions of varying length across diverse genomic contexts, linked to fitness and molecular readouts. Such data, especially if complemented by analogous data from genome shuffling (Pinglay et al. 2025; Koeppel et al. 2025), could serve as training material for next-generation models with improved ability to predict variant effects across scale (*i.e.* from point mutations to SVs) while also enabling programmable biology in research, synthetic biology, and clinical settings.

Finally, the ability to systematically test which genomic sequences are dispensable lays the groundwork for the design of a viable cell line bearing a minimal version of the human genome, which we anticipate could have broad applications in synthetic biology, gene and cell therapy, and bioproduction. Previous bacterial minimal genome efforts, such as JCVI-syn3.0, determined essentiality through ORF-level analysis of transposon mutagenesis data — an approach poorly suited to mammalian genomes, where the vast majority of sequence is non-coding (Hutchison et al. 2016). Shred-seq directly addresses this gap, and the saturating genome-wide scans proposed above would provide a roadmap for both top-down (progressive deletion from an existing genome) and bottom-up (*de novo* genome synthesis) strategies for constructing a minimal human genome.

## AUTHOR INFORMATION

## Acknowledgments

We thank the members of the Pinglay and Shendure Labs for helpful discussions. We thank Yan Zhang, Zhonggang Hou, and Renke Tan for valuable advice on performing optimal NlaCas3 experiments and for sharing resources. We thank the Allen Institute’s Immunology team for allowing us to use their flow cytometry analyzers and sorters, particularly Vaish Parthasarathy, Julian Reading, Veronica Hernandez, and Tyanna Stuckey, for providing training and assistance. We thank the Allen Institute’s Brain Science team for NextSeq 2000 access. This work was supported by the NIH (RM1-HG009491 subaward to S.P., DP5-OD036167 to S.P., R01HG010632 to J.S.), Chan Zuckerberg Initiative (award numbers CZIF2024-349901, CZIF2025-011511 to S.P.), Defense Advanced Research Projects Agency (DARPA) (contract number HR0011263E037 to S.P.), the Seattle Hub for Synthetic Biology, a collaboration between the Allen Institute, Chan Zuckerberg Initiative and University of Washington (award number CZIF2023-008738 to J.S.), and the Brotman Baty Institute for Precision Medicine (S.P., J.S.). J.K. is supported by an EMBO Postdoctoral Fellowship (ALTF-717-2024). E.A. is supported by an ARCS Foundation scholar award. J.S. is an Investigator of the Howard Hughes Medical Institute.

## Author contributions

Conceptualized and initiated the study: S.P., J.S., J.K.

Performed experiments: J.K., S.S., A.K., R.D., P.C., with help from F.Q., Z.S.

Analyzed the data: J.K. with help from E.A.

Supervised the project: S.P., J.S.

Wrote the manuscript: J.K., S.P., J.S., with support from all authors.

## AI disclosure statement

We disclose that language editing, proofreading and coding were supported by AI-based tools; these were not used for conceptual development or primary manuscript writing. The authors take full responsibility for the contents of this manuscript.

## Competing interests

J.S. is on the scientific advisory board, a consultant, and/or a co-founder of Guardant Health, Phase Genomics, Sixth Street Capital, Pacific Biosciences, Cellular Intelligence and 10x Genomics. All other authors declare no competing interests.

## METHODS

### Cell lines

Haploid HAP1 cells were purchased from Haplogen and cultured in IMDM (Gibco, 12440061), supplemented with 10% FBS (Fisher Scientific, SH3039603), 100 U ml^−1^ penicillin, and 100 µg ml^−1^ streptomycin (Gibco, 15140163) at 37 °C and 5% CO_2_.

Diploid wild-type HAP1 cells were purchased from Horizon Discovery (C631) and cultured in IMDM (Gibco, 12440061), supplemented with 10% FBS (Fisher Scientific, SH3039603), 100 U ml^−1^ penicillin, and 100 µg ml^−1^ streptomycin (Gibco, 15140163) at 37 °C and 5% CO_2_.

HEK293T cells were purchased from ATCC (CRL-3216) and cultured in Advanced DMEM (Gibco, 12491015) supplemented with 10% FBS (Fisher Scientific, SH3039603), 100 U ml^−1^ penicillin, and 100 µg ml^−1^ streptomycin (Gibco, 15140163) at 37 °C and 5% CO_2_.

### Assessment of HAP1 ploidy and enrichment for haploid cells

Wild-type and treated HAP1 cells were trypsinized (Trypsin-EDTA (0.05%) phenol red, Thermo Fisher Scientific, 25300054), counted, and collected by centrifugation (300g, 3 mins). 500,000 cells were resuspended in 1mL IMDM media containing Hoechst-33342 (Invitrogen) at a concentration of 1:2000 and incubated at 37 °C and 5% CO_2_ for 2 hours in a 96-Well Round Bottom Plate (Corning). DNA staining of the treated samples was compared to known haploid and diploid populations via flow cytometry (Channel PB450). For a mixed haploid and diploid cell population, a trimodal histogram is expected, corresponding to the following cell state/cycle stages: haploid G1; haploid G2 or diploid G1; diploid G2. Haploid cells were gated and sorted based upon the peak corresponding to haploid G1, as well as low forward and side scatter to select for smaller cells.

### Flow cytometry

Samples were run on the CytoFLEX Flow Cytometer (Beckman) for analysis. The data was acquired with the CytExpert software and analyzed with Floreada.io. Events were first gated for cells based on forward and side scatter. Next, singlets were distinguished from doublets based on the width and height of the side scatter. Finally, cells were analyzed for their respective fluorescence channels. Sensitivity was set such that the mean fluorescence intensity of the negative population was around 10^1^ - 10^3^. Cell sorting was performed on the BD FACSMelody Cell Sorter.

### Molecular cloning

#### Beacon cloning

v1 beacon sequences were purchased as gene fragments (IDT, Supplementary Table 4) with a cloning site to enable barcode insertion. The gene fragment was cloned into a minimal PiggyBac backbone (X. Li et al. 2024, Plasmid 1, **Supplementary Table 6**) by restriction cloning. The backbone and gene fragment were digested with NheI-HF (New England Biolabs (NEB), R3131S) and SpeI-HF (NEB, R3133L), and the backbone was additionally dephosphorylated with QuickCIP (NEB, M0525S). Isolated backbone (Zymoclean Gel DNA Recovery Kit, Zymo Research, D4008) and gene fragment (Zymo DNA Clean-up and Concentration kit, Zymo Research, D4004) were ligated using T4 DNA ligase (NEB, M0202L), heat-inactivated, and transformed into NEB stable chemically competent *E. coli* cells (NEB, C3040H). To add a 12N barcode, an ssDNA oligonucleotide with degenerate bases and 24 nt of overlapping sequence (oligo 1, Supplementary Table 5) was ordered from Integrated DNA Technologies (IDT). The intermediate v1 beacon backbone was digested with BsmBI-v2 (NEB, R0739L) and the ssDNA oligo inserted using Gibson Assembly. Briefly, a 20 µL reaction with NEBuilder HiFi DNA Assembly Master Mix (E2621S), 1 pmol of ssDNA oligo, 0.005 pmol of digested backbone, and nuclease-free water was set up on ice and incubated at 50 °C for 60 minutes. The reaction was cleaned up (Zymo DNA Clean-up and Concentration kit, Zymo Research, D4008) and eluted in 6 µL of nuclease-free water. 2 µL of the clean Gibson reaction were electroporated into electrocompetent NEB 10-beta *E. coli* cells (NEB, C3020K). A diversity of > 300,000 and a background of < 5% were maintained for all libraries. The plasmid sequence is available as Plasmid 2 in Supplementary Table 6.

#### v2 beacon sequences (PiggyBac, Lentivirus, Sleeping Beauty)

All v2 beacon sequences were cloned in the following principal steps with construct-specific details noted below: (i) A donor backbone for the respective delivery method was linearized by restriction digest. (ii) A v2 insert with backbone-compatible overlaps was generated by PCR using the v1 beacon sequence as template (Plasmid 2, Supplementary Table 6). The forward primers were designed to bind the CS2 sequence on the template and contained an overhang with the improved T7 promoter that had the optimal padding sequence. (iii) Inserts were cloned into backbones by Gibson assembly using 2x NEBuilder HiFi DNA Assembly master mix (E2621S) following the manufacturer’s instructions. (iv) The Gibson reaction was purified (Zymo DNA Clean-up and Concentration kit) and electroporated into electrocompetent NEB beta *E. coli* cells (NEB, C3020K). A diversity of > 300,000 and a background of < 5% were maintained for all libraries. <u>PiggyBac</u>: The v1 PiggyBac backbone (Plasmid 2, Supplementary Table 6) was digested with SpeI and Bsp119I. The v2 PiggyBac beacon integration cassette was generated by PCR using oligos 2+3 (Supplementary Table 5). <u>Lentivirus</u>: A lentiviral backbone (Plasmid 3, Supplementary Table 6) was digested with SpeI and Bsp1407I. The v2 beacon integration cassette was generated by PCR using oligos 4+5 (Supplementary Table 5). <u>Sleeping Beauty</u>: A Sleeping Beauty backbone (Plasmid 4, Supplementary Table 6) was digested with XbaI and KflI. The v2 beacon integration cassette was generated by PCR using oligos 6+7 (**Supplementary Table 5**). The v2 beacon sequences are available as Plasmids 5-7 in **Supplementary Table 6**.

#### v3 beacon sequences

The v3 PiggyBac beacon was cloned by digesting the v2 PiggyBac beacon (Plasmid 5, Supplementary Table 6) with NheI-HF (NEB, R3131S) and EcoRI-HF (NEB, R3101S) and thereby removing the iT7-CS2-12N-CS1-T7 cassette. A replacement cassette replacing one T7 promoter with SP6 was purchased as an ssDNA oligo with 12 degenerate bases in the barcode position (oligo 8, Supplementary Table 5, IDT ultramer) and assembled by Gibson Assembly following the same protocol used for integrating the 12N barcode in v1 beacons described above. The v3 beacon is available as Plasmid 8 in **Supplementary Table 6**.

#### crRNA cloning

The crRNA acceptor backbone was ordered from Addgene (178883) and linearized using BbsI-HF (NEB, R3539S), dephosphorylated with QuickCIP (NEB, M0525S) and purified using the Zymo DNA Clean-up and Concentration kit (Zymo Research). To clone the GFP-targeting crRNA, top and bottom strand phosphorylated oligos (oligos 9 + 10) with matching overhangs were ordered (IDT) and annealed by heating at 95 °C for 5 mins followed by gradual cooling to room temperature (0.1 °C per second). 30 ng of digested backbone were combined with 1 µL of 1 µM annealed oligos, 0.25 µL BbsI-HF (NEB, R3539S), 1 µL T4 DNA ligase (NEB, M0202L), 1 µL 10x T4 DNA ligase buffer to a total volume of 10 µL and incubated at 16 °C for 30 mins followed by heat inactivation at 65 °C for 10 mins. 1 µL of the ligation product was transformed into chemically competent NEB Stable *E. coli* cells (NEB, C3040H).

All plasmids were purified using the Zymopure Plasmid Miniprep, Midiprep, or Maxiprep kits (Zymo Research, D4212/D4201/D4202) and sequence-verified by whole-plasmid sequencing (Plasmidsaurus).

### PiggyBac and Sleeping Beauty transposition

Plasmids harboring either the v1 or v2 Shred-seq beacons flanked by PiggyBac (PB) or Sleeping Beauty (SB) ITRs were cloned into a library and assessed for sufficient barcode diversity. The transposon libraries were then introduced into wild-type HAP1 cells along with a plasmid expressing either hyperactive PB transposase (Yusa et al. 2011) or 100x SB transposase (Jin et al. 2011) via nucleofection using the Lonza 4D-Nucleofector and the SF Cell Line 4D-Nucleofector X Kit XL (V4XC-2024). Per biological replicate, 1 million cells were nucleofected in a 100 µL cuvette. Prior to nucleofection, cells were trypsinized, counted, and collected by centrifugation (300g, 3 mins). The pellets were washed twice in PBS and resuspended in SF solution, supplemented 1:4.5 with Supplement 1 as well as 2.5 µg of the transposon and 0.5 µg of the transposase. The cell + DNA mixture was transferred to cuvettes, nucleofected using the EN138 program, and recovered in 10 cm dishes with pre-warmed IMDM media supplemented with FBS and Pen/Strep. The cells were expanded over 10-14 days to allow for depletion of any episomal plasmids. Depending on bottleneck size, between 400 and 50,000 GFP-positive cells (to enrich for beacon integrations) were sorted by FACS (see flow cytometry section) and expanded for 7 days before beacon integration site mapping and Cas3 deletion screening.

### Lentivirus production and infection of v2 beacons

Third-generation lentivirus was produced in HEK293T cells that were transfected with Lipofectamine 3000 (Invitrogen, L3000015). 500,000 cells were seeded in a 6-well plate one day before transfection. 1.5 μg of a lentiviral transfer plasmid, 1 μg of pMDLg/pRRE (Addgene 12251), 0.5 µg of pRSV-Rev (Addgene 12253), and 0.5 μg of pMD2.G (Addgene 12259) were mixed in 125 µL Opti-MEM (Gibco, 51985091) together with 3.5 μl P3000 reagent and incubated for 5 min at room temperature. In a separate tube, 5 μl of the Lipofectamine 3000 reagent was mixed with 125 µL of Opti-MEM. The two mixes were combined and incubated for another 30 min at room temperature. 250 µL of the transfection mix was then added to 80% confluent cells in 2 mL DMEM media in a well of a six-well plate. After 48h, the supernatant was collected and stored at -20°C. HAP1 cells were spin-infected in 6-well plates at 550g for 1 hour at 32 °C.

### Cas3 editing

Plasmids expressing Cas3 and Cascade open reading frames from the EF1 alpha promoter were ordered from Addgene (Addgene no. 178878, 178879, 178880, 178881, 178882) and mixed in the following ratio: 45 ng Cas3, 22.5 ng Cas5, 67.5 ng Cas7, 270 ng Cas8, 45 ng Cas11, and 50 ng of a puromycin encoding vector to select for successful transfection. The HAP1 cells were nucleofected using the Lonza 4D-Nucleofector and the SF Cell Line 4D-Nucleofector X Kit XL (V4XC-2024). Per biological replicate, 6 million cells were nucleofected across two 100 µL cuvettes. Prior to nucleofection, cells were trypsinized, counted, and collected by centrifugation (300g, 3 mins). The pellets were washed twice in PBS and resuspended in SF solution, supplemented 1:4.5 with Supplement 1 as well as 7.5 µg of the Cas3:Cascade mix and 756 ng of the crRNA plasmid (Plasmid 9, **Supplementary Table 6**). The crRNA plasmid was omitted for negative controls. The cells were transferred to cuvettes, nucleofected using the EN138 program, and recovered in 10 cm dishes with pre-warmed IMDM media supplemented with FBS and Pen/Strep. Puromycin (2 µg/ml) was added 24 hours after nucleofection. On day 3, cells were split. 2-3 million cells were collected for sequencing, and the remaining cells were reseeded in T225 flasks in puromycin-free media.

### T7 mapping and deletion calling for short-read sequencing

#### *In vitro* transcription

Genomic DNA (gDNA) was extracted from pellets of 2-3 million cells using the DNeasy Blood & Tissue kit (Qiagen) with the addition of 50 µg RNaseA (NEB, T3018L) at the resuspension step. Two technical replicates of *in vitro* transcription were performed per sample using the HiScribe T7 High Yield RNA Synthesis Kit (NEB, E2040). For each reaction, 1500 ng gDNA was mixed with 30 µL NTP buffer mix, 3 µL DTT, and 6 µL T7 RNA polymerase mix and incubated at 37 °C for 16 hours. Following IVT, gDNA was digested using the Turbo DNA-free kit (Invitrogen, AM1907) by adding 10 µL 10x buffer, 2 µL Turbo DNase, and 28 µL nuclease-free water. The reaction was incubated at 37 °C for 60 minutes. To inactivate the DNase, 10 µL of DNase inactivation reagent was added, incubated for 5 mins, and spun down at 10,000g for 2 mins. 85 µL of supernatant was transferred to a fresh tube. The RNA was cleaned up using 48 µL (0.8 volume) of RNA clean XP beads (Beckman, A63987).

#### Reverse transcription

The purified RNA was reverse transcribed using the Maxima H Minus RT (Thermo Fisher Scientific, EP0752). 400-1000 ng of RNA were mixed with 1 µL of 10 mM dNTP, 0.5 µL of a 100 µM random RT primer (oligo 11, Supplementary Table 5) in a total volume of 13 µl. The mix was incubated at 65 °C for 5 min and cooled on ice for another 5 min. A mix consisting of 4 µL 5x RT buffer, 1 µL Nuclease-free water, 1 µL RNase Out (Invitrogen, 10777019), and 1 µL Maxima H Minus RT was added to the sample and incubated at 23 °C for 10 mins, 42 °C for 90 min, and 85 °C for 5 min. The remaining RNA was digested by adding 1 µL of RNase Cocktail (Invitrogen, AM2286) and incubating at 37 °C for 10min. The RT-product was subsequently purified using 0.8 volumes of Ampure XP beads (Beckman).

#### PCR

Sequencing libraries were prepared using nested PCRs. For v1 and v2 beacons containing samples, three 50 µL PCR1 reactions using KAPA2G Robust HotStart ReadyMix (Roche, 08041113001) were performed for each technical replicate (resulting in 6 PCR1s per sample). For each reaction, 10% of the RT product was mixed with 25 µL of 2x KAPA2G Robust HotStart ReadyMix (Roche), 2.5 µL oligo 12, 2.5 µL oligo 13 (mapping) or oligo 14 (deletion calling), and adjusted to 50 µL with nuclease-free water. The PCR program consisted of 3 mins at 95 °C for the initial denaturation, followed by 13 cycles of 15s at 95 °C, 15s at 58 °C, 30s at 72 °C, and a final extension cycle of 1 min at 72 °C. The three PCR1 reactions were pooled and 50 µL of the pool was purified using 0.8x Ampure XP beads and eluted in 20 µl. One µL of purified PCR1 product was carried forward for 14 cycles of the indexing PCR (3 mins at 95 °C, followed by 14 cycles of 30s at 98 °C, 30s at 58 °C, 30s at 72 °C, and 1 min at 72 °C) using 1 µL of forward and reverse indexing primers (oligos 15 and 16, Supplementary Table 5) and 2x KAPA Hifi Hotstart ReadyMix MM (Roche, 07958935001) in 50 µl. The indexing PCRs were pooled and gel-purified. 40 µL of the pooled PCR product was mixed with 20 µL of water and loaded across three wells of a 1 or 2% eGel (Invitrogen) and ran for 10-20 mins. Products between 600 and 1,200 bp were excised and purified using the Zymoclean Gel DNA recovery kit (Zymo Research). The resulting libraries were sequenced on the NextSeq 2000 (Illumina) using 200 cycles P1/P2 or P3 kits. 55 cycles were allocated to read 2 (for barcode extraction), 6 or 10 cycles for index1, 10 cycles for index2, and the remaining cycles for read 1. For v3 beacons, 8 cycles of PCR1 was performed using 2x LongAmp Hotstart Taq MM (NEB; instead of 2x Kapa Robust MM). The rest of the protocol was identical to the one described above.

### T7 deletion calling for long-read sequencing

T7 deletion calling for long-read sequencing was performed as per the protocol for short-read sequencing with the following modifications. (1) LongAmp polymerase (NEB, M0533S) was used instead of Kapa Robust for PCR1 with the following cycling conditions: 3 mins at 94 °C for the initial denaturation, followed by 14 cycles of 20s at 94 °C, 30s at 55 °C, 3 min at 65 °C, and a final extension cycle of 5 min at 65 °C. (2) The indexing PCR cycling conditions were changed to: 98°C 3 min; 98 °C 30 s, 58 °C 30 s, 72 °C 3 min (14 cycles), 72 °C 3 min. (3) Pooled indexing PCR samples were cleaned up by the addition of 0.7x Ampure XP beads instead of gel extraction. (4) Libraries were sequenced using the MinION (Oxford Nanopore) (see Long-read sequencing).

### Long-range PCR

Genomic DNA was prepared as described in the “*In vitro* transcription” section. Unique molecular identifiers (UMIs) were added in a 5-cycle PCR1 reaction: 25 µL of 2x LongAmp Hot Start Taq 2X Master Mix (NEB) was mixed with 2 µL each of 10 µM primers (oligos 17 + 18, Supplementary Table 5), 600 ng of template genomic DNA, and adjusted to 50 µL with nuclease-free water and amplified with the following cycling conditions: 3 mins at 94 °C for the initial denaturation, followed by 5 cycles of 20s at 94 °C, 30s at 59 °C, 25 min at 65 °C, and a final extension cycle of 25 min at 65 °C. The PCR product was purified using 0.8x Ampure XP beads and resuspended in 20 µL of nuclease-free water. A second round of indexing PCR (oligos 15 and 16, Supplementary Table 5) was done using 10 µL of the purified first PCR product as input (2x LongAmp Hot Start Taq 2X Master Mix (NEB): 18 cycles of 20s at 94 °C, 30s at 59 °C, 25 min at 65 °C, and a final extension cycle of 25 min at 65 °C). The indexing PCR product was purified using 0.8x Ampure XP beads.

### Long-read sequencing

500 ng of pooled PCR products were taken forward for library preparation using the Ligation Sequencing Kit v14 (Oxford Nanopore Technologies, SQK-LSK114) according to the manufacturer’s instructions. The libraries were sequenced on the MinION (Oxford Nanopore Technologies) using MinION (R10.4.1) flow cells (Flo-MIN114). Dorado (v0.9.0) was used for basecalling using high-accuracy or super-high-accuracy models (v5).

### Clone isolation

Clones were isolated from bottlenecked HAP1 cells harboring Shred-seq beacons via limiting dilution. Cells were trypsinized, counted, and collected by centrifugation (300g, 3 mins). Subsequently, cells were resuspended in 3-5 mL IMDM media supplemented with FBS and Pen/Strep and serially diluted to a concentration of ∼1.5 cells/mL. This suspension was plated onto individual wells of a 96-well tissue culture plate, bringing the estimated cell count per well to ∼0.3 cells. Clones were grown for 10-14 days before expansion into larger plate formats for genotyping and Cas3 editing.

### Single-cell Shred seq

v3 Shred-seq beacon-containing HAP1 clones were isolated from 1000-cell bottlenecked HAP1 population with coordinate-mapped v3 beacons, by diluting to 0.3 cells/well and distributing across all wells of a 96-well plate. Selected clones were nucleofected with Cas3/Cascade components and the beacon-targeting crRNA-107 (Plasmid 10, **Supplementary Table 6**) and selected for 48 hours with 2 µg/ml Puromycin for successful transfection as described above. Cells with GFP-loss were isolated by FACS five days post-transfection. For each clone, 150 cells were bottlenecked and expanded for 10 days with the final split two days before harvest. On the day of harvest, cells from clone 1 and clone 2 were mixed 90:10, then fixed and processed for *in situ* transcription and single-cell capture.

#### Methanol fixation

Two million cells were harvested by trypsinization, pelleted by centrifugation (300g, 5 mins) in a pre-chilled centrifuge, washed twice with 2 mL ice-cold PBS, and resuspended in 800 µL PBS. Cells were fixed by dropwise addition of 3.2 mL ice-cold methanol (pre-chilled at -20 °C) while gently swirling the tube by hand. Fixed cells were incubated on ice for 30 min, with gentle mixing every 5 min. Cells were then rehydrated by slow addition of 8 mL cold PBS with gentle swirling, pelleted, and resuspended in 60 µL PBS for counting and downstream processing. PBS was pre-chilled to 4°C for all washing and rehydration steps.

#### T7 *in situ* transcription

Cells were adjusted to approximately 10,000 cells/µL, and 10.5 µL of cell suspension was distributed into strip tubes on ice. HiScribe T7 High Yield RNA Synthesis Kit (NEB, E2050L) was used for T7 in situ transcription reactions. 15 µL of NTP buffer mix, 1.5 µL of water, and 3 µL of T7 RNA polymerase mix were added to the cells for a final reaction volume of 30 µL. Reactions were incubated at 37°C for 30 min. Cells were counted and adjusted to 1228 cells/µl, and 23.6 µL of cells were added to a new strip tube (aiming for 20,000 output or 29,000 input cells).

#### 10x Capture, Poly(A)-tailing, reverse transcription

Single-cell Shred-seq followed the Chromium GEM-X Single Cell 3′ workflow with several modifications. To enable poly(A) tailing post emulsion, *E. coli* Poly(A) Polymerase (NEB, M0276) was directly added to the RT master mix. The RT/polyA mix consisted of 16.3 µL RT Reagent E (10x genomics), 3.1 µL of 100 µM LNA template switching oligo (oligo 20), 2 µL Reducing Agent B, RT enzyme E, *E. coli* Poly(A) Polymerase (5U/µl), and 10 µL ATP (10 mM, NEB). 41.4 µL of the RT/poly(A)-tailing mix was added to 23.6 µL of cells. The cells in the RT/poly(A)-tailing buffer were loaded directly onto one lane of the GEM-X chip and run on the Chromium X-series instrument (GEM-X 3P program). The droplets were collected and incubated according to a modified protocol of 25 min at 37 °C, 45 min at 53 °C, and 5 min at 85 °C. The 37 °C step was added to allow for polyadenylation to occur (Dinçaslan et al. 2024). Cleanup and cDNA amplification were performed following the 10x Genomics user guide. To more efficiently capture T7 IST transcripts, a small amount (0.5 µM) of a CS2-binding primer (oligo 21, Supplementary Table 5) was spiked into the cDNA amplification step.

#### T7 IST library generation

A first PCR was performed in 50 µL reactions containing LongAmp polymerase mix and oligos 21 + 22 (Supplementary Table 5), using 4 µL purified preamplified cDNA as input. Cycling was performed with the following conditions: 95 °C for 3 min; 8 cycles of 95 °C for 15 s, 59 °C for 30 s, and 72 °C for 45 s; followed by 72 °C for 2 min. PCR products were purified using a 0.8x Ampure XP bead cleanup and eluted in 20 µL water. Indexed sequencing libraries were then generated in 50 µL reactions containing KAPA HiFi mix, water, and sample-specific P5 and P7 indexing primers (oligos 23 + 24, Supplementary Table 5). Reactions were cycled as follows: 98 °C for 3 min; 14 cycles of 98 °C for 30 s, 60 °C for 30 s, and 72 °C for 30 s; followed by 72 °C for 1 min.

Indexed libraries were assessed on a 1% agarose gel, where successful reactions typically produced a smear from approximately 400 bp to 1.5 kb; adapter dimers at approximately 120-170 nt were also monitored. Libraries were pooled and resolved on a 1% agarose gel, and fragments in the ∼600 bp to 1.2 kb size range were excised and purified using a Zymo gel extraction kit. Final libraries were quantified on an Agilent TapeStation using HS5000 reagents and diluted to prepare a 2 nM pool for sequencing using the Illumina NextSeq 2000 platform (P2 kit 300 cycles at 650 pM loading concentration; R1:55, I1:10, I2:10, R2:263).

#### Transcriptome library generation

Transcriptome libraries were constructed from amplified cDNA according to the Chromium GEM-X Single Cell 3’ Reagent Kits v4 user guide. rDNA was depleted using the SEQuoia RiboDepletion Kit (Bio Rad, 17006487) following the manufacturer’s instructions and using 20 ng as input and 7 PCR cycles. The ribodepleted libraries were pooled and quantified on an Agilent TapeStation using HS1000 reagents and diluted to prepare a 2 nM pool for sequencing using the Illumina NextSeq 2000 platform (P2 kit 100 cycles at 650 pM loading concentration; R1:28, I1:10, I2:10, R2:90).

## Computational analysis

### Barcode amplicon sequencing

Genomic DNA (gDNA) was extracted from pellets of 2-3 million cells using the DNeasy Blood & Tissue kit (Qiagen) with the addition of 50 µg RNaseA (NEB) at the resuspension step. Two technical replicates were performed per sample. PCR1 was performed in a 50 µL reaction with KAPA HiFi HotStart ReadyMix MM (Roche), with 2.5 µL of each 10 µM primer (oligos 14 + 19, Supplementary Table 5) and ∼250 ng gDNA template at the following cycling parameters: 98 °C - 3 min; 23 cycles of 98 °C - 20s, 60 °C - 30s, 72 °C - 45s; 72 °C - 2 min. PCR1 was cleaned up using AmpureXP beads (1.2x) and eluted in 20 µL of H_2_O. PCR2 (sample indexing and sequencing adapter addition) was again performed with KAPA HiFi HotStart ReadyMix MM (Roche) in a 50 µL reaction using 1 µL of eluate from the previous step as template and 1 µL of each 10 µM sample indexing primer (oligos 15 + 16, Supplementary Table 5). PCR2 was run at the following cycling parameters: 98 °C - 3min; 8 cycles of 98 °C - 20s, 60 °C - 30s, 72 °C - 45s; 72 °C - 2 min. PCR2 reactions were pooled and cleaned up with AmpureXP beads (1x) and eluted in 20 µL of H_2_O. Sample quality was confirmed on a TapeStation High Sensitivity D1000 ScreenTape (Agilent), after which sequencing was performed on an Illumina NextSeq2000 using 200 cycles P1/P2 or P3 kits with the following read lengths: 155 read1, 8 index1, 8 index2, and 45 on read2.

### Beacon integration site mapping

A modified version of the pipeline presented in (X. Li et al. 2024) was used. Raw paired-end FASTQ files were processed using a sequential filtering, annotation, alignment, and breakpoint-calling workflow (mapping_pipeline.sh). Read 2 captures a PCR unique molecular identifier (UMI) and the T7 transcript start, containing a 10X Capture Sequence 2 (CS2), a unique beacon barcode, and a 10X Capture Sequence 1 (CS1), while Read 1 captures genomic context and is used for integration site mapping. First UMIs were extracted from the first 10 bases of read 2 and incorporated into the read name using fastp (Chen et al. 2018). Reads failing a minimum average base quality threshold (Q15) were removed, and poly-G artifacts were trimmed. Next, reads were filtered for the presence of a constant sequence on the beacon (CS1: TTGCTAGGACCGGCCTTAAAGC) using cutadapt (Martin 2011). Beacon barcodes were then captured and appended to the read name using fastp. Read pairs were further filtered to retain the correct context (GCTCACCTATT) following the beacon barcode on read 2. Finally, only read 1 sequences that mapped into the ITRs/LTRs were retained (e.g. CCCTAGAAAGATAATCATAT for PiggyBac). The filtered reads were then aligned to the hg38 genome using BWA-MEM (Li and Durbin 2009), and alignments were sorted with samtools (Li et al. 2009). UMIs were deduplicated using UMI-tools (Smith et al. 2017). Genomic alignment coordinates were subsequently exported to BED format using sam2bed. Coordinates for read groups were collapsed using a custom python script (collapse_barcodes.py) to obtain candidate integration site coordinates.

The candidate integration sites were further consolidated and filtered (integration_filtering.R). To account for small differences in alignment position around the same integration site, alignments associated with the same barcode, chromosome, and strand were grouped when their genomic coordinates fell within 500 bp of one another. Within each group, the genomic coordinate with the greatest UMI support was retained, and UMI counts across the cluster were summed. Barcode sequences differing by a Hamming distance of ≤1 at the same candidate integration were consolidated. Because sequencing depth differed among experiments, the minimum UMI-support threshold was adjusted by sample, ranging from >5 to >20 UMIs for the large-scale datasets used in Figure 2. For PiggyBac and lentiviral beacon libraries, uniquely mapped integration sites were required to account for >90% of the mapping signal for the corresponding barcode and for barcode-specific UMIs to comprise >80% of reads at the assigned locus. For Sleeping Beauty integrations, which showed a greater frequency of multiple mappings per barcode, the unique-mapping fraction threshold was relaxed to >50%, while retaining the >80% locus-support threshold. The resulting high-confidence beacon integration maps were used to associate subsequent Cas3-induced deletion junctions with their corresponding genomic launchpads. The custom mapping_pipeline.sh, collapse_barcodes.py, and integration_filtering.R scripts used for integration site mapping are available at https://github.com/pinglaylab/deletion_scanning.

### Chromatin states and epigenetic analyses

The following publicly available datasets were collected for HAP1 cells: DNase-seq ENCFF162WTC, H3K4me1-ChIP-seq ENCFF639UYT, H3K36me3-ChIP-seq ENCFF216JJJ, H3K9me3-ChIP-seq GSM6165952, H3K27me3-ChIP-seq GSM2897158, H3K4me3-ChIP-seq ENCFF461TZF, and H3K27ac-ChIP-seq ENCFF742SZS (ENCODE Project Consortium 2012; Zhang et al. 2020; Schick et al. 2019). For data sets from ENCODE, hg38-aligned bam files were downloaded, and for data sets from the sequence read archive, fastq files were downloaded and aligned to the hg38 genome build using bwa-mem. The resulting bam files were binarized using ChromHMM (Ernst and Kellis 2012), and a 15-state model was learned. The trained ChromHMM model and state annotations are available at https://github.com/pinglaylab/deletion_scanning.

### Deletion calling

#### From short-read T7 mapping data

Raw paired-end FASTQ files were processed using a sequential filtering, annotation, alignment, and breakpoint-calling workflow (deletion_pipeline.sh, **Figure S2**). First UMIs were extracted from the first 10 bases of read 2 and incorporated into the read name using fastp. Reads failing a minimum average base quality threshold (Q15) were removed, and poly-G artifacts were trimmed. Next, reads were filtered for the presence of a constant sequence on the beacon (CS2: CCTTAGCCGCTAATAGGTGAGC) using cutadapt. Beacon barcodes were then captured and appended to the read name using fastp. Read pairs were further filtered to retain the correct context (GCTTTAAGGCC) following the beacon barcode. Reference genomes were prepared for each beacon construct by appending the beacon sequence as an additional chromosome to the hg38 build of the human genome. Filtered reads (R1) were aligned to the appropriate reference using BWA-MEM, and alignments were sorted and indexed with samtools to generate coordinate-sorted BAM files. Genomic alignment coordinates were subsequently exported to BED format using sam2bed. In downstream analysis (R), BED records were parsed to recover UMI and barcode identifiers from read names, and putative deletions were summarized by counting distinct UMIs per barcode and alignment coordinate. The T7 molecule table was then mapped to beacon integration sites by fuzzy matching barcodes with up to one mismatch (Hamming distance ≤1). Deletions were identified by filtering reads where the genomic alignment was on the same chromosome as the beacon and where the mapping strand and beacon orientation were consistent with Cas3-induced deletions. Mapped deletions were further filtered to retain events within 1000-500,000 bp of a mapped beacon barcode. Finally, unique lists of deletions were generated by merging putative deletion calls whose breakends fall within 300 bp of one another (random N priming in the reverse transcription will create molecules of varying length from transcript originating at the same beacon and consequently different alignments from the same breakpoint. We allow 300 bp to account for that). The custom deletion_pipeline.sh and deletion_filtering_pipeline.R scripts used for deletion calling are available at https://github.com/pinglaylab/deletion_scanning.

#### Split read-based deletion calling

A split-read based deletion calling approach was used for long-read data and for the analysis of deletion initiation and breakend repair. Raw FASTQ files were first processed and aligned identically to the short-read workflow to extract UMIs and beacon barcodes and to retain only reads with the expected beacon context. To identify deletions, a custom Python script (find_deletions.py) was used that searched for reads with one segment aligning to the beacon reference chromosome and a second segment aligning elsewhere in the genome. For each such split read, the breakpoint on the beacon side was defined as the end coordinate of the alignment to the reference chromosome, and the genomic breakpoint was defined from the distal alignment according to its strand orientation. The script outputs a table of candidate deletion-supporting reads together with the associated barcode and a separate BAM file with all deletion-supporting alignments. Deletion candidates were further filtered and mapped to beacon sites identically to the short-read workflow, with an additional filter based on the deletion start site on the beacon to distinguish deletions from long sequencing reads that map into genomic context downstream of an intact beacon. The custom longread_deletion_pipeline.sh and deletion_filtering_pipeline.R scripts are available at https://github.com/pinglaylab/deletion_scanning

#### From long-range amplicon sequencing

Sequencing reads were first enriched *in silico* for molecules containing the expected forward and reverse primer sequences using seqkit amplicon (Shen et al. 2016) and cutadapt, allowing up to three mismatches. Unique molecular identifiers (UMIs) were extracted and appended to the read name using fastp. Processed reads were aligned to a custom hg38 reference genome, with the beacon construct appended as an additional chromosome, using minimap2 (Li 2018) with HiFi settings and allowing shorter alignment seeds (-s40). Alignments were sorted and indexed with samtools. Candidate deletion-supporting reads were identified from the aligned BAM file using a custom split-read parsing script analogous to that used for long-read T7 mapping. Candidate deletions were first restricted to the expected chromosome and orientation. UMIs were collapsed using Hamming distance ≤1, and consensus breakpoint coordinates were assigned. Nearby breakpoint calls were then merged by clustering breakpoints that were within 20 bp on the beacon side or within 100 bp on the genome side. Supporting UMIs were summed for merged deletions calls. Deletions with 10 or less UMIs were discarded. The custom longrange_PCR_deletion_pipeline.sh, find_deletions_delamp.py, and longrange_PCR_deletion_filtering.R scripts are available at https://github.com/pinglaylab/deletion_scanning.

### Genome coverage estimation

The fraction of the hg38 reference genome covered by independent deletion events was calculated at increasing coverage depths. Per-base deletion depth was computed as the number of independent deletions overlapping each genomic position. Ten random deletion sets were generated that preserved the observed number and length distribution of deletions but placed deletions uniformly across the hg38 genome, with chromosome choice weighted by chromosome length. Simulated deletions were constrained to remain within chromosome boundaries. Genome coverage was then recalculated for each simulated set using the same procedure as for the observed Shred-seq deletions.

### Modeling deletion lengths

Cas3 pre-selection deletion-length distributions were modeled using maximum-likelihood estimation. Three parametric candidate distributions were evaluated for deletions > 1000 bp: exponential (rate parameter lambda), log-normal (mu, sigma on the log scale), and Weibull (shape k, scale lambda). Model fit was compared using log-likelihood and Akaike information criterion (AIC), computed from the fitted parameter values. The distribution with the lowest AIC was interpreted as providing the best relative fit among the tested families.

### Breakend repair characterization

Only reads that spanned a deletion breakpoint were used for this analysis whereas all reads mapping to the genome were used for the short-read deletion calling pipeline. Sequencing reads were processed using a custom R pipeline (breakend_processing.R; utilizing *Rsamtools*, *GenomicAlignments*, and *Biostrings*) (Lawrence et al. 2013) to characterize repair outcomes at integration sites. The following logic was applied: A sequencing read indicative of a deletion will have a primary alignment with a soft-clipped sequence and a supplementary alignment. Computationally, primary and supplementary alignments were extracted from BAM files using scanBam (R samtools), and alignment intervals were reconstructed from CIGAR strings to extract the soft-clipped sequences. The junction type can then be inferred by comparing the soft-clipped sequence to the supplementary alignment. A supplementary alignment shorter than the soft-clipped sequence with leftover nucleotides at the breakend indicated the insertion of extra nucleotides, often a hallmark of NHEJ. Identical soft-clipped and supplementary alignment sequences indicate a junction without non-genomic nucleotides. Finally, supplementary alignments exceeding the soft-clipped sequence indicate sequence overlap between genomic and reporter alignments (i.e., homology). Individual reads were parsed by this logic and subsequently clustered if their endpoints fell within a 200 bp window. Consensus repair characteristics for each cluster were determined using the statistical mode of the constituent reads. The custom breakend_processing.R script is available at https://github.com/pinglaylab/deletion_scanning.

### SV feature annotation

A set of genome annotation tracks was assembled to quantify chromatin state, three-dimensional genome organization, replication timing, nuclear lamina association, mutational constraint, gene architecture, gene biotype, and other sequence properties (**Supplementary Table 2**). For features spanning broad genomic domains, including replication timing, lamina-associated domains, A/B compartments, and chromatin states, each deletion was annotated by the fraction of its length overlapping the corresponding feature. For all other annotations, overlap was treated as binary: deletions with any overlap were assigned a value of 1, and deletions with no overlap were assigned a value of 0 (**Supplementary Table 2**). TAD boundary intervals and lamina-associated domains were imported from BED-format resources, and mutational constraint windows were derived from a genome-wide constraint z-score track by thresholding at z > 3. A/B compartment annotations were obtained by importing a Hi-C compartment eigenvector bigWig track and assigning bins with positive values to the A compartment and bins with negative values to the B compartment. Replication timing values were discretized into earlier- and later-replicating segments according to the sign of the replication timing score. Chromatin states were determined by ChromHMM (see Chromatin states and epigenetic analysis). An Appris-filtered GTF table (Rodriguez et al. 2018) was used to define gene-structure segments and gene biotypes. Sequence-level features, including phyloP conservation and GC content, were computed per deletion. Protein and lncRNA essentialities were derived from CRISPR screens in HAP1 or across thousands of cancer cell lines (DepMap) (Arafeh et al. 2025). Each deletion was annotated with the score of the most essential gene (only considering exons) that it overlapped. The custom feature_annotation.R script is available at https://github.com/pinglaylab/deletion_scanning.

### Association of SV features with survival

Single-feature associations with post-selection survival were quantified by logistic regression. Selection status was encoded as a binary response indicating whether a deletion was observed post-selection. Each feature was tested in a separate model in which the feature value was standardized to a z-score within the corresponding ploidy subset and was included as a predictor together with the Shred-seq experiment batch as a covariate. Multiple testing across features was controlled using Benjamini-Hochberg false discovery rate correction (Benjamini and Hochberg 1995). Analyses were performed separately for haploid and diploid deletions. To assess whether associations of other genomic features persisted after accounting for coding-gene essentiality, additional models were fit either for a subset of deletions that did not overlap protein-coding genes or in which coding-gene dispensability from CRISPR screening was included as an additional covariate.

A multivariate model integrating all features was trained using elastic-net regularized logistic regression. Regularization was implemented with glmnet (Friedman et al. 2010) using a binomial family and an elastic-net mixing parameter of α = 0.5 to improve stability in the presence of correlated predictors. Feature standardization was performed within glmnet by enabling standardization. The regularization parameter λ was selected by 10-fold cross-validation with cv.glmnet, and the λ.1se solution was used to obtain a parsimonious model whose cross-validated error was within one standard error of the minimum.

### Quantifying deviations from log-normal deletion distributions

Beacons with more than ten deletions in the pre- and post-selection data were considered. Plots showing all considered deletion profiles are available at https://github.com/pinglaylab/deletion_scanning. The lengths were analyzed on the log scale for all statistical tests, and all P-values were adjusted across beacons using the Benjamini-Hochberg procedure. Global estimates of the mean and standard deviation for Cas3-induced deletions were derived from the fitted log-normal distributions on pre- and post-selection deletions (see *Modeling deletion lengths* section). To identify beacons with deletions significantly shorter than expected by the log-normal model, one-sided, one-sample t-tests comparing the mean log deletion length at each beacon to the global fitted mean were performed. Beacons with FDR < 10% were considered “selected against”. To identify beacons whose deletion profiles remained consistent with the global log-normal mean, two one-sided equivalence tests were performed on the mean log deletion length. Equivalence was tested using bounds of ±0.5σ and ±1σ around the fitted global mean, where σ is the fitted standard deviation of log deletion lengths. For each equivalence test, the larger of the two one-sided P-values was used as the TOST P-value (Schuirmann 1987). Beacons with FDR < 10% were classified as “Dispensable ±0.5 SD / ±1 SD”. Beacons whose pre-selection deletion lengths were already significantly shorter than the fitted mean were discarded.

### Single-cell analysis

#### Running Cell Ranger

To quantify beacon-derived GFP transcripts alongside the cellular transcriptome, a custom Cell Ranger reference was generated by appending a GFP contig to the Ensembl GRCh38 reference (release 115). The primary assembly FASTA and corresponding gene annotation GTF were downloaded from Ensembl. The GTF was filtered with cellranger mkgtf to retain protein-coding genes only. A GFP contig and a corresponding GFP gene model were then appended to the reference genome and GTF. The custom reference was built with cellranger mkref (v8.0.0). Transcriptome library FASTQ files were processed with cellranger count (v8.0.0) against the custom hg38+GFP reference.

#### Transcriptome library analysis

Filtered feature-barcode matrices generated by Cell Ranger were imported into R and analyzed using Seurat (Hao et al. 2021). Standard quality-control metrics were calculated for each cell, including the total UMI count, the number of detected genes, and the fraction of reads mapping to mitochondrial genes. Cells were retained if they had 400-3,000 detected genes, 500-4,000 UMIs, and ≤5% mitochondrial RNA. Gene expression counts for filtered cells were log-normalized (NormalizeData, Seurat). Normalized expression values were then scaled while regressing out total transcriptome UMI count (ScaleData, Seurat). Principal component analysis was performed on the rescaled data. Based on elbow plots, the first 10 principal components were used for neighborhood graph construction, clustering, and UMAP visualization. The set of 13,944 filtered cell barcodes was used for downstream analysis. The custom Seurat.R script used for this analysis is available at https://github.com/pinglaylab/deletion_scanning.

#### IST library preprocessing

(1) Fastq filtering and preprocessing: Single-cell T7 IST libraries were processed to recover the genomic sequence adjacent to each T7-derived molecule together with the associated cell barcode, molecular UMI, and beacon barcode (Schematic of read structure depicted in **Figure 5A**). First, the CS2 sequence was detected in read 2 and removed using cutadapt. The following 12 nt beacon barcode was extracted using fastp and appended to the read name. Next, the 16 nt 10x cell barcode was extracted from the start of read 1 and appended to the read name. Finally, the 12 nt 10x UMI was extracted from the updated read 1 sequence and also appended to the read name. Next, the poly(A) sequence was trimmed from read 2, and remaining sequences shorter than 20 nt after trimming were discarded. (2) Genome alignment and deletion calling: The processed fastq files were aligned to a custom reference containing the human genome and an additional chromosome with the v3 beacon sequence using BWA-MEM. Alignments were sorted and indexed with samtools and converted to the BED format using sam2bed. Finally, UMIs were tallied for each beacon barcode, cell barcode, and genomic alignment position to generate an IST molecule table. This table was restricted to the 13,944 cell barcodes corresponding to high-confidence transcriptomes and matched to the known clone 1 and clone 2 beacon barcodes, allowing a Hamming distance of 1. The custom single_cell_IST_pipeline.sh and T7_IST_processing.R scripts used for this analysis are available at https://github.com/pinglaylab/deletion_scanning.

#### Clone assignment from T7 IST barcode counts

Because each clone carried a known set of mapped Shred-seq beacon barcodes, clone identity was inferred from the distribution of barcode-supporting T7 IST UMIs in each cell. Cells with less than 100 total T7 UMIs were excluded, and the remaining cells were assigned to clones if they met the following purity thresholds: 0.95 for Clone 1, and 0.7 for Clone 2.

#### Deletion calling from T7 IST data

Deletions were identified from the T7 IST molecule table by applying the same positional and orientation filters used for bulk short-read deletion calling. Briefly, candidate deletion-supporting molecules were required to contain a beacon barcode matched to a mapped beacon, a genomic alignment on the same chromosome as that beacon, and an alignment orientation consistent with the direction of Cas3-induced deletion from the beacon. Candidate breakends were further restricted to positions 100-500,000 bp from the mapped beacon integration site. Putative deletion calls with breakends within 300 bp of one another were merged. To associate deletions to individual cells, UMIs were counted for each deletion-cell pair. High-confidence deletion assignments required more than 10 supporting UMIs for a given deletion in a given cell. In addition, the clone identity inferred from the deletion junction was required to match the clone identity inferred independently from beacon barcode counts.

#### Associating deletions with gene expression changes

The gene expression profiles for individual cells were annotated with high-confidence deletion and clone calls. Only cells with confident clone assignment were considered for further analysis. Gene expression was quantified using Seurat log-normalized expression values. Expression of target genes across different deletions was compared using two-tailed Wilcoxon rank-sum tests. P-values were adjusted for multiple comparisons using the Benjamini-Hochberg procedure, where multiple deletions were tested for the same gene. For genome-wide differential expression analyses, cells carrying deletions that overlapped a given gene were compared with cells carrying deletions from the same locus that spared that gene. Differential expression was calculated using the FindMarkers function in Seurat with Wilcoxon rank-sum testing on log-normalized expression values. Genes were ranked by adjusted P-value and effect size. Genome-wide P-values were corrected for multiple testing using the Benjamini-Hochberg procedure.

## Software

Genomic software: bedops (2.4.41), bedtools (2.31.1), bwa (0.7.17), cellranger (8.0.0), cutadapt (4.6), dorado (0.9.0), fastp (0.24.0), minimap2 (2.26), python (3.12.1), R (4.4.1), samtools (1.19), seqkit (2.9.0), umi_tools (1.1.6)

R packages: BiocManager (1.30.25), Biostrings (2.72.1), BSgenome.Hsapiens.UCSC.hg38 (1.4.5), eulerr (7.0.2), fuzzyjoin (0.1.6), GenomicAlignments (1.40.0), GenomicRanges (1.56.2), igraph (2.1.4), Rsamtools (2.20.0), rtracklayer (1.64.0), Seurat (5.4.0), spgs (1.0-4), stringdist (0.9.15), tidyverse (2.0.0), TOSTER (0.8.4)

Python packages: pandas (2.2.2), pysam (0.22.1)

## Data availability

Processed data are deposited alongside this manuscript as Data S1: Annotated deletions, Data S2: Annotated transcriptomes. Scripts used to analyze the data, additional necessary input files, and intermediate data files to run the scripts are deposited on https://github.com/pinglaylab/deletion_scanning. Raw sequencing data have been deposited in the Sequence Read Archive as bioproject PRJNA1469634 (https://www.ncbi.nlm.nih.gov/sra).

## Supplementary Figures

**Figure S1.**
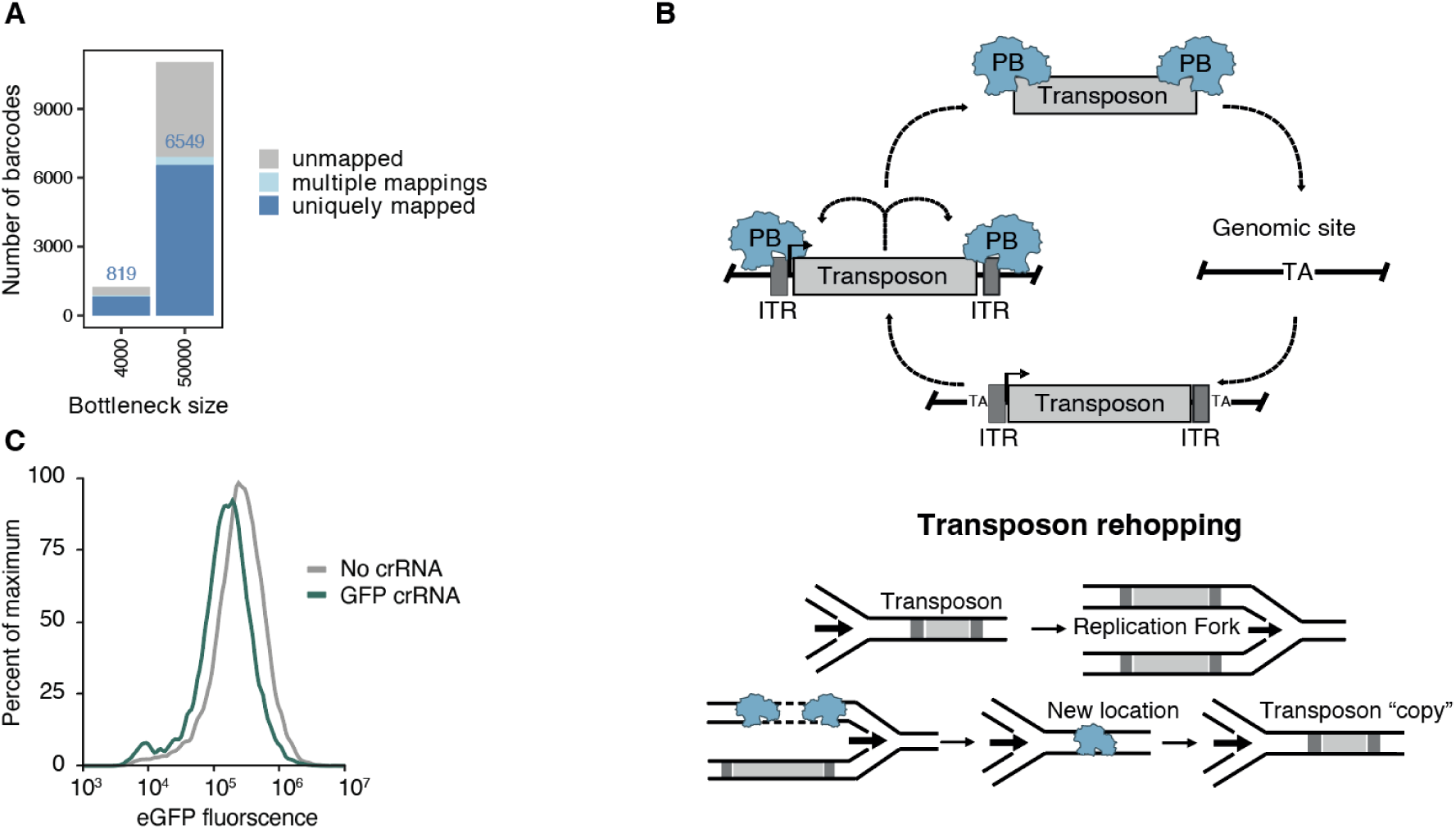
Shred-seq pilot. (**A**) Number of barcodes recovered (y-axis) after bottlenecking Shred-seq pilot populations at two different bottleneck sizes (x-axis). Stacked bars indicate barcodes that could be mapped uniquely, barcodes mapping to multiple loci, and unmapped barcodes. (**B**) Schematic illustrating PiggyBac transposon re-hopping, which can generate multiple genomic locations associated with the same beacon barcode, potentially during DNA replication. (**C**) eGFP fluorescence distributions for cells transfected with a polycistronic Cas3-Cascade construct and either a GFP-targeting crRNA or no crRNA control seven days after Cas3 nucleofection.

**Figure S2.**
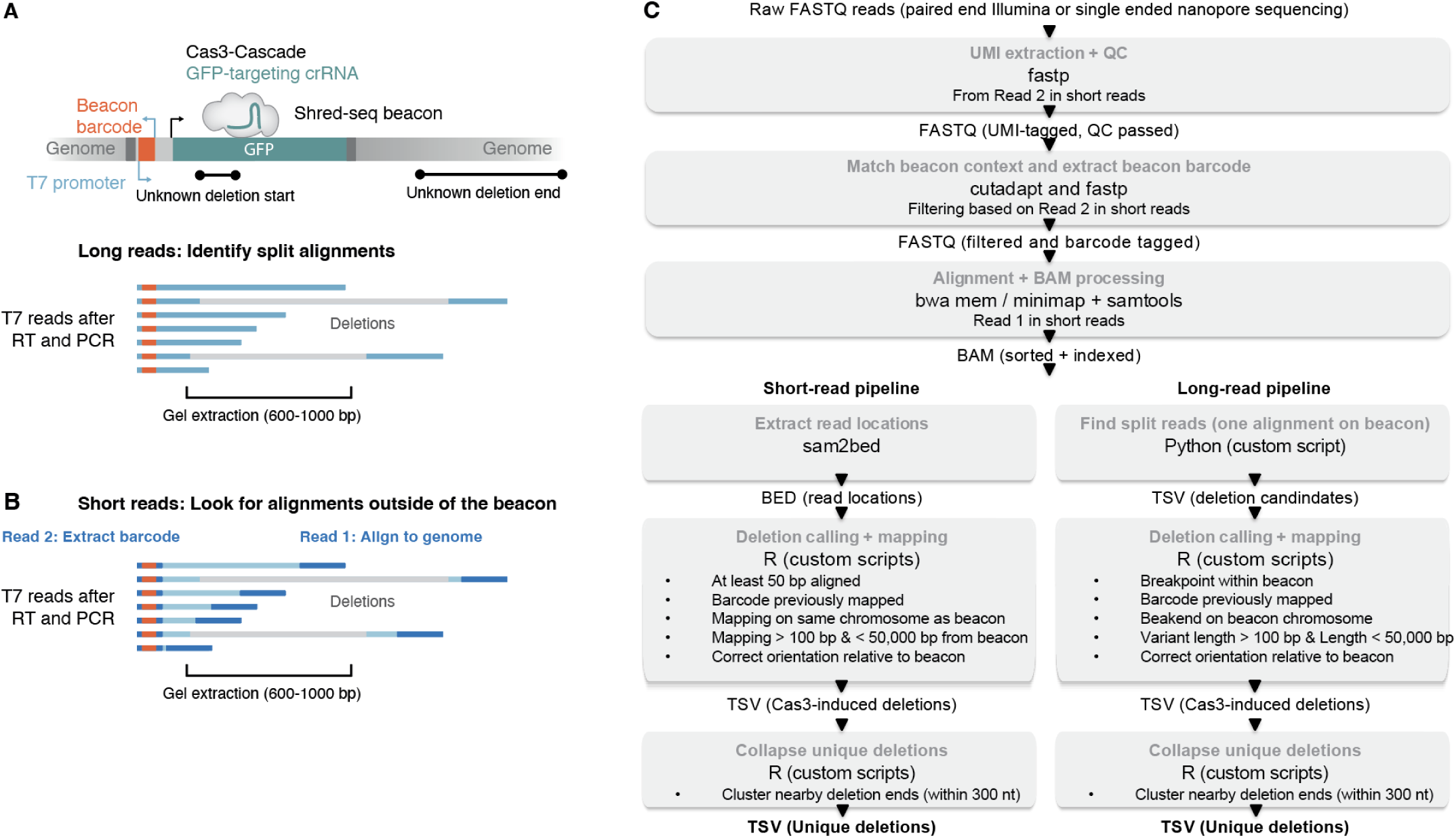
T7 deletion mapping pipeline. (**A**) Schematic of long-read deletion mapping. Following T7 *in vitro* transcription, reverse transcription, and PCR, long-read sequencing captures individual molecules spanning the beacon barcode and deletion junction, allowing direct identification of split alignments and deletion breakpoints. (**B**) Schematic of short-read deletion mapping. Read 2 captures the beacon barcode, whereas Read 1 captures adjacent genomic sequence; deletion-supporting reads are identified as barcode-associated reads aligning outside the beacon sequence. (**C**) Overview of the computational pipeline used for deletion calls from raw sequencing data. FASTQ files were processed by quality filtering, barcode extraction, alignment, deletion calling, mapping to previously established beacon integration sites, and collapsing of nearby events into unique deletions for both short-read and long-read workflows.

**Figure S3.**
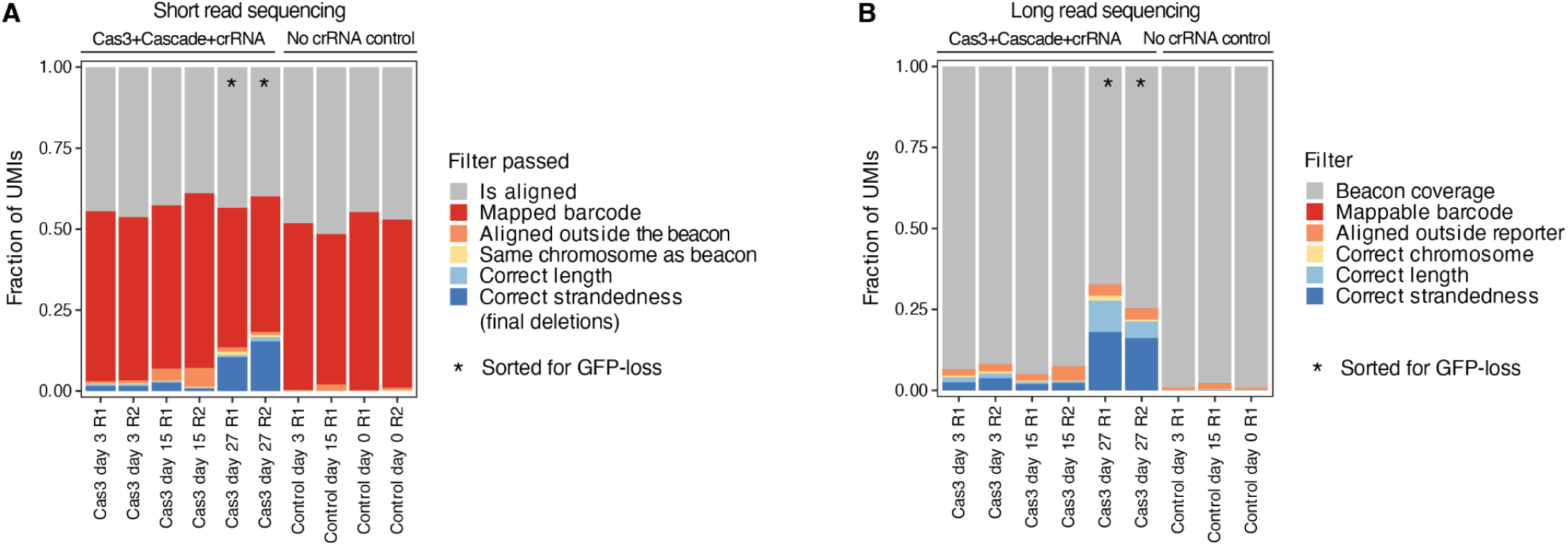
High sensitivity deletion calling. (**A**) Fraction of UMIs retained after successive filtering steps (y-axis) in the short-read pipeline across Cas3-treated and no-crRNA control samples (x-axis). Colored segments indicate the fraction of UMIs passing each criterion. Asterisks denote samples enriched for GFP loss. (**B**) As in panel **A**, but for long-read sequencing data. R1, R2 indicate replicate 1 and replicate 2.

**Figure S4.**
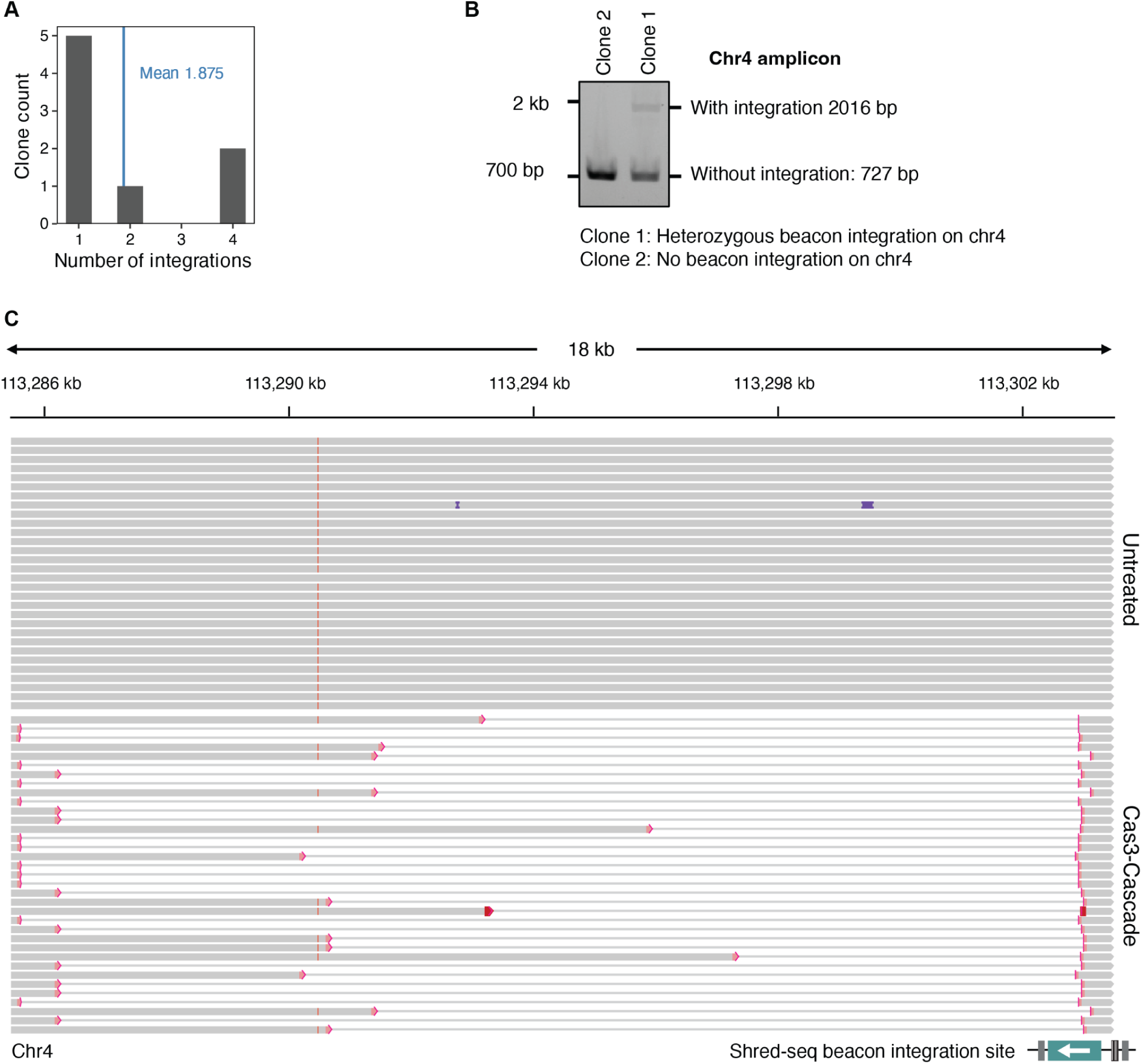
Shred-seq in individual clones. (**A**) Distribution of the number of beacon integrations detected per isolated clone from the pilot PiggyBac population. The vertical line indicates the mean number of integrations per clone. (**B**) PCR genotyping of individual clones with a candidate beacon integration on chromosome 4. Band sizes for alleles with and without integration are indicated. (**C**) Comparison of long-range amplicon sequencing profiles for cells from a clone with a beacon integration site on chromosome 4 that were left untreated (top) or transfected with Cas3, Cascade, and a GFP-targeting crRNA (bottom). Gray bars represent aligned sequencing reads and thin lines represent deletions. The Shred-seq beacon integration location is indicated at the bottom.

**Figure S5.**
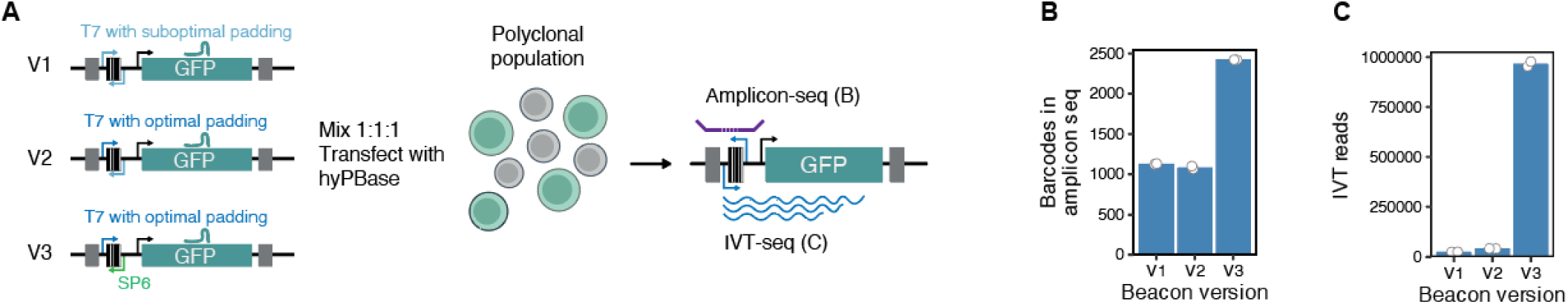
Improving T7 readout. (**A**) Schematic of Shred-seq beacon versions v1, v2, and v3 used to optimize T7-based readout. v2 includes improved T7 promoter padding, whereas v3 additionally replaces one T7 promoter with SP6 to reduce transcriptional interference. Mixed beacon populations were generated and analyzed by amplicon sequencing and IVT-seq. (**B**) Number of barcodes detected by amplicon sequencing (y-axis) for each beacon version (x-axis). (**C**) T7-derived IVT reads recovered from each beacon version.

**Figure S6.**
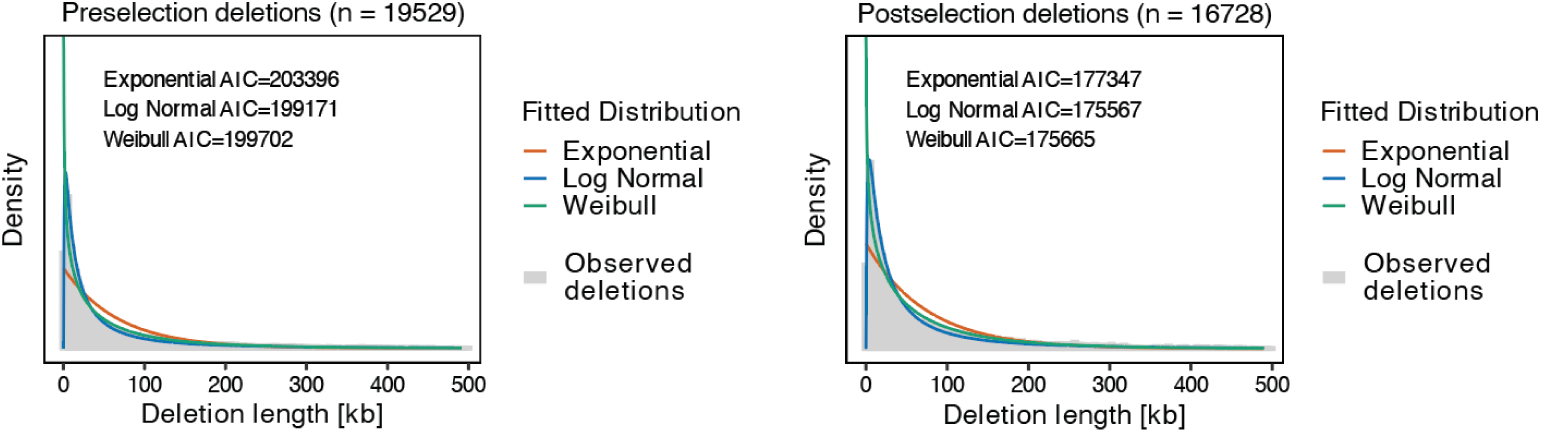
Cas3 deletions follow a log-normal distribution. Observed deletion length distributions (gray histogram) for pre-selection deletions (left) and post-selection deletions (right), together with fitted exponential, log-normal, and Weibull distributions (colored lines). Gray histograms indicate observed deletion lengths from a union of all pre-selection experiments. Insets report the Akaike information criterion (AIC) for each fit.

**Figure S7.**
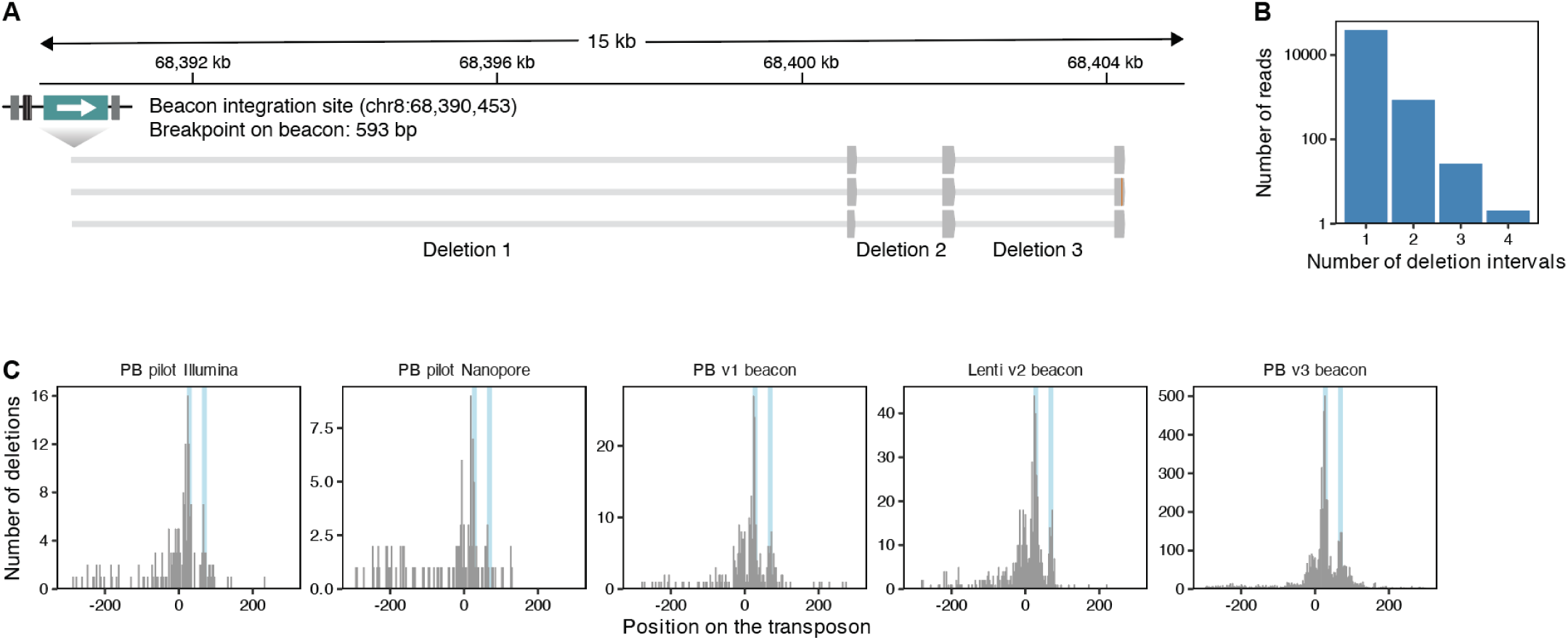
Patterns of Cas3 deletion initiation. (**A**) Example of sequencing reads with evidence for multiple successive deletions originating from a single beacon integration site. Gray bars represent aligned sequencing reads and thin lines represent deletions. (**B**) Number of sequencing reads (y-axis, log-scale) across a different number of deletion intervals (x-axis). (**C**) Histograms showing the position of deletion initiation relative to the beacon/protospacer across different experiments, beacon versions, delivery modalities, and sequencing strategies (panels). Vertical lines mark two regions with high deletion initiation.

**Figure S8.**
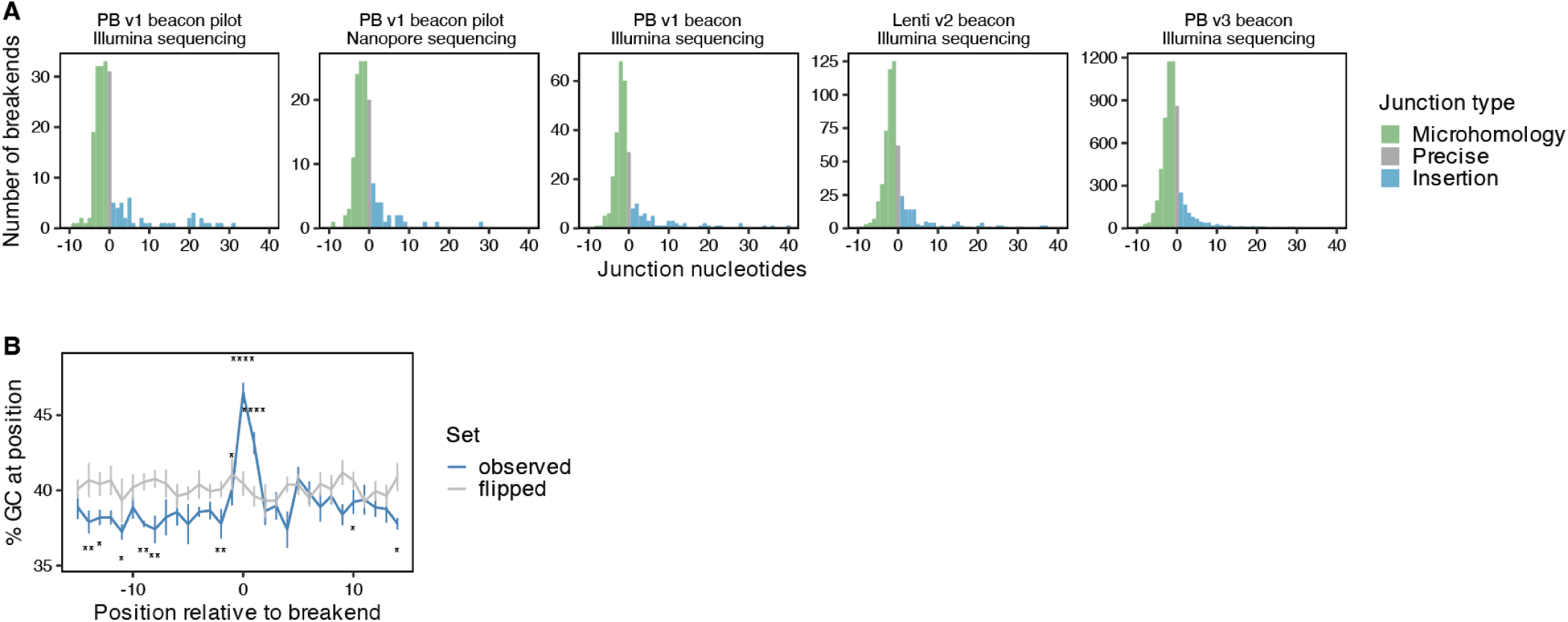
Cas3 junctions are repaired by microhomology-mediated end joining. (**A**) Distribution of junction nucleotide lengths (x-axis) across different experiments and sequencing strategies (panes). Negative values indicate overlapping sequence at the junction (microhomology), zero indicates a precise junction, and positive values indicate inserted nucleotides. Bars are colored by inferred repair class. (**B**) Average GC content (y-axis) across a 30 bp window around the deletion breakend (x-axis) for observed deletions and flipped controls (lines and colors). Whiskers show the standard error of the mean. P-values were computed with a two-sided test for equality of proportions and corrected for multiple testing across positions (Benjamini-Hochberg).

**Figure S9.**
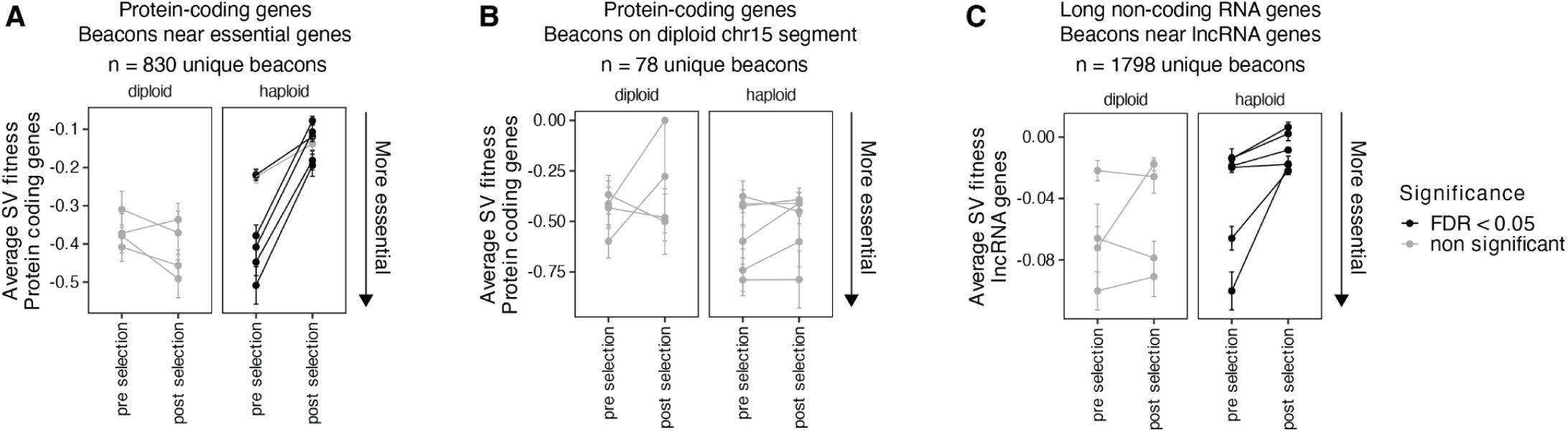
Essential protein-coding and long non-coding RNA genes are depleted from post-selection deletions. (**A**) The average protein-coding fitness of deletions (y-axis) originating from a subset of beacons within 500 kb of a core-essential gene (mean Chronos score of 1000+ dependency map cell lines < 0.5) across Shred-seq experiments and biological replicates (points and lines) pre and post selection (x-axis) separated by ploidy (panels) and shaded by FDR adjusted for multiple hypothesis testing. Whiskers represent the standard error of mean. (**B**) As in panel **A** but for a subset of beacons on a 30 Mb interval on chromosome 15 that is diploid in HAP1 cells (chr15:60812801-89346769). (**C**) As in panel **A** but for long-non coding RNA fitness and a subset of beacons with expressed lncRNAs (TPM > 1) downstream.

**Figure S10.**
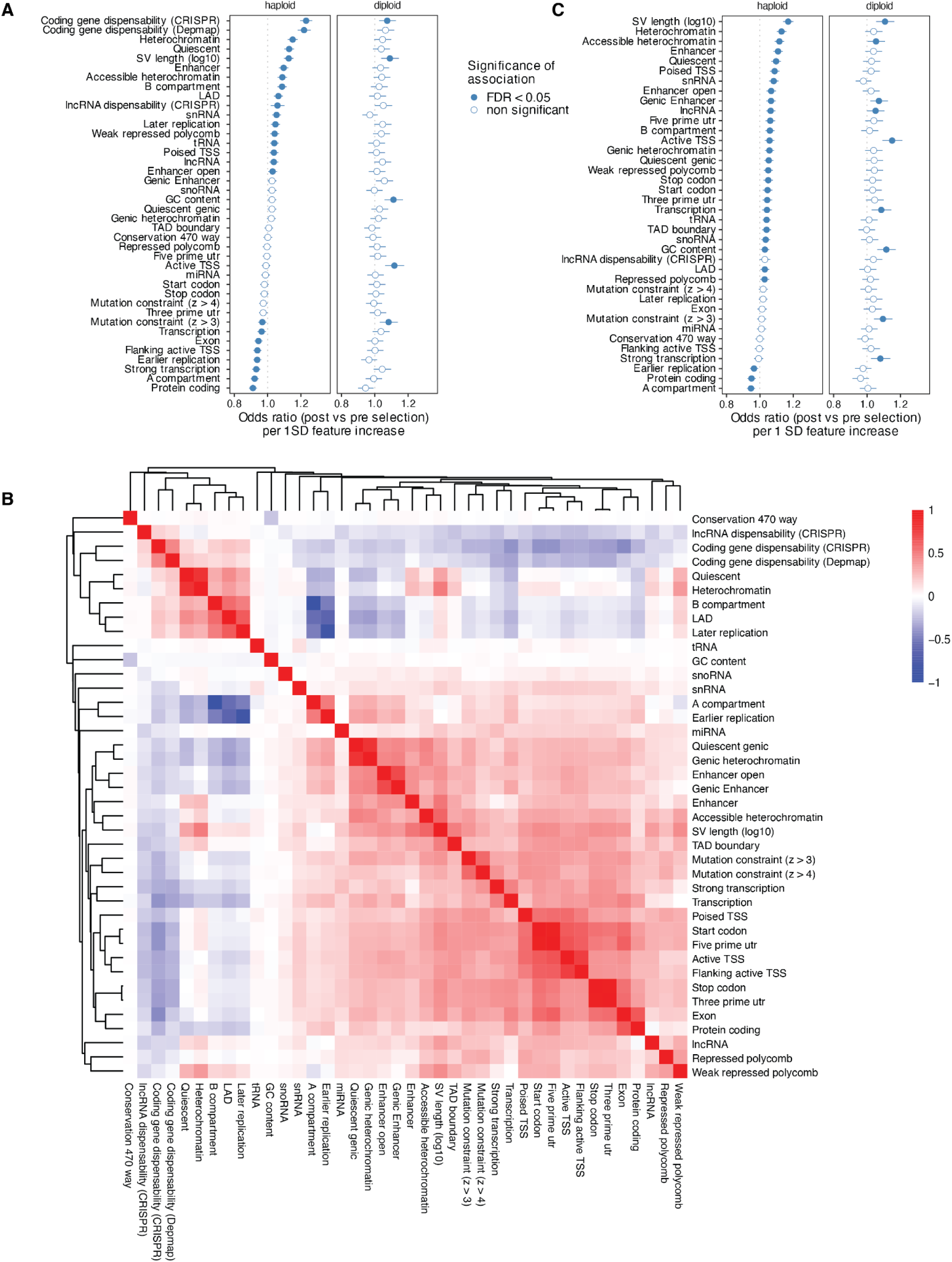
A combination of features shapes SV selection. (**A**) Odds ratios per standard deviation increase (x-axis) in features (y-axis) between post vs pre-selection variants in haploid or diploid HAP1 cells (panels). Whiskers indicate 95% confidence intervals. (**B**) Heatmap of pairwise Spearman rank correlations between genomic features in pre-selection haploid deletions. Correlations were calculated using pairwise complete observations, and colors denote the Spearman correlation coefficient (ρ). (**C**) As panel **A** but including protein-coding gene essentiality as a covariate.

**Figure S11.**
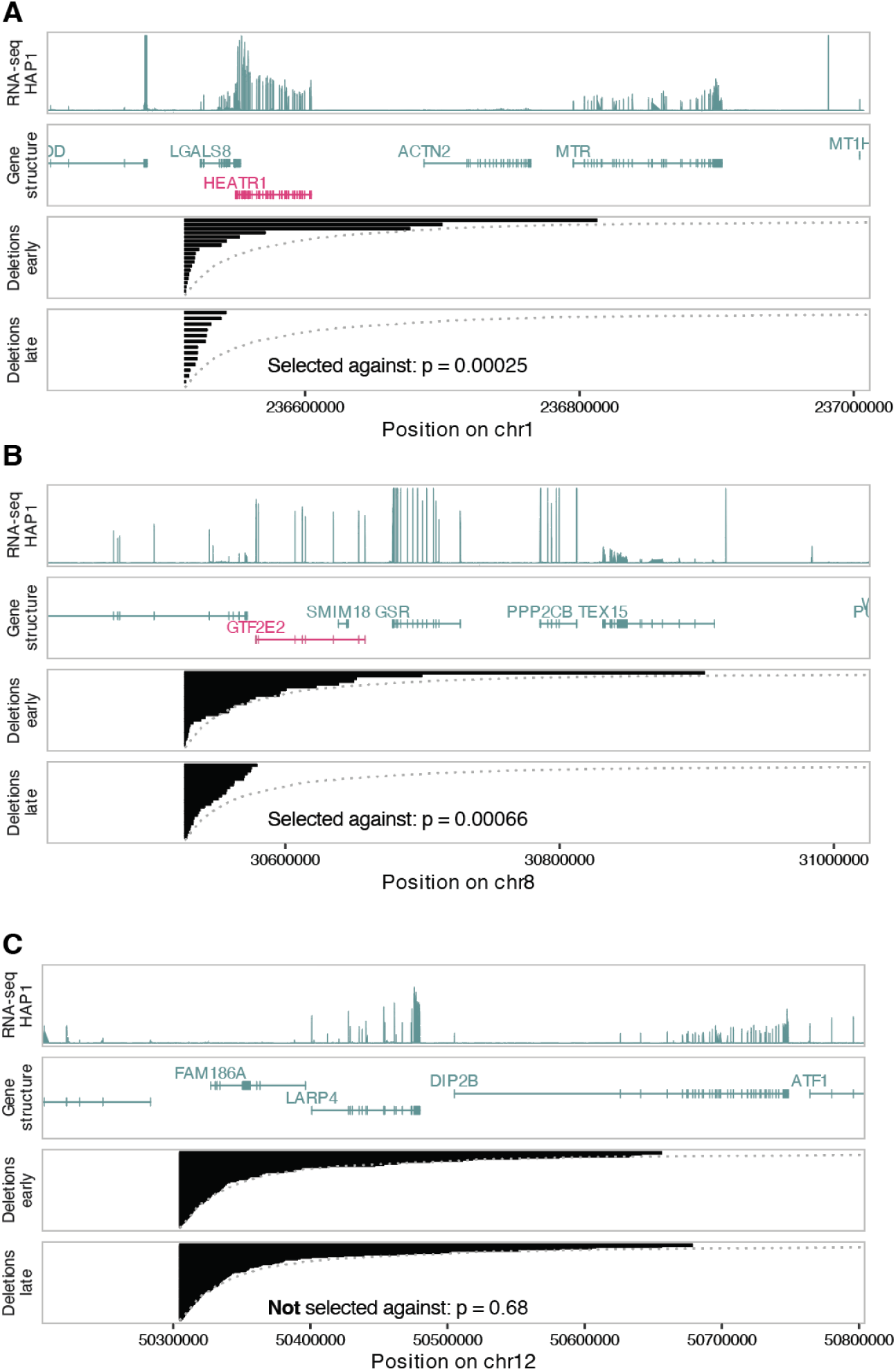
Examples of regions with or without selection. (**A**) Genomic regions surrounding an individual Shred-seq beacon integration site on chromosome 1 with evidence of negative selection. From top to bottom, tracks show HAP1 RNA-seq coverage, annotated gene structures (genes marked in red are common essential), deletion intervals observed at the early time point, and deletion intervals observed at the late time point. Dotted curves indicate the expected log-normal deletion profile for deletions originating from the corresponding beacon. The P-value is from a one-sided student’s t-test comparing log lengths of the observed distribution to the mean of the log-normal expectation. (**B**) As in panel **A** but for another genomic region with evidence of selection on chromosome 8. (**C**) As in panel **A** but for a genomic region without evidence of selection.

**Figure S12.**
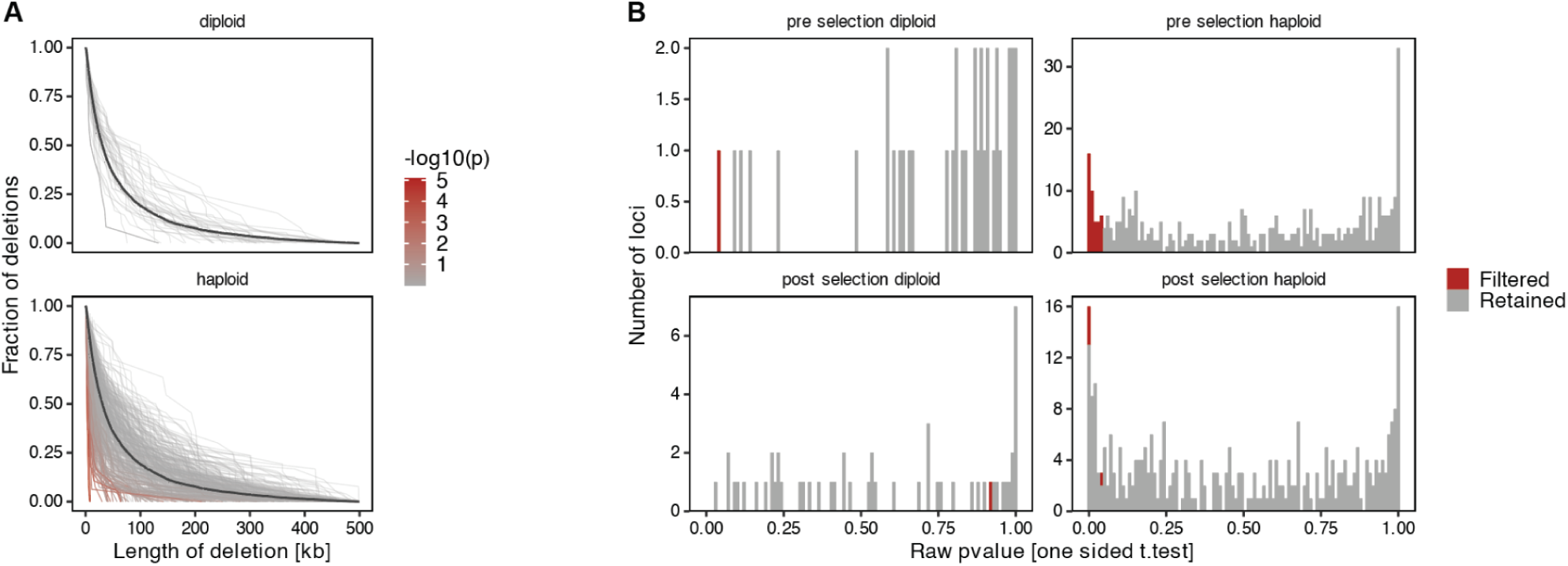
Deviations from log-normal expectation. (**A**) Cumulative deletion length distributions for beacon sites with >10 deletions, separated by ploidy. Each line represents the deletion length profile for an individual beacon. Line color indicates the strength of deviation from the fitted log-normal expectation (one-sided Student’s t-test). (**B**) Distribution of raw P-values for deviation from the log-normal expectation across beacon sites, separated by ploidy and by pre-selection or post-selection time point. Red bars indicate beacon sites filtered from downstream analyses because their pre-selection deletion profiles already deviated from the expected distribution.

**Figure S13.**
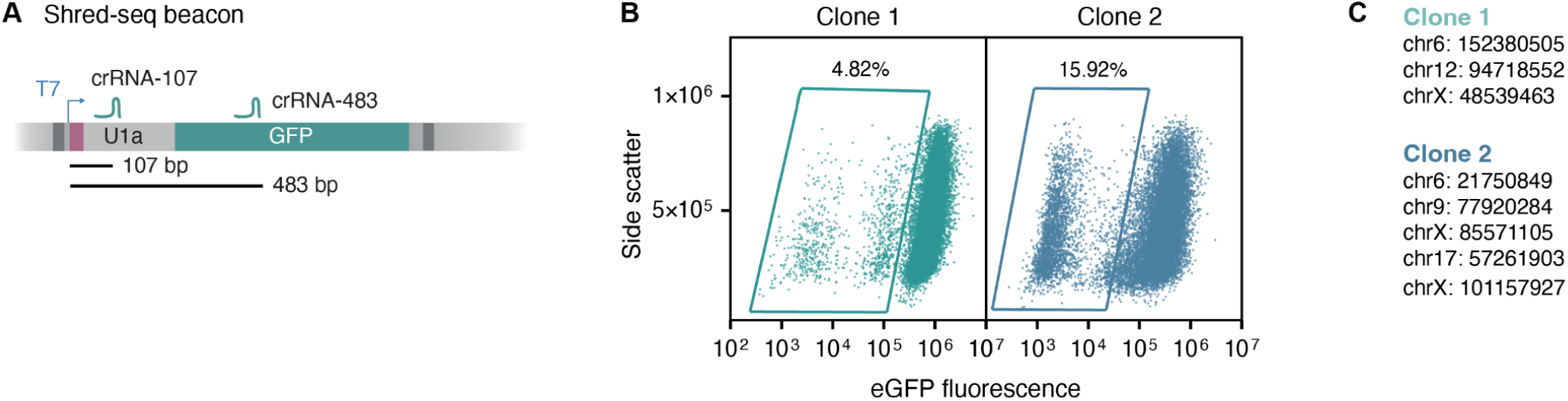
Single-clones with v3 beacons. (**A**) Schematic illustrating the binding position of two crRNAs on a genomically-integrated beacon. Lines and numbers represent distances from the first PAM-proximal base of the crRNA to the first base transcribed by T7 polymerase. (**B**) Side scatter (y-axis) compared to eGFP fluorescence (x-axis) for two selected clones (columns) nucleofected with plasmids encoding Cas3, Cascade and crRNA-107. Each dot represents a single-cell measured by flow cytometry. Illustrative gates that highlight the fraction of GFP-negative cells are drawn. (**C**) Beacon integration locations for clone 1 and clone 2.

**Figure S14.**
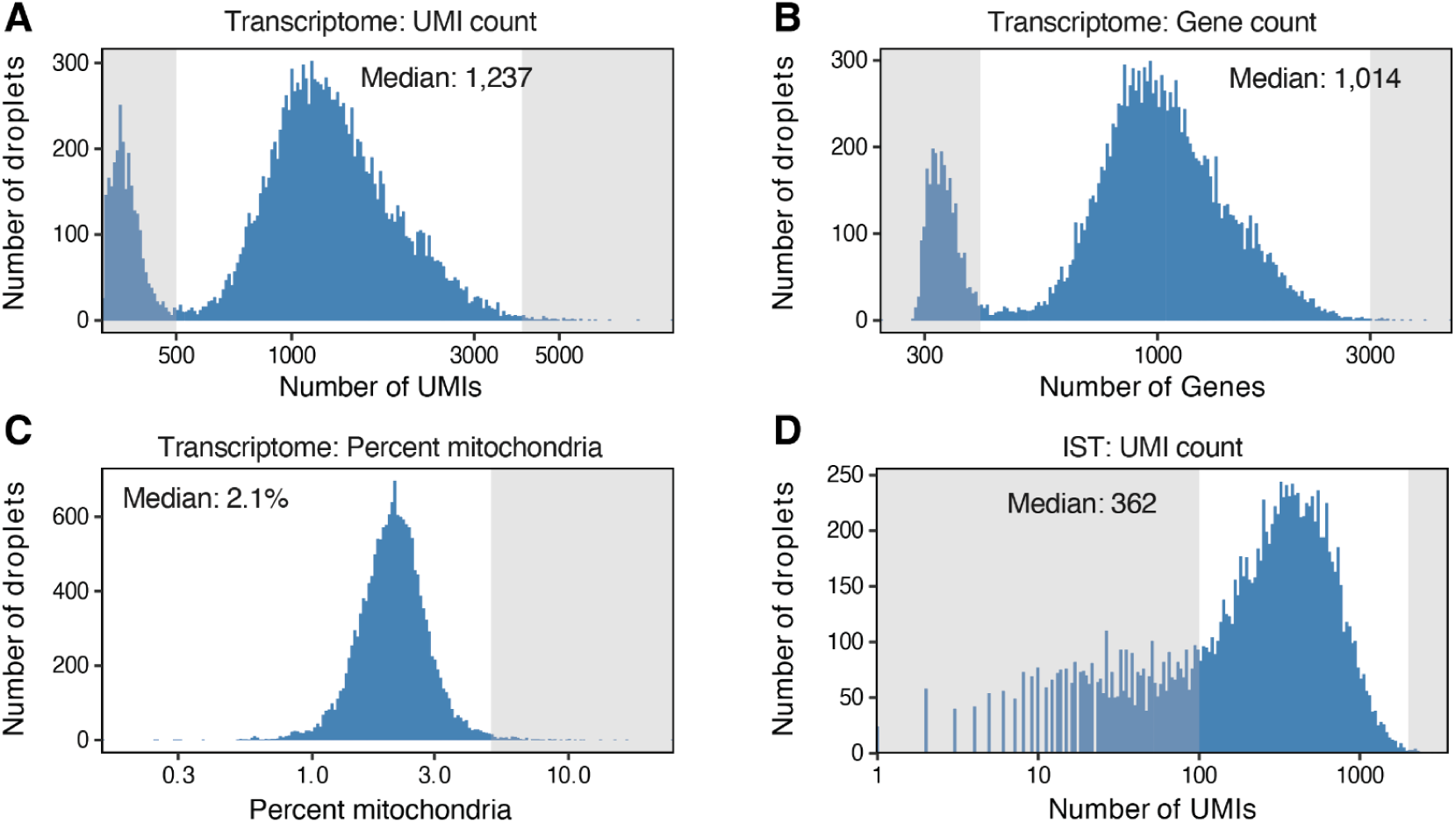
Quality control of single-cell transcriptome and IST libraries. (**A**) Distribution of transcriptome UMI counts (x-axis) per droplet after processing with Cell Ranger. The median transcriptome UMI count after filtering is indicated. Shaded regions mark excluded droplets. (**B**) As in panel **A** but for the number of identified genes. (**C**) As in panel **A** but for the mitochondrial transcript fraction per droplet. (**D**) Distribution of T7 IST UMI counts per droplet. The median IST UMI count is indicated, and shaded regions mark excluded droplets.

**Figure S15.**
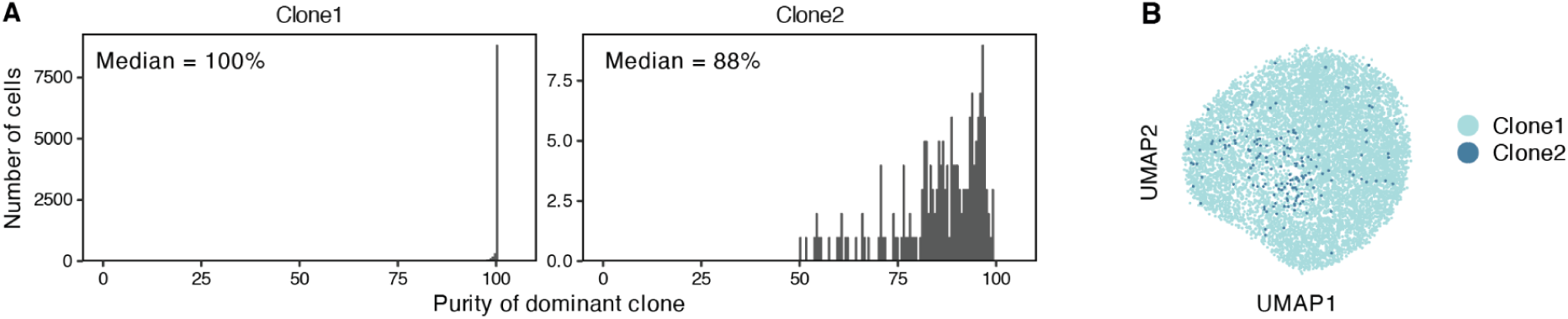
High clonal purities in IST libraries. (**A**) Distribution of dominant-clone purity among cells assigned to each scShred-seq clone based on beacon barcodes. Purity was calculated as the fraction of UMIs assigned to the dominant clone in each cell. Median dominant-clone purity is indicated for each clone. (**B**) UMAP projection of single-cell transcriptomes. Cells are colored by assigned clone identity.

**Figure S16.**
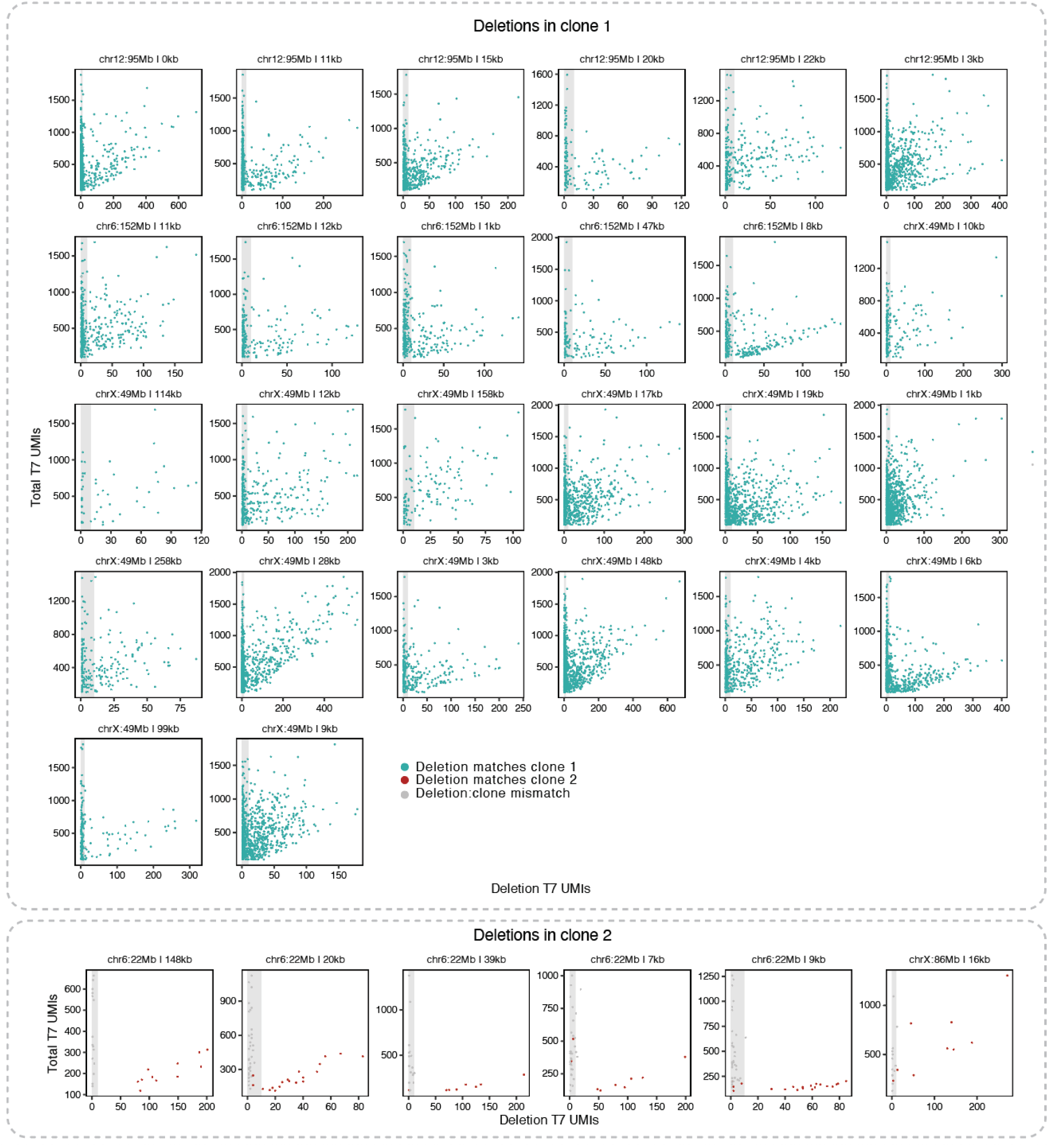
Single-cell deletion calling and junction recovery from IST libraries. Deletion-specific T7 UMIs (x-axis) and total T7 UMIs (y-axis) for deletions with more than 20 cells, separated by clone of origin for deletion (top clone 1, bottom clone 2). Cells (dots) are colored based on the matching of the deletion assignment and clone identity. Cells in the shaded area were excluded.

**Figure S17.**
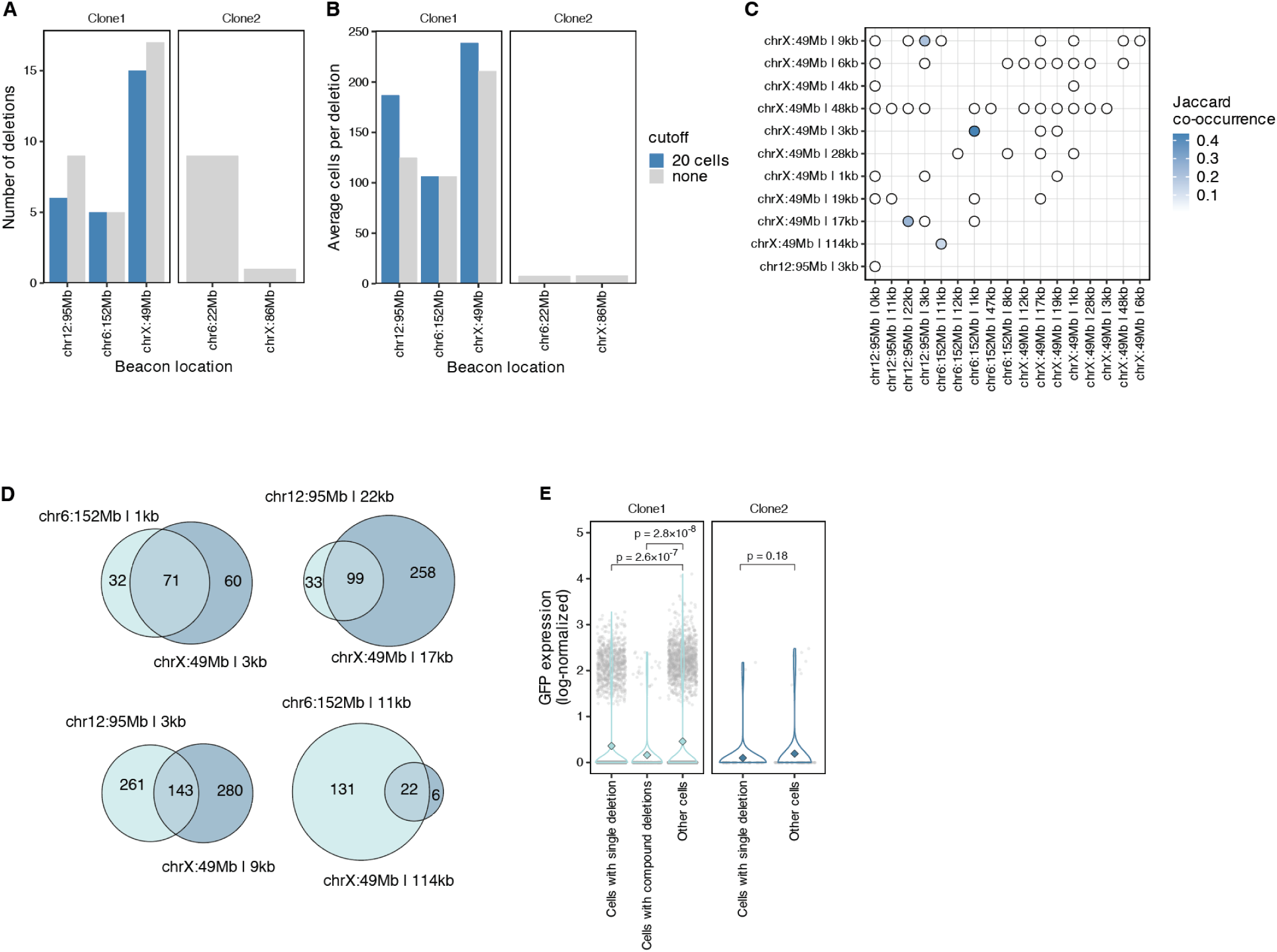
Validation of single-cell deletion calls. (**A**) Number of deletions (y-axis) across various beacons (x-axis) in the two profiled clones (panels), colored based on whether a 20 cell per deletion cutoff was used or not. (**B**) As in panel **A** but showing the average number of cells per deletion. (**C**) Co-occurrence of deletions within the same single-cells. Each point represents a pair of deletions detected in at least three cells, and color indicates the Jaccard co-occurrence index for that deletion pair. (**D**) Size-proportional Venn diagrams for the four frequently co-occurring deletions. The number of cells with one or both confident deletion assignments are indicated. (**E**) *GFP* expression (y-axis) for cells based on deletion status (x-axis) across various clones (panels and colors). Markers represent individual cells, violins show the distribution, and the diamond represents the mean expression. P-values were calculated using a two-tailed Wilcoxon test.

**Figure S18.**
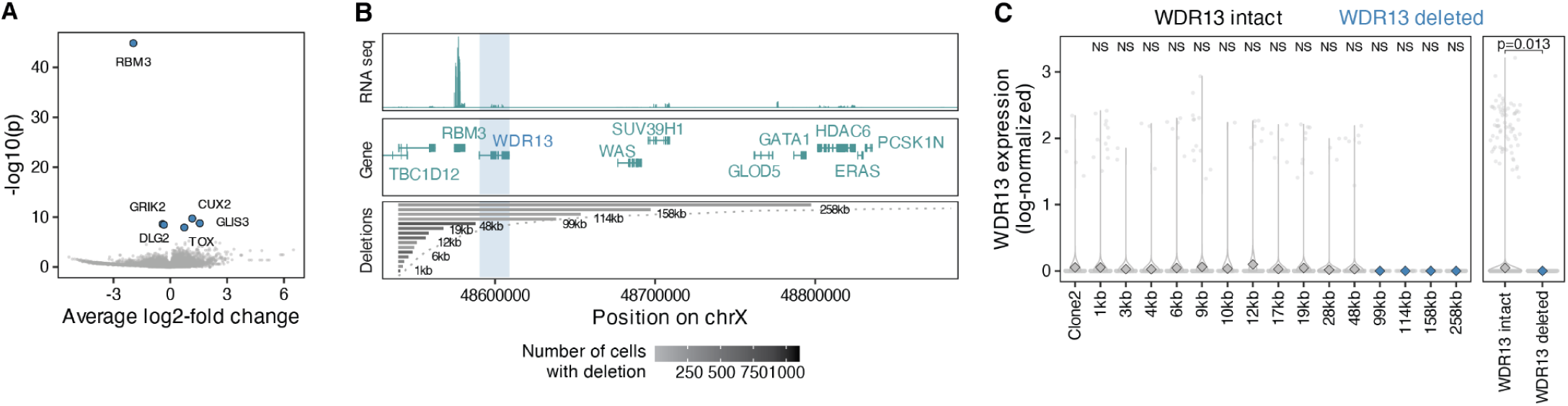
scShred-seq associates deletion genotypes to transcriptional phenotypes. (**A**) P-values (y-axis) and log2-fold changes (x-axis) from genome-wide differential expression testing (two-tailed Wilcoxon test) between cells with deletions from Figure 5H that overlap or spare *RBM3*. Points represent individual genes. The six most significantly differentially expressed genes are highlighted in blue. (**B**) Deletions and genomic features (panels) at a region on chromosome X bearing an integrated beacon (x-axis). From top to bottom: (i) RNA-seq coverage in HAP1 cells. (ii) Exon structure and names of genes in region. (iii) Deletions observed in the scShred-seq experiment, with each line corresponding to the length of one deletion and shaded according to the number of cells with deletion evidence. Genomic locations overlapping the *WDR13* gene are shaded. (**C**) *WDR13* expression (y-axis) for cells based on deletion status (x-axis). Deletions overlapping *WDR13* are highlighted in blue. Markers represent individual cells, violins show the distribution, and the diamond represents the mean expression. P-values were calculated using a two-tailed Wilcoxon test and adjusted for multiple hypothesis testing (Benjamini-Hochberg) comparing cells with deletions to cells from clone 2. The right panel shows a comparison between cells with deletions that overlap or spare *WDR13*.

## Supplementary Note 1. A constrained deletion profile with no overlapping annotated essential gene

One beacon integration site (chr3:49419658) exhibited a significantly constrained (FDR = 0.02) post-selection deletion-length profile in haploid cells, despite the absence of an annotated essential protein-coding gene in the downstream region spanned by deletions originating at this beacon. The nested deletion series directly overlapped *TCTA, AMT, NICN1, DAG1*, and *BSN* (**Fig. SN1**; on the next page). Each of these genes is transcribed in HAP1 cells, but none are classified as essential either in the HAP1-specific CRISPR screen or in the Dependency Map dataset. Nevertheless, there are several essential genes in proximity.

Several explanations, not all of which are mutually exclusive, could account for the observed constraint: (i) One or more genes directly overlapped by the deletions is essential in HAP1 cells but was not detected as such by the CRISPR screens. (ii) The transcription start site of *RHOA* lies 5.7 kb upstream of the beacon and is oriented away from the beacon (**Fig. SN1**). RHOA fitness in HAP1 is 0.34; we consider anything below 0.5 essential. Deletions from this beacon could remove a *RHOA* regulatory element or impact *RHOA* expression through the removal of nearby expressed genes through supercoiling-mediated feedback. (iii) There is a cluster of highly essential genes further downstream that includes *MST1* (fitness 0.42) and *GMPPB* (fitness 0.12) (**Fig. SN1**). Deletions that do not directly overlap these genes could nevertheless alter their expression by removing distal regulatory elements or disrupting local chromatin architecture. Notably, there is an accessible CTCF site (black arrow in **Fig. SN1**) that coincides with the approximate boundary beyond which post-selection deletions become depleted. Disruption of that site might affect the expression of the essential genes nearby. (iv) Finally, because we test many beacon-level deletion profiles and apply an FDR of 10%, the observed constraint in this region may represent a statistical false positive.

Although this locus lacks an immediately apparent essential gene directly explaining its constrained deletion profile, it illustrates how Shred-seq may identify candidate fitness-relevant intervals that are not readily interpretable from existing gene essentiality annotations alone. Targeted follow-up experiments would be required to determine whether the observation corresponds to a missed gene-level dependency, disruption of local regulatory architecture, or a statistical false positive.

**Figure SN1.**
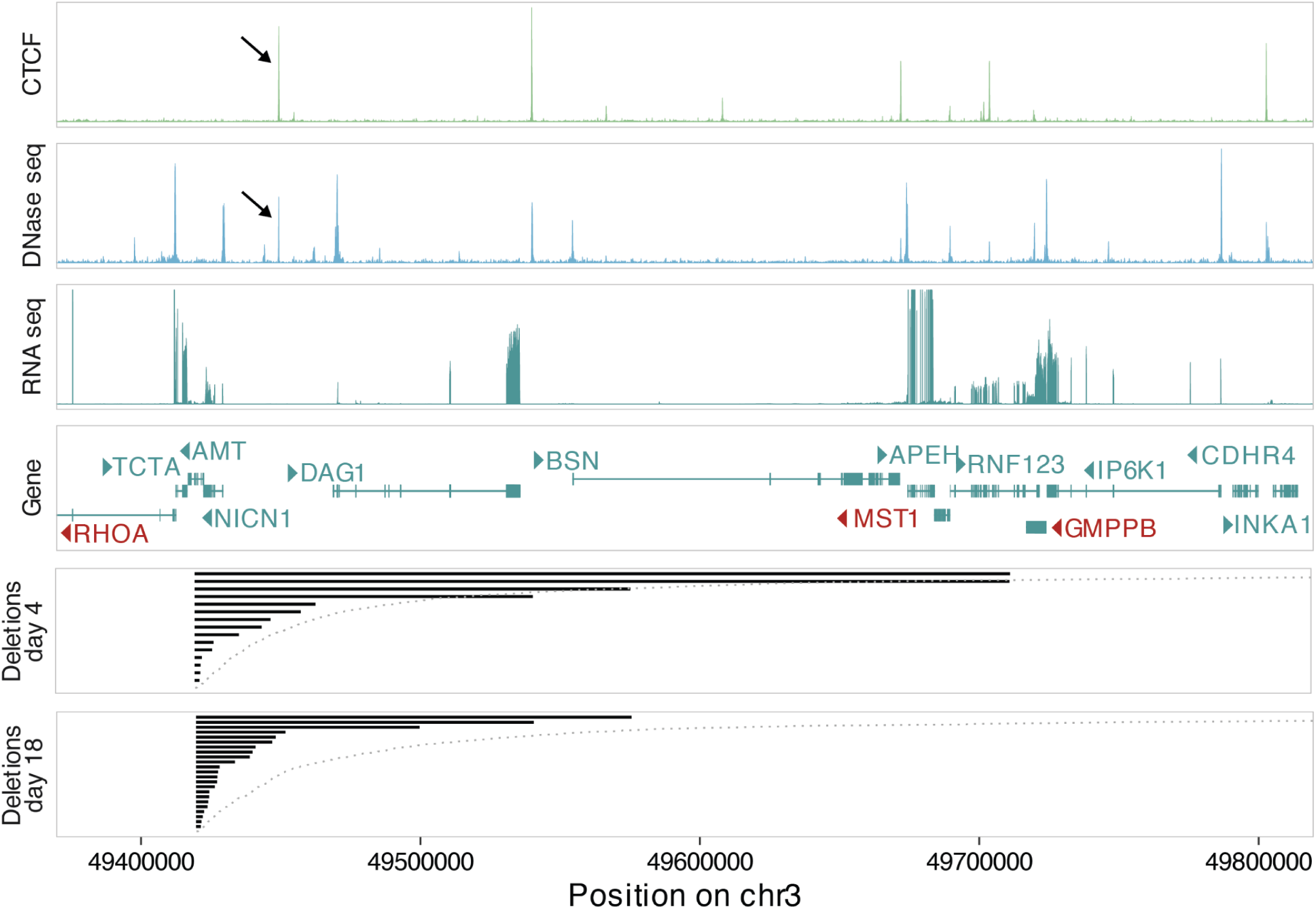
Example of a beacon integration site with a constrained deletion profile, but no established essential gene downstream. Genomic features and deletions (panels) at a region on chromosome 3 bearing an integrated beacon. From top to bottom: (i) CTCF-Chip-Seq coverage track in HAP1 cells. (ii) DNase-seq coverage track in HAP1 cells. (iii) RNA-seq coverage in HAP1 cells. (iv) Exon structure and names of genes in region. Essential genes are marked in Red. Arrows indicate direction of transcription. (v) Deletions observed on day 4, with each line corresponding to length of one deletion. (vi) Deletions observed on day 18. Arrows indicate the location of an accessible CTCF site.

## Notes

https://github.com/pinglaylab/deletion_scanning

