## Supplementary Figures 1-18 for "Gigabase-scale deletion scanning of the human genome"

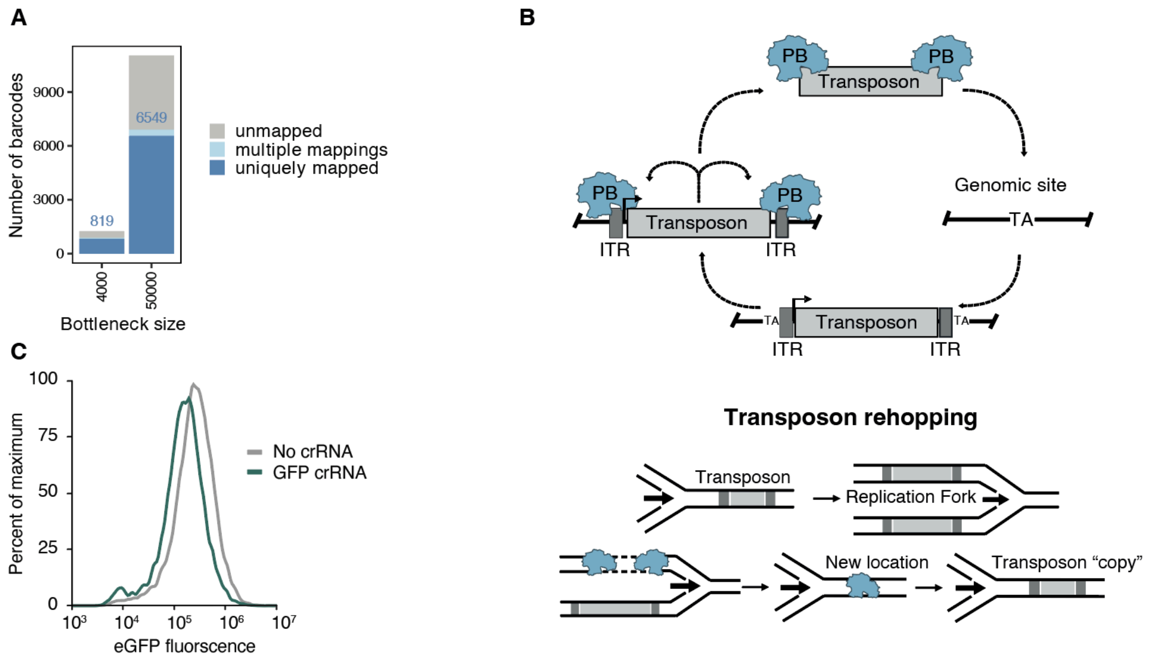


**Figure S1. Shred-seq pilot** (**A**) Number of barcodes recovered (y-axis) after bottlenecking Shred-seq pilot populations at two different bottleneck sizes (x-axis). Stacked bars indicate barcodes that could be mapped uniquely, barcodes mapping to multiple loci, and unmapped barcodes. (**B**) Schematic illustrating PiggyBac transposon re-hopping, which can generate multiple genomic locations associated with the same beacon barcode, potentially during DNA replication. (**C**) eGFP fluorescence distributions for cells transfected with a polycistronic Cas3-Cascade construct and either a GFP-targeting crRNA or no crRNA control seven days after Cas3 nucleofection.

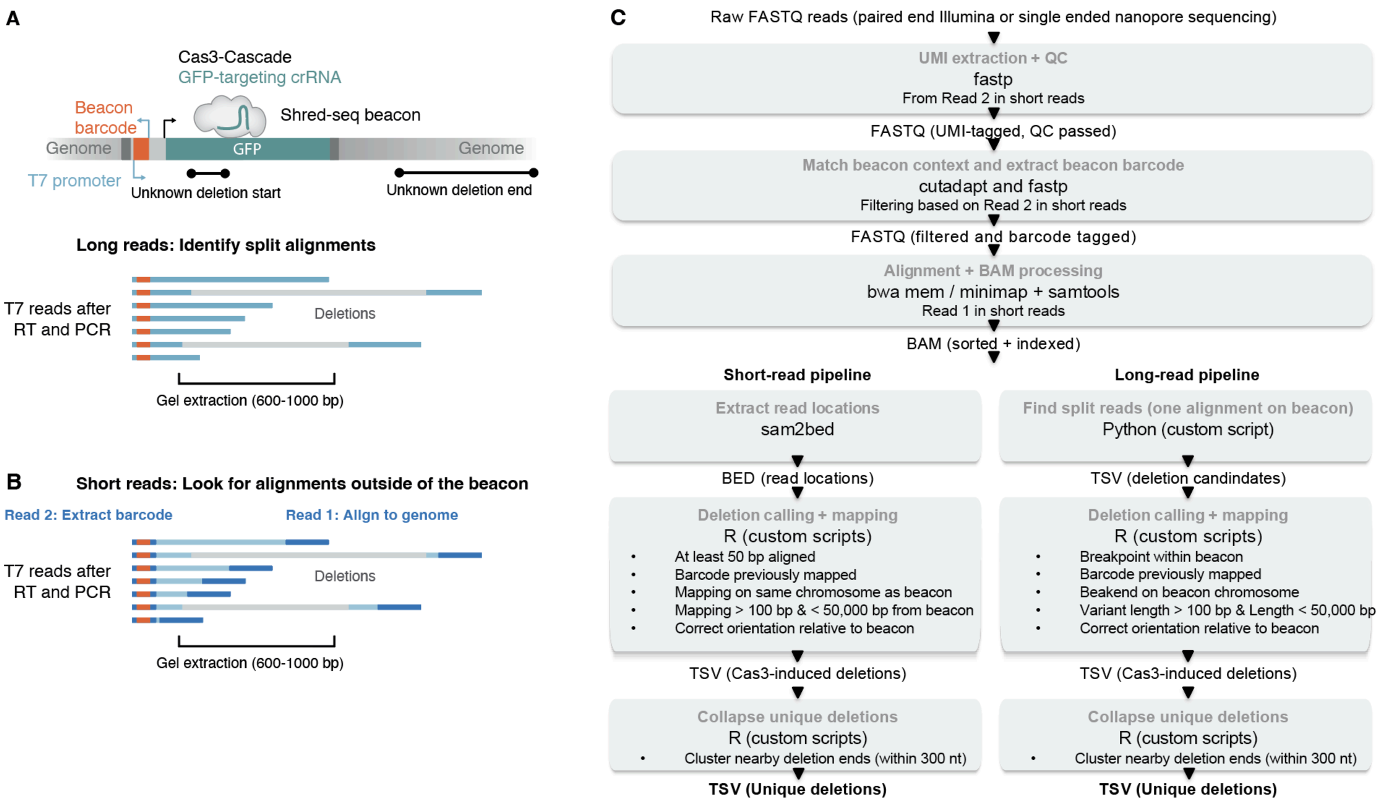


**Figure S2. T7 deletion mapping pipeline.** (**A**) Schematic of long-read deletion mapping. Following T7 *in vitro* transcription, reverse transcription, and PCR, long-read sequencing captures individual molecules spanning the beacon barcode and deletion junction, allowing direct identification of split alignments and deletion breakpoints. (**B**) Schematic of short-read deletion mapping. Read 2 captures the beacon barcode, whereas Read 1 captures adjacent genomic sequence; deletion-supporting reads are identified as barcode-associated reads aligning outside the beacon sequence. (**C**) Overview of the computational pipeline used for deletion calls from raw sequencing data. FASTQ files were processed by quality filtering, barcode extraction, alignment, deletion calling, mapping to previously established beacon integration sites, and collapsing of nearby events into unique deletions for both short-read and long-read workflows.

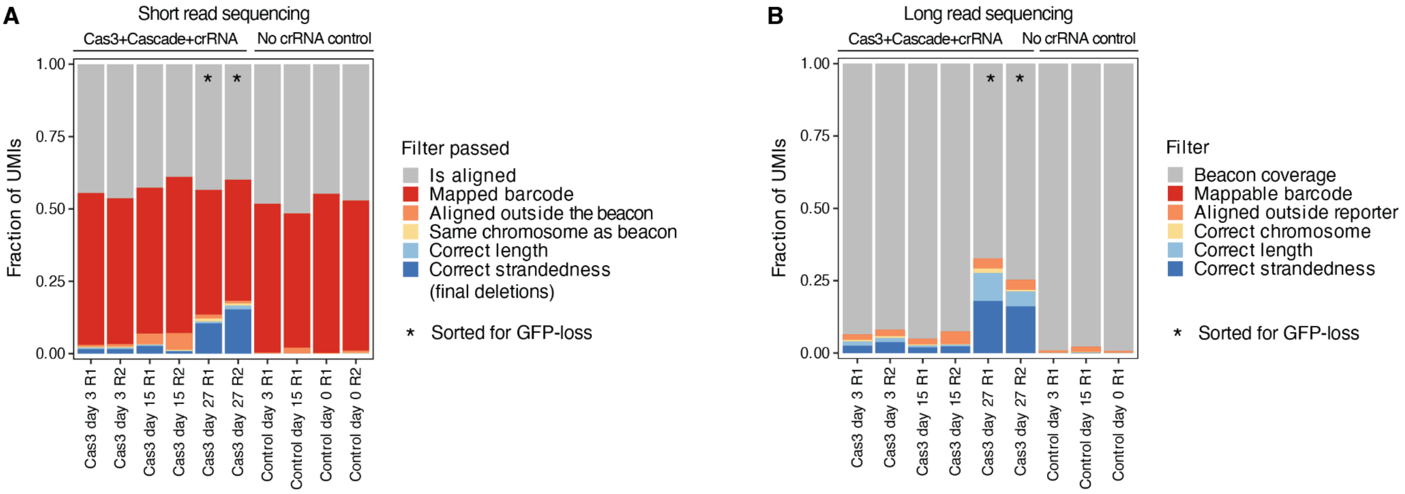


**Figure S3. High sensitivity deletion calling.** (**A**) Fraction of UMIs retained after successive filtering steps (y-axis) in the short-read pipeline across Cas3-treated and no-crRNA control samples (x-axis). Colored segments indicate the fraction of UMIs passing each criterion. Asterisks denote samples enriched for GFP loss. (**B**) As in panel **A**, but for long-read sequencing data. R1, R2 indicate replicate 1 and replicate 2.

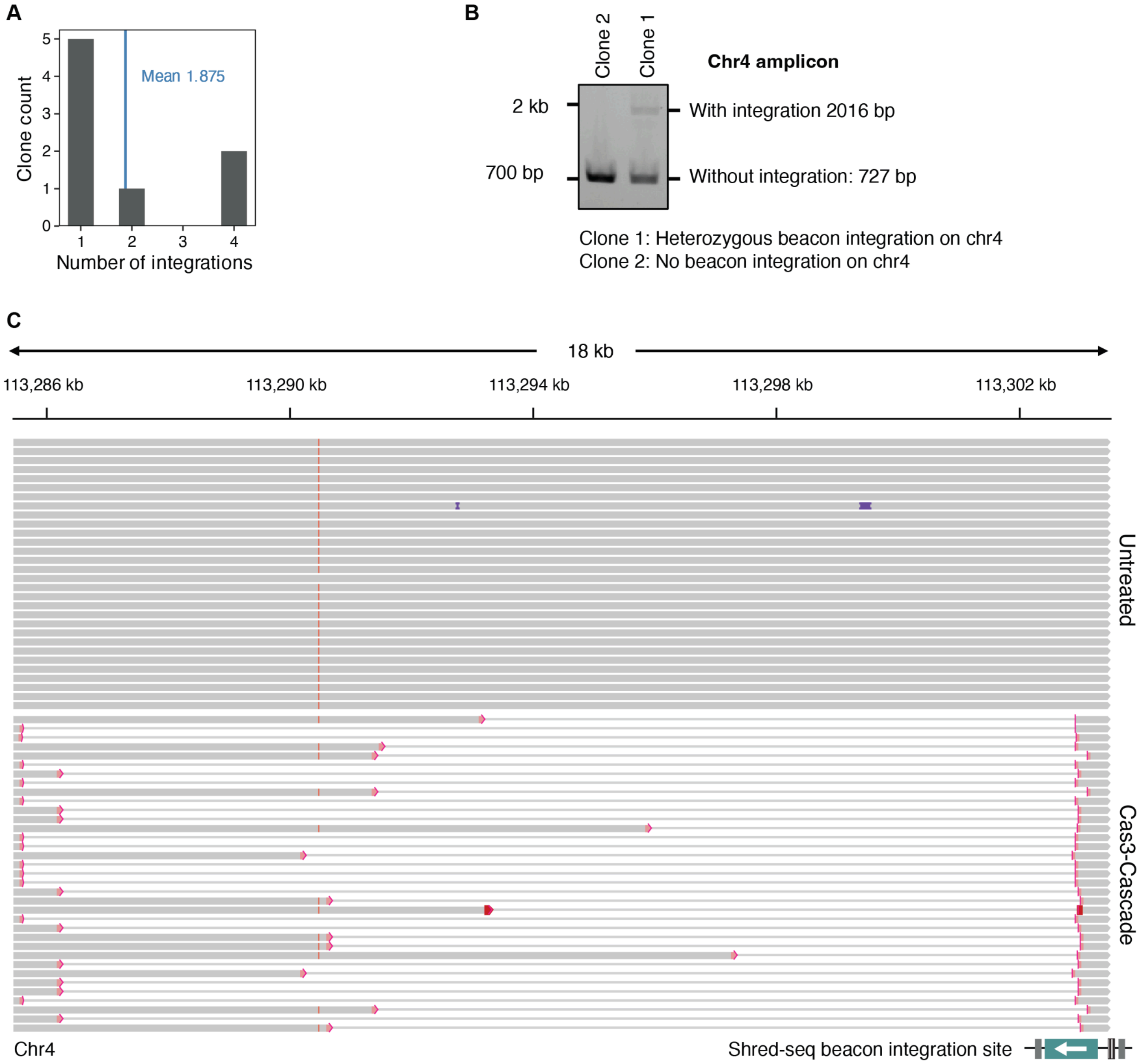


**Figure S4. Shred-seq in individual clones.** (**A**) Distribution of the number of beacon integrations detected per isolated clone from the pilot PiggyBac population. The vertical line indicates the mean number of integrations per clone. (**B**) PCR genotyping of individual clones with a candidate beacon integration on chromosome 4. Band sizes for alleles with and without integration are indicated. (**C**) Comparison of long-range amplicon sequencing profiles for cells from a clone with a beacon integration site on chromosome 4 that were left untreated (top) or transfected with Cas3, Cascade, and a GFP-targeting crRNA (bottom). Gray bars represent aligned sequencing reads and thin lines represent deletions. The Shred-seq beacon integration location is indicated at the bottom.

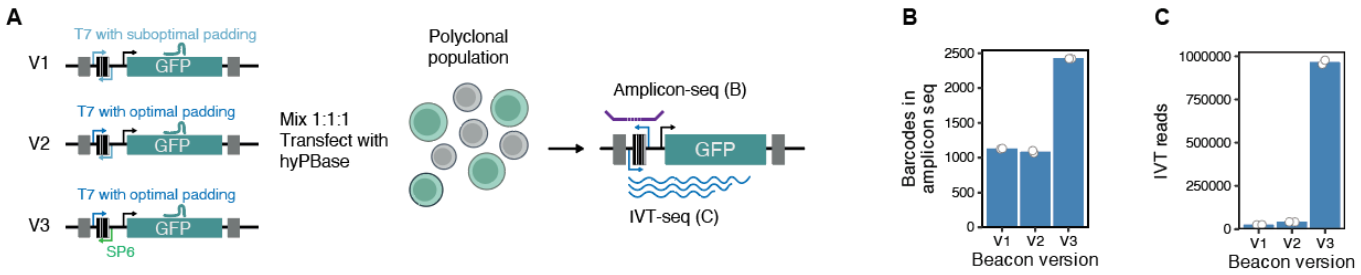


**Figure S5. Improving T7 readout.** (**A**) Schematic of Shred-seq beacon versions v1, v2, and v3 used to optimize T7-based readout. v2 includes improved T7 promoter padding, whereas v3 additionally replaces one T7 promoter with SP6 to reduce transcriptional interference. Mixed beacon populations were generated and analyzed by amplicon sequencing and IVT-seq. (**B**) Number of barcodes detected by amplicon sequencing (y-axis) for each beacon version (x-axis). (**C**) T7-derived IVT reads recovered from each beacon version.

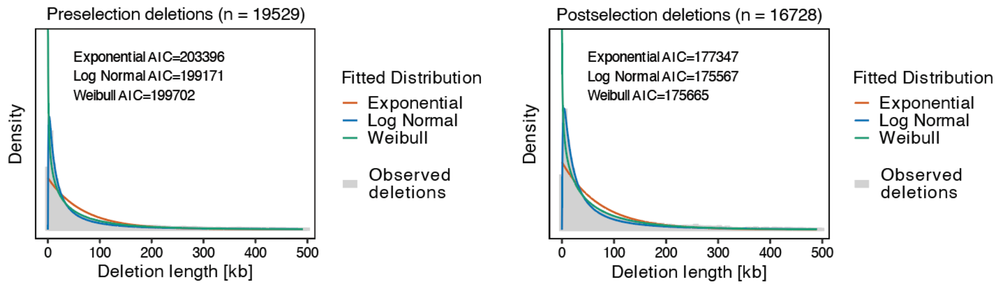


**Figure S6. Cas3 deletions follow a log-normal distribution.** Observed deletion length distributions (gray histogram) for pre-selection deletions (left) and post-selection deletions (right), together with fitted exponential, log-normal, and Weibull distributions (colored lines). Gray histograms indicate observed deletion lengths from a union of all pre-selection experiments. Insets report the Akaike information criterion (AIC) for each fit.

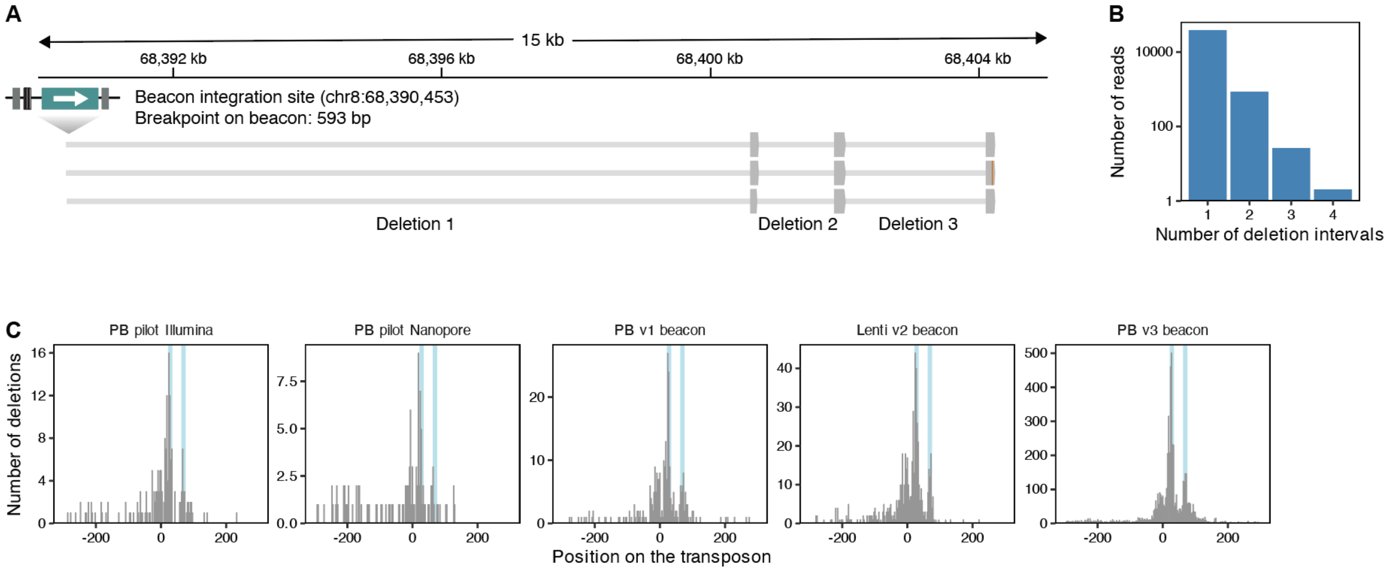


**Figure S7. Patterns of Cas3 deletion initiation.** (**A**) Example of sequencing reads with evidence for multiple successive deletions originating from a single beacon integration site. Gray bars represent aligned sequencing reads and thin lines represent deletions. (**B**) Number of sequencing reads (y-axis, log-scale) across a different number of deletion intervals (x-axis). (**C**) Histograms showing the position of deletion initiation relative to the beacon/protospacer across different experiments, beacon versions, delivery modalities, and sequencing strategies (panels). Vertical lines mark two regions with high deletion initiation.

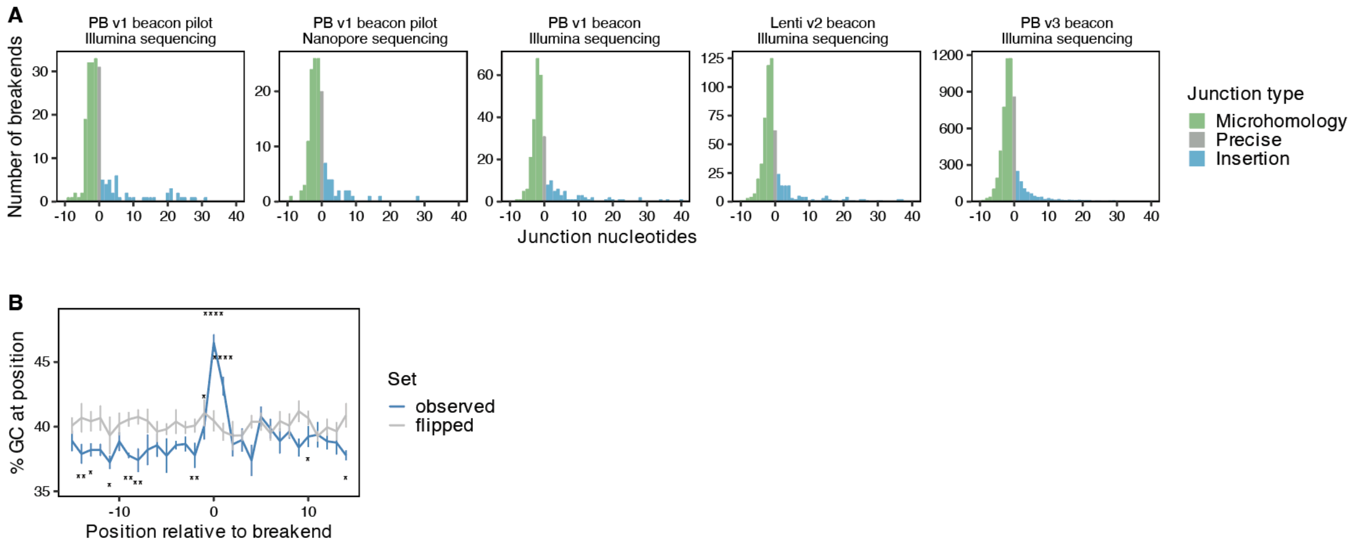


**Figure S8. Cas3 junctions are repaired by microhomology-mediated end joining.** (**A**) Distribution of junction nucleotide lengths (x-axis) across different experiments and sequencing strategies (panes). Negative values indicate overlapping sequence at the junction (microhomology), zero indicates a precise junction, and positive values indicate inserted nucleotides. Bars are colored by inferred repair class. (**B**) Average GC content (y-axis) across a 30 bp window around the deletion breakend (x-axis) for observed deletions and flipped controls (lines and colors). Whiskers show the standard error of the mean. P-values were computed with a two-sided test for equality of proportions and corrected for multiple testing across positions (Benjamini-Hochberg).

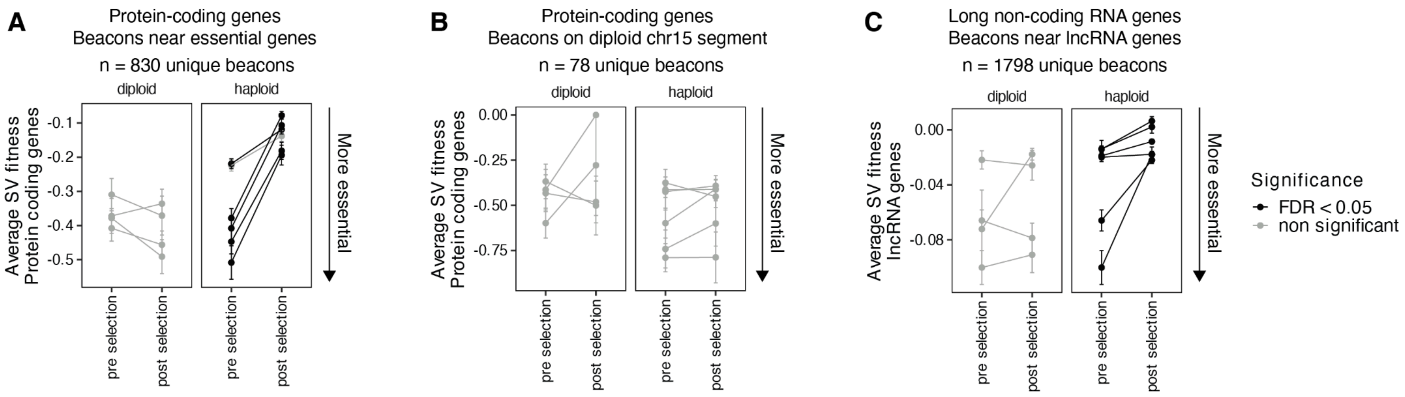


**Figure S9. Essential protein-coding and long non-coding RNA genes are depleted from post-selection deletions.** (**A**) The average protein-coding fitness of deletions (y-axis) originating from a subset of beacons within 500 kb of a core-essential gene (mean Chronos score of 1000+ dependency map cell lines < 0.5) across Shred-seq experiments and biological replicates (points and lines) pre and post selection (x-axis) separated by ploidy (panels) and shaded by FDR adjusted for multiple hypothesis testing. Whiskers represent the standard error of mean. (**B**) As in panel **A** but for a subset of beacons on a 30 Mb interval on chromosome 15 that is diploid in HAP1 cells (chr15:60812801-89346769). (**C**) As in panel **A** but for long-non coding RNA fitness and a subset of beacons with expressed lncRNAs (TPM > 1) downstream.

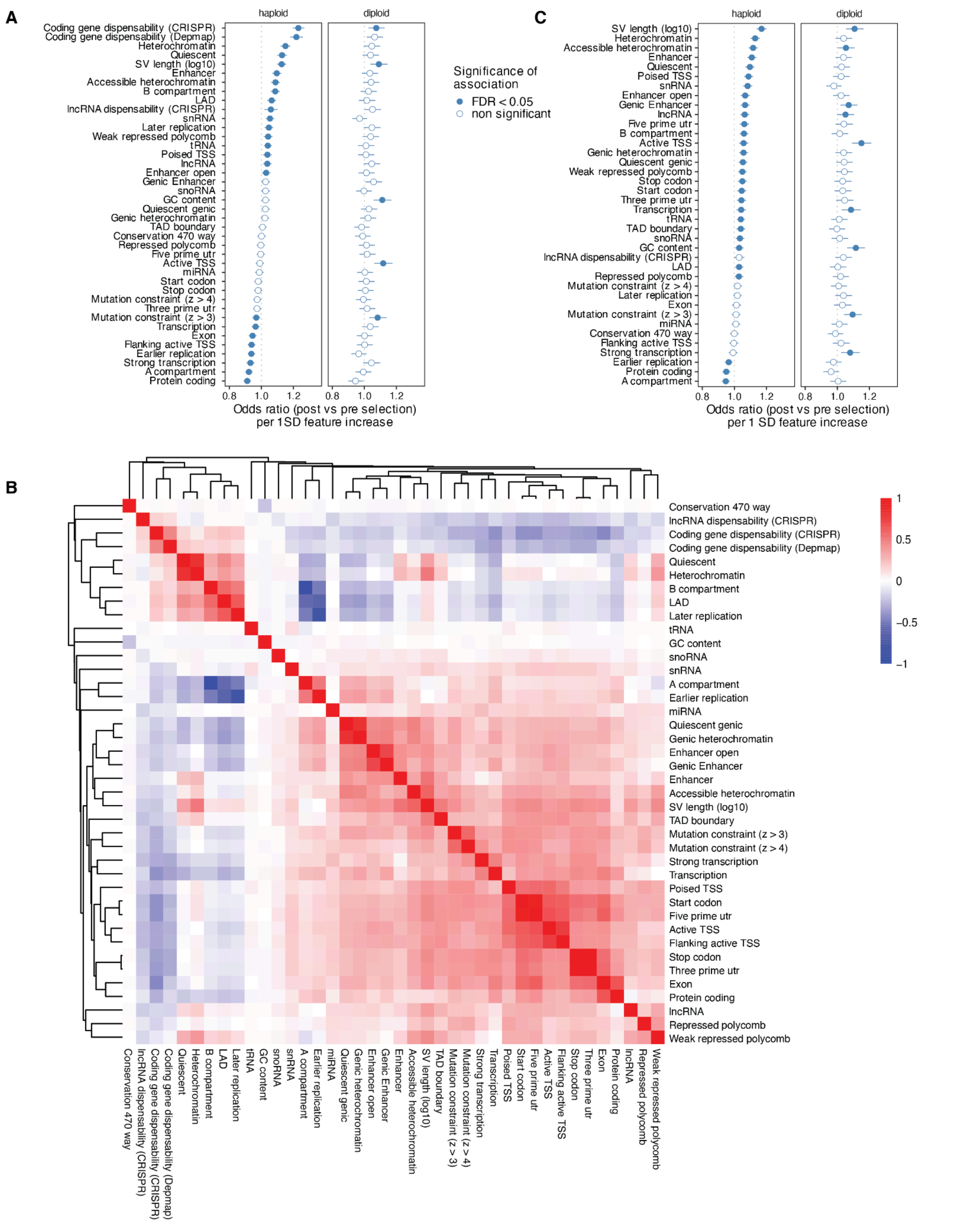


**Figure S10. A combination of features shapes SV selection.** (**A**) Odds ratios per standard deviation increase (x-axis) in features (y-axis) between post vs pre-selection variants in haploid or diploid HAP1 cells (panels). Whiskers indicate 95% confidence intervals. (**B**) Heatmap of pairwise Spearman rank correlations between genomic features in pre-selection haploid deletions. Correlations were calculated using pairwise complete observations, and colors denote the Spearman correlation coefficient (ρ). (**C**) As panel **A** but including protein-coding gene essentiality as a covariate.

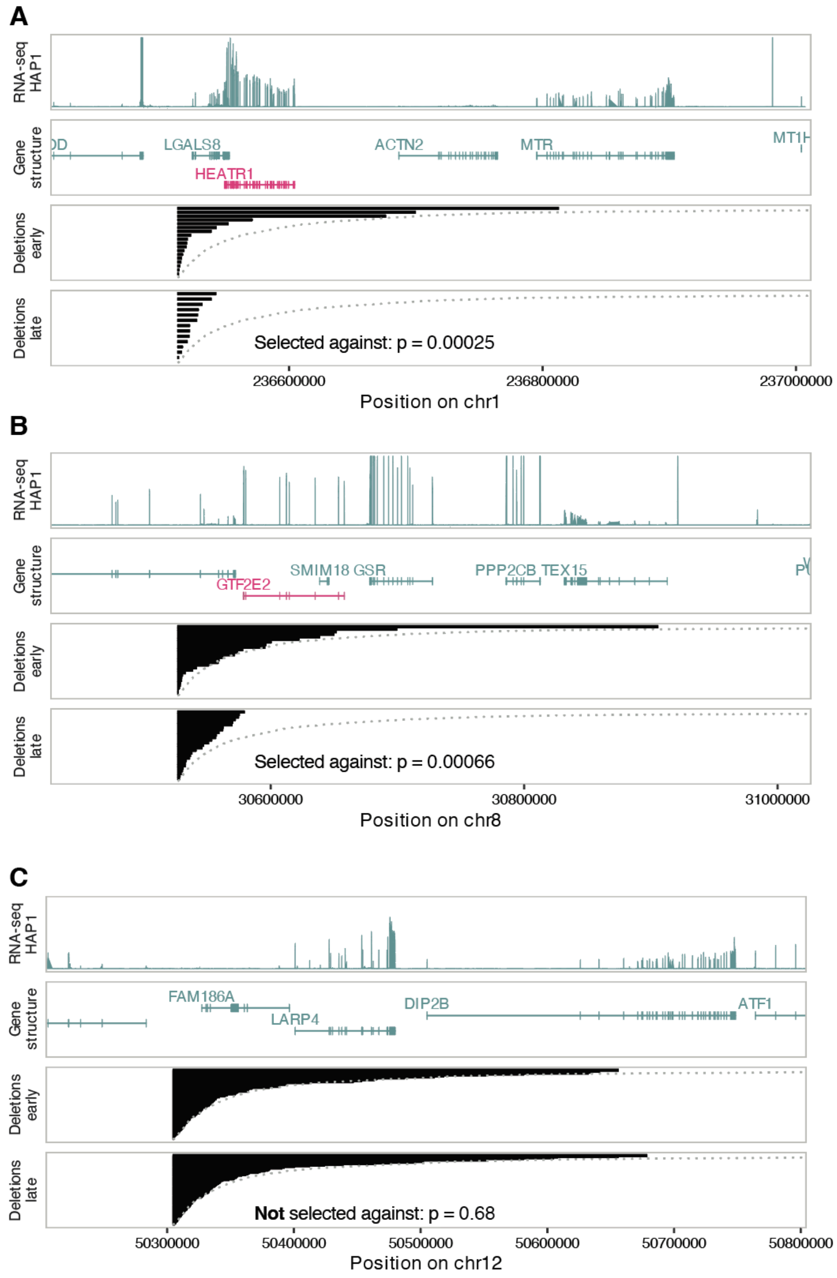


**Figure S11. Examples of regions with or without selection.** (**A**) Genomic regions surrounding an individual Shred-seq beacon integration site on chromosome 1 with evidence of negative selection. From top to bottom, tracks show HAP1 RNA-seq coverage, annotated gene structures (genes marked in red are common essential), deletion intervals observed at the early time point, and deletion intervals observed at the late time point. Dotted curves indicate the expected log-normal deletion profile for deletions originating from the corresponding beacon. The P-value is from a one-sided student's t-test comparing log lengths of the observed distribution to the mean of the log-normal expectation. (**B**) As in panel **A** but for another genomic region with evidence of selection on chromosome 8. (**C**) As in panel **A** but for a genomic region without evidence of selection.

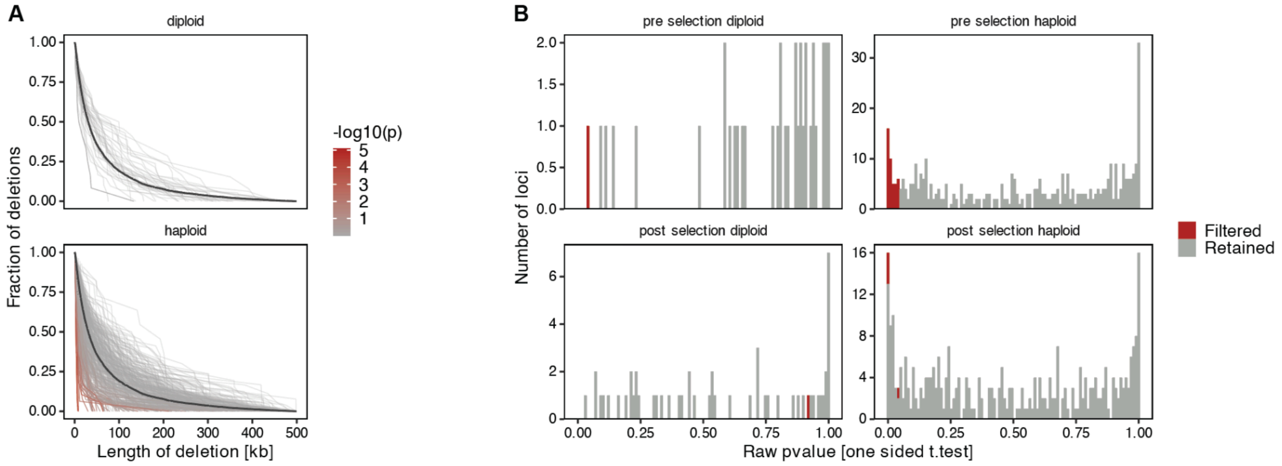


**Figure S12. Deviations from log-normal expectation.** (**A**) Cumulative deletion length distributions for beacon sites with >10 deletions, separated by ploidy. Each line represents the deletion length profile for an individual beacon. Line color indicates the strength of deviation from the fitted log-normal expectation (one-sided Student's t-test). (**B**) Distribution of raw P-values for deviation from the log-normal expectation across beacon sites, separated by ploidy and by pre-selection or post-selection time point. Red bars indicate beacon sites filtered from downstream analyses because their pre-selection deletion profiles already deviated from the expected distribution.

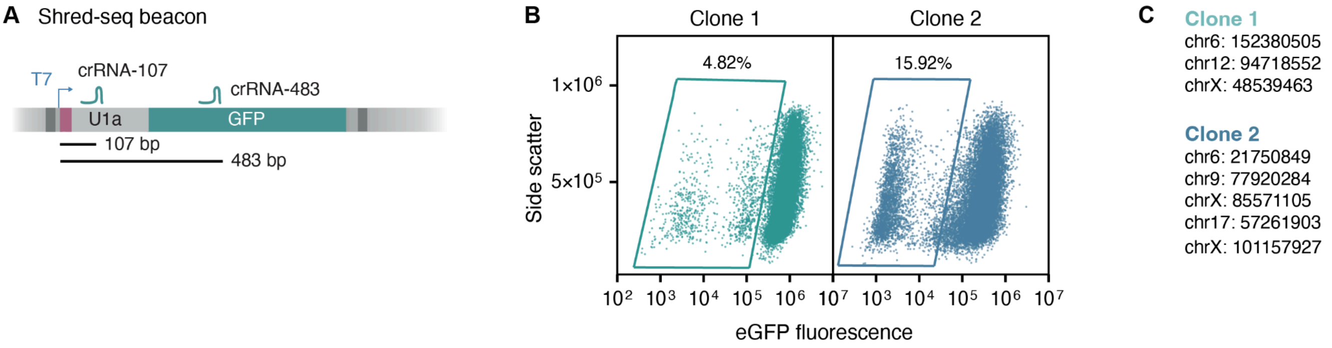


**Figure S13. Single-clones with v3 beacons.** (**A**) Schematic illustrating the binding position of two crRNAs on a genomically-integrated beacon. Lines and numbers represent distances from the first PAM-proximal base of the crRNA to the first base transcribed by T7 polymerase. (**B**) Side scatter (y-axis) compared to eGFP fluorescence (x-axis) for two selected clones (columns) nucleofected with plasmids encoding Cas3, Cascade and crRNA-107. Each dot represents a single-cell measured by flow cytometry. Illustrative gates that highlight the fraction of GFP-negative cells are drawn. (**C**) Beacon integration locations for clone 1 and clone 2.

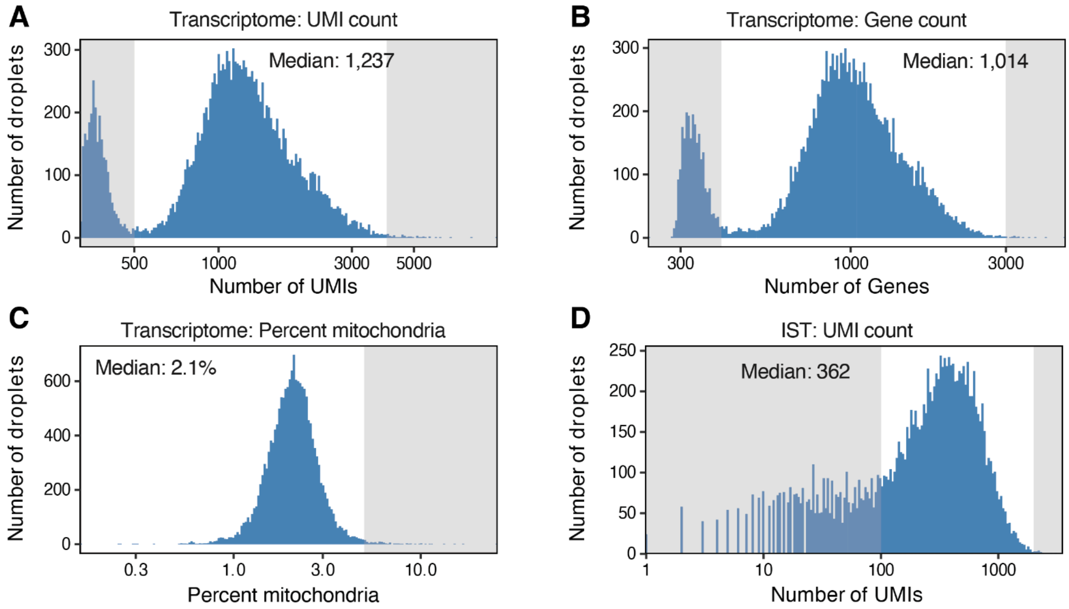


**Figure S14. Quality control of single-cell transcriptome and IST libraries.** (**A**) Distribution of transcriptome UMI counts (x-axis) per droplet after processing with Cell Ranger. The median transcriptome UMI count after filtering is indicated. Shaded regions mark excluded droplets. (**B**) As in panel **A** but for the number of identified genes. (**C**) As in panel **A** but for the mitochondrial transcript fraction per droplet. (**D**) Distribution of T7 IST UMI counts per droplet. The median IST UMI count is indicated, and shaded regions mark excluded droplets.

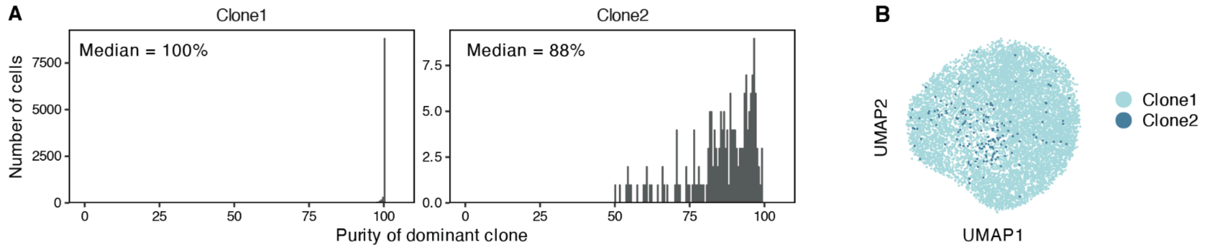


**Figure S15. High clonal purities in IST libraries.** (**A**) Distribution of dominant-clone purity among cells assigned to each scShred-seq clone based on beacon barcodes. Purity was calculated as the fraction of UMIs assigned to the dominant clone in each cell. Median dominant-clone purity is indicated for each clone. (**B**) UMAP projection of single-cell transcriptomes. Cells are colored by assigned clone identity.

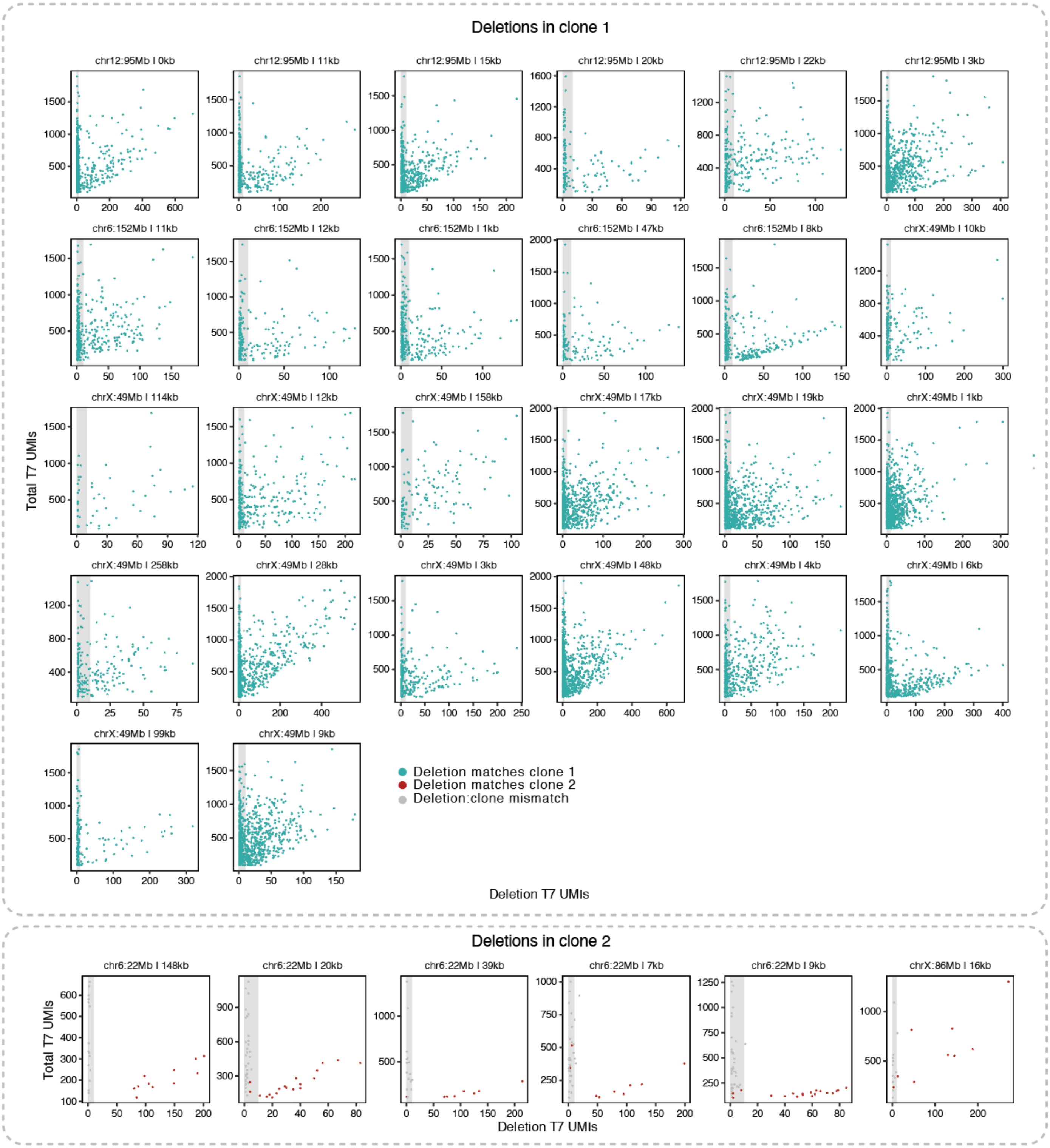


**Figure S16. Single-cell deletion calling and junction recovery from IST libraries.** Deletion-specific T7 UMIs (x-axis) and total T7 UMIs (y-axis) for deletions with more than 20 cells, separated by clone of origin for deletion (top clone 1, bottom clone 2). Cells (dots) are colored based on the matching of the deletion assignment and clone identity. Cells in the shaded area were excluded.

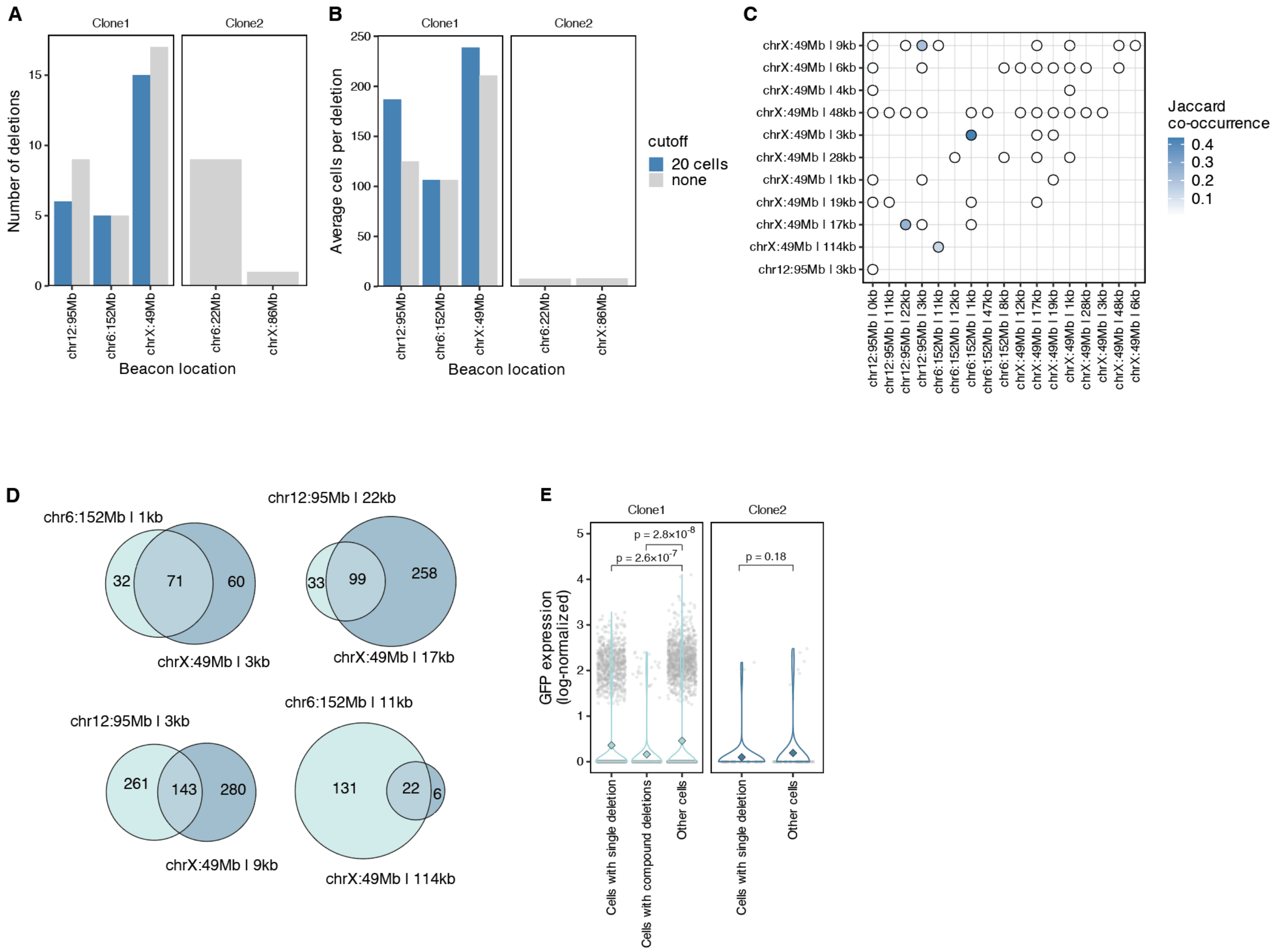


**Figure S17. Validation of single-cell deletion calls.** (**A**) Number of deletions (y-axis) across various beacons (x-axis) in the two profiled clones (panels), colored based on whether a 20 cell per deletion cutoff was used or not. (**B**) As in panel **A** but showing the average number of cells per deletion. (**C**) Co-occurrence of deletions within the same single-cells. Each point represents a pair of deletions detected in at least three cells, and color indicates the Jaccard co-occurrence index for that deletion pair. (**D**) Size-proportional Venn diagrams for the four frequently co-occurring deletions. The number of cells with one or both confident deletion assignments are indicated. (**E**) *GFP* expression (y-axis) for cells based on deletion status (x-axis) across various clones (panels and colors). Markers represent individual cells, violins show the distribution, and the diamond represents the mean expression. P-values were calculated using a two-tailed Wilcoxon test.

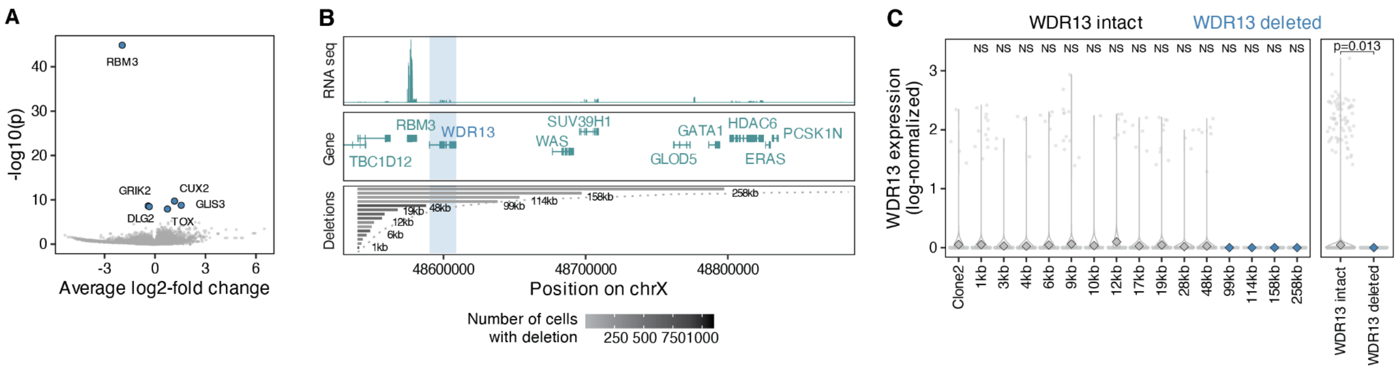

**Figure S18. scShred-seq associates deletion genotypes to transcriptional phenotypes.** (**A**) P-values (y-axis) and log2-fold changes (x-axis) from genome-wide differential expression testing (two-tailed Wilcoxon test) between cells with deletions from Figure 5H that overlap or spare *RBM3*. Points represent individual genes. The six most significantly differentially expressed genes are highlighted in blue. (**B**) Deletions and genomic features (panels) at a region on chromosome X bearing an integrated beacon (x-axis). From top to bottom: (i) RNA-seq coverage in HAP1 cells. (ii) Exon structure and names of genes in region. (iii) Deletions observed in the scShred-seq experiment, with each line corresponding to the length of one deletion and shaded according to the number of cells with deletion evidence. Genomic locations overlapping the *WDR13* gene are shaded. (**C**) *WDR13* expression (y-axis) for cells based on deletion status (x-axis). Deletions overlapping *WDR13* are highlighted in blue. Markers represent individual cells, violins show the distribution, and the diamond represents the mean expression. P-values were calculated using a two-tailed Wilcoxon test and adjusted for multiple hypothesis testing (Benjamini-Hochberg) comparing cells with deletions to cells from clone 2. The right panel shows a comparison between cells with deletions that overlap or spare *WDR13*.
