## Supplementary Note 1 for "Gigabase-scale deletion scanning of the human genome"

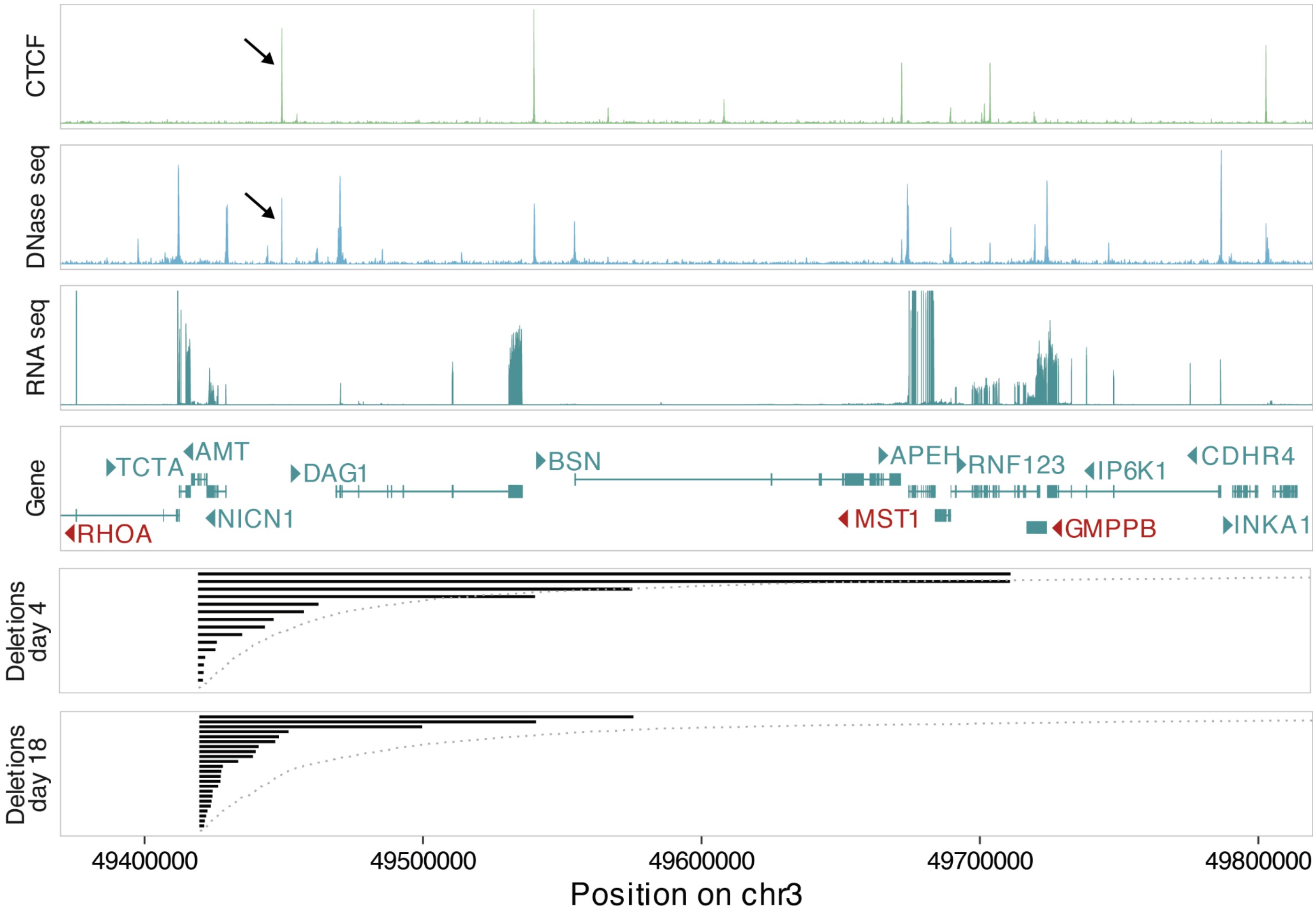


**Figure SN1. Example of a beacon integration site with a constrained deletion profile, but no established essential gene downstream.** Genomic features and deletions (panels) at a region on chromosome 3 bearing an integrated beacon. From top to bottom: (i) CTCF-Chip-Seq coverage track in HAP1 cells. (ii) DNase-seq coverage track in HAP1 cells. (iii) RNA-seq coverage in HAP1 cells. (iv) Exon structure and names of genes in region. Essential genes are marked in Red. Arrows indicate direction of transcription. (v) Deletions observed on day 4, with each line corresponding to length of one deletion. (vi) Deletions observed on day 18. Arrows indicate the location of an accessible CTCF site.
